# Optical tools for visualizing and controlling human GLP-1 receptor activation with high spatiotemporal resolution

**DOI:** 10.1101/2023.02.14.528498

**Authors:** Loïc Duffet, Elyse T. Williams, Andrea Gresch, Simin Chen, Musadiq A. Bhat, Dietmar Benke, Nina Hartrampf, Tommaso Patriarchi

**Author notes:** co-first authors (Loïc Duffet, Elyse T. Williams).

## Abstract

The glucagon-like peptide-1 receptor (GLP1R) is a broadly expressed target of peptide hormones with essential roles in energy and glucose homeostasis, as well as of the blockbuster weight-loss drugs semaglutide and liraglutide. Despite its large clinical relevance, tools to investigate the precise activation dynamics of this receptor with high spatiotemporal resolution are limited. Here we introduce a novel genetically-encoded sensor based on the engineering of a circularly-permuted green fluorescent protein into the human GLP1R, named GLPLight1. We demonstrate that fluorescence signal from GLPLight1 accurately reports the expected receptor conformational activation in response to pharmacological ligands with high sensitivity (max ΔF/F_0_ = 528%) and temporal resolution (τ_ON_ = 4.7 sec). We further demonstrated that GLPLight1 shows comparable responses to GLP-1 derivatives as observed for the native receptor. Using GLPLight1, we established an all-optical assay to characterize a novel photocaged GLP-1 derivative (photo-GLP1) and to demonstrate optical control of GLP1R activation. Thus, the new all-optical toolkit introduced here enhances our ability to study GLP1R activation with high spatiotemporal resolution.

## Introduction

The glucagon-like peptide-1 receptor (GLP1R) is expressed in various parts of the brain, especially in the basolateral amygdala and hypothalamic regions (Alvarez *et al*., 2005; Cork *et al*., 2015; Trapp and Brierley, 2022, p. 1; Turton *et al*., 1996, p. 1), as well as broadly outside the central nervous system (Campos *et al*., 1994). Its endogenous ligand, glucagon-like peptide-1 (GLP-1), is a peptide, fully conserved across mammals, that carries out both central and endocrine hormonal functions for the control of energy homeostasis (Andersen *et al*., 2018, p. 1). GLP-1 is produced mainly by two cell types: preproglucagon (PPG) neurons principally located in the Nucleus of the Solitary Tract (NTS) of the brain (Trapp and Brierley, 2022; Turton *et al*., 1996), and enterocrine cells (ECs) located in the gut (Trapp and Brierley, 2022). Upon ingestion of a meal, GLP-1 is rapidly released along with gastric inhibitory polypeptide (GIP) from the gut into the bloodstream where it targets β-cells of the pancreas and stimulates the production and secretion of insulin under hyperglycemic conditions (Andersen *et al*., 2018, p. 1). This phenomenon, known as the “incretin effect” (Nauck and Meier, 2018), is impaired in metabolic disorders, such as type-2 diabetes mellitus (Holst *et al*., 2009), making GLP-1 signaling an attractive therapeutic target for the treatment of these disorders. In addition to its role in controlling satiety and food intake, central GLP-1 has also been shown to play central neuroprotective roles (Hölscher, 2022), illustrating its multifaceted role in human physiology.

The human GLP1R (hmGLP1R) is a prime target for drug screening and drug development efforts, since GLP-1 receptor agonists (GLP1RAs) have been used for decades for the treatment of type-2 diabetes and have more recently become some of the most effective and widely-used weight-loss drugs (Shah and Vella, 2014). Among the techniques that can be adopted in these screening efforts are those able to monitor ligand binding to GLP1R through radioactivity-based assays (Knudsen *et al*., 2007) or fluorescently-labelled ligands (Ast *et al*., 2020), and those able to monitor the coupling of GLP1R to downstream signaling pathways, for example through scintillation (Runge *et al*., 2003), fluorescence (Biggs *et al*., 2018) or bioluminescence resonance energy transfer assays (Zhang *et al*., 2020, p. 1). A technology to directly probe ligand-induced GLP1R conformational activation with high sensitivity, molecular specificity and spatiotemporal resolution could facilitate drug screening efforts and open important new applications (Chen *et al*., 2022; Frank *et al*., 2018), but is currently lacking.

To overcome these limitations, here we set out to engineer and characterize a new genetically-encoded sensor based on the GLP1R, using an established protein engineering strategy (Duffet *et al*., 2022a; Patriarchi *et al*., 2018, 2019; Sun *et al*., 2018). This sensor, which we call GLPLight1, offers a direct and real-time optical readout of GLP1R conformational activation in cells, thus opening unprecedented opportunities to investigate GLP1R physiological and pharmacological regulation in detail under a variety of conditions and systems. We demonstrated its potential for use in pharmaceutical screening assays targeting GLP1R, by confirming that GLP1R and GLPLight1 show similar ligand recognition profiles, including high specificity towards GLP-1 over other class-B1 GPCR ligands, low-affinity for glucagon, and specific functional deficits of GLP-1 alanine mutants. Finally, to extend the optical toolkit further, we also developed a photocaged GLP-1 derivative (photo-GLP1) and adopted it in concert with GLPLight1 to enable all optical control and visualization of GLP1R activation.

## Results

### Development of a genetically-encoded sensor to monitor hmGLP1R activation

To develop a genetically encoded sensor based on the hmGLP1R, we initially replaced the third intracellular loop (ICL3) of hmGLP1R with a cpGFP module from the dopamine sensor dLight1.3b (Patriarchi *et al*., 2018), between residues K336 and T343 (**Figure 1a**). This initial sensor prototype had poor surface expression and a very small fluorescence response upon addition of a saturating concentration (10 µM) of GLP-1 (ΔF/F_0_ = 39%, **Figure 1-figure supplement 1a**). Removal of the endogenous GLP1R N-terminal secretory sequence (amino acids 1-23) from this construct improved the membrane expression and the fluorescence response to GLP-1 (ΔF/F_0_ = 107%, **Figure 1-figure supplement Fig. 1a**). We then performed a lysine scan on the residues spanning the intracellular loop-2 (ICL2) of the sensor. From this screening we identified one beneficial mutation (L260K) that more than doubled the dynamic range of the sensor (ΔF/F_0_ = 180%, **Figure 1-figure supplement 1b**). Next, we performed site-saturated mutagenesis on both receptor residues adjacent to the cpGFP and screened a subset of 95 variants. This small-scale screening led us to identification of a new variant (containing the mutations K336Y and T343N) with ΔF/F_0_ of about 341% (**Figure 1-figure supplement 1c–d**). To further enhance surface expression of the sensor, we introduced a C-terminal endoplasmic reticulum export sequence (Stockklausner and Klocker, 2003) on this variant (**Figure 1-figure supplement 1e**). We then introduced three previously-described (Wan *et al*., 2021) mutations in the cpGFP moiety, which improved the basal brightness of the probe without affecting its dynamic range (**Figure 1-figure supplement 1f–g**). Finally, we mutated eight phosphorylation sites on the C-terminal domain that are responsible for GLP1R internalization (Thompson and Kanamarlapudi, 2015) (S431A, S432A, T440A, S441A, S442A, S444A, S445A, and T448A) aiming to maximally reduce the possibility of sensor internalization. The resulting sensor variant showed good membrane expression and a 528% maximal fluorescence response upon GLP-1 binding (**Figure 1b–c, Figure 1-figure supplement 1h**). This final variant was named GLPLight1 and was chosen for further characterization. To aid with control experiments during GLPLight1 validation, we also set out to develop a sensor variant carrying mutations in the peptide binding pocket. We screened a panel of 14 single point mutations and identified a combination of 3 mutations (L141A, N300A and E387A) that abolished the fluorescent response to GLP-1 application while showing a good membrane expression of the sensor (**Figure 1-figure supplement 2a–c**). This control variant was named GLPLight-ctr.

**Figure 1:**
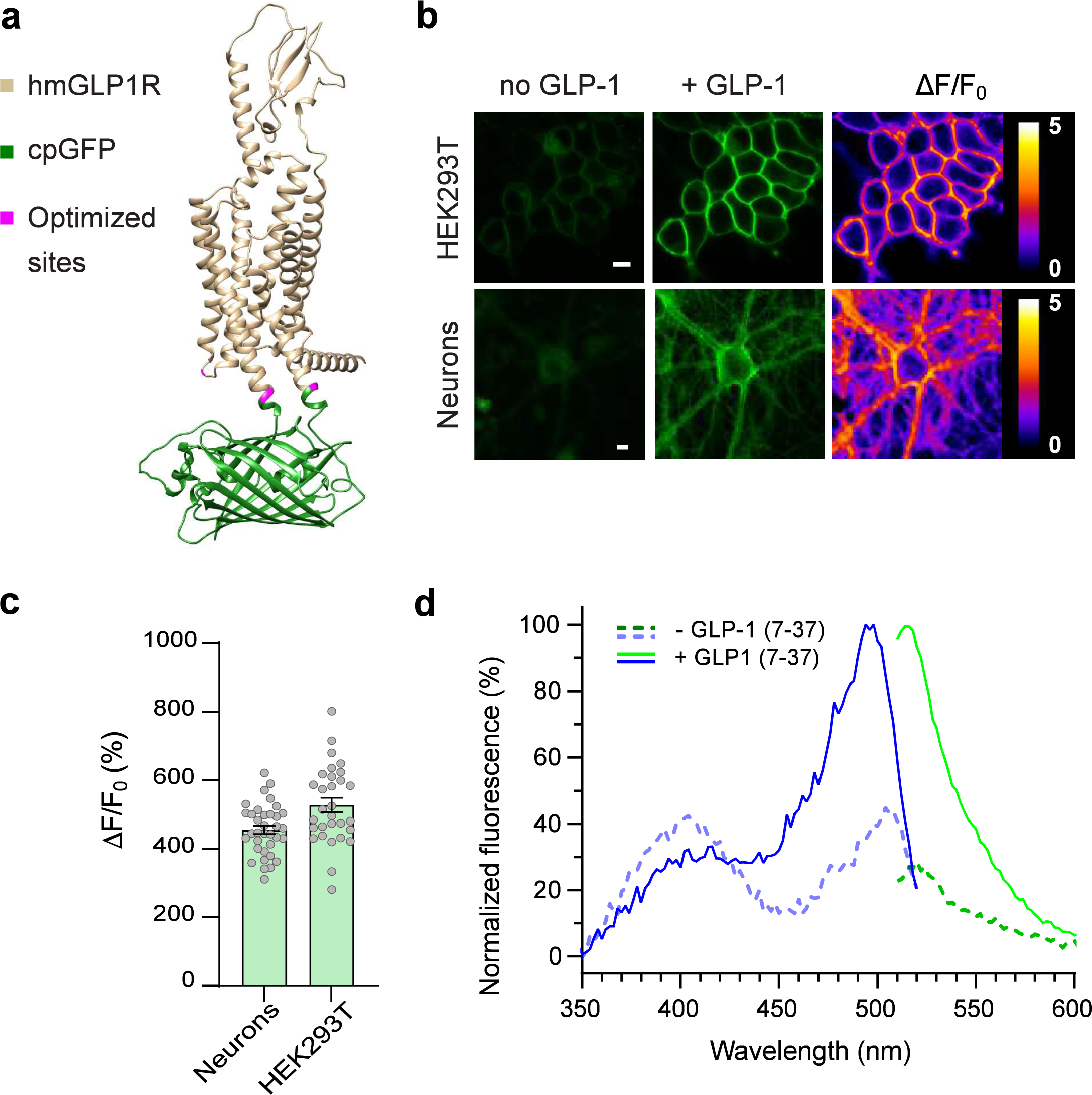
Development and optical properties of GLPLight1. **a**) Structural model of GLPLight1 obtained using Alphafold (Mirdita *et al*., 2022). The human GLP1R is shown in gold, cpGFP in green and residue targets of mutagenesis are shown in magenta. **b**) Representative images showing GLPLight1 expression and fluorescence intensity change before (left) and after (center) addition of 10 μM GLP-1(7**–**37) as well as their respective pixel-wise ΔF/F_0_ images in HEK293T cells (top) and primary cortical neurons (bottom). Scale bars, 10 μm. **c)** Maximal fluorescence response of GLPLight1 expressed in the indicated cell types after the addition of 10 μM GLP1. n = 35 neurons and n = 30 HEK293T cells, from 3 independent experiments. **d)** One-photon excitation/emission spectra of GLPLight1-expressing HEK293T cells before (dark green and dark blue) and after (light green and light blue) addition of 10 μM GLP-1(7**–**37) normalized to the peak excitation and emission of the GLP-1-bound state of the sensor. Data were obtained from 3 independent experiments. Only mean values are shown. All data shown as mean ± SEM unless stated otherwise.

### In vitro characterization of GLPLight1

To establish the utility of GLPLight1 as a new tool to investigate the human GLP1R in pharmacological assays, we first characterized its properties in vitro. We started by comparing sensor expression and fluorescent response among different cell types. To do so, we expressed GLPLight1 in primary cortical neurons in culture, via adeno-associated virus (AAV) transduction. Two weeks post-transduction, GLPLight1 was well expressed on the neuronal membrane and showed a maximal response of 456% to GLP-1 application (10 µM) (**Figure 1b–c**). We then measured the spectral properties of the sensor in HEK cells. The fluorescence spectra were similar to those of previously described green GPCR-sensors (Duffet *et al*., 2022a; Sun *et al*., 2018), and showed a peak excitation around 500 nm, peak emission around 512 nm, and an isosbestic point at around 425 nm (**Figure 1d**). Work on previously developed GPCR-based sensors that respond to neuropeptide ligands (Duffet *et al*., 2022a; Ino *et al*., 2022) revealed that the conformational activation kinetics of these receptor types is at least an order of magnitude slower than what has been reported for monoamine receptors (Feng *et al*., 2019; Patriarchi *et al*., 2018; Sun *et al*., 2018; Wan *et al*., 2021), likely reflecting the more complex and polytopic binding mode of peptide ligands to their receptor.

Next, we compared the coupling of GLPLight1 and its parent receptor (WT GLP1R) to downstream signaling. We first measured the agonist-induced membrane recruitment of cytosolic mini-G proteins and β-arrestin-2 using a split nanoluciferase complementation assay (Dixon *et al*., 2016). In this assay, both the sensor/receptor and the mini-G proteins contains part of a functional luciferase (smBit on the sensor/receptor and LgBit for Mini-G proteins) that becomes active only when these two partners are in close proximity (Wan *et al*., 2018). In agreement with the known pleiotropic signaling of WT GLP1R (Rowlands *et al*., 2018), in our assay activation of the receptor led to a strong recruitment of miniGs, miniGq, miniGi, β-arrestin-2 as well as miniG12, albeit to a lower extent. In comparison to WT GLP1R, the coupling of GLPLight1 to all tested signaling partners was significantly reduced **(Figure 1-figure supplement 3a-j**). To further confirm the absence of coupling to intracellular cyclic-AMP (cAMP) signaling of GLPLight1, we performed a titration of GLP-1 on the sensor and WT GLP1R in a luminescence-based cAMP assay. This revealed that the WT GLP1R showed could potently elicit intracellular cAMP increases with an EC_50_ of 8.0 pM whereas no such increase was observed for GLPLight1 even at the highest GLP-1 concentrations tested (100 nM, **Figure 1-figure supplement 3k**). We also performed a titration of GLP-1 induced recruitment of miniGs protein where we could show that GLP1R effectively recruits miniGs proteins with an EC_50_ of 3.8 nM (**Figure 1-figure supplement 3l**). These results indicate that GLPLight1 is unlikely to couple with endogenous intracellular signaling pathways.

### Application of GLPLight1 as a tool for pharmacological screening

GLPLight1 is a novel genetically-encoded sensor capable of providing a sensitive intensiometric readout of hmGLP1R activation in response to its endogenous ligands. As such, this tool could have great potential for applications in the drug discovery and development field; however, a more careful characterization of its pharmacological response profile is needed to ensure its implementation as a screening tool. We thus performed a series of in vitro pharmacological experiments in which we characterized GLPLight1 responses under different conditions and with a variety of ligands with known pharmacological effects on GLP1R, with the aim to demonstrate the applicability of this sensor as a pharmacological screening tool. We started by testing the reversibility of sensor response via competition of GLP-1 with an antagonist peptide. To do so, we imaged GLPLight1-expressing HEK293T cells upon addition, in sequence, of 1.0 µM GLP-1 followed by 10 µM Exendin-9 (Ex-9), a well-known peptide antagonist of GLP1R. Ex-9 could partially reverse the signal to 42% of the maximal GLP-1 response, within less than 5 minutes in vitro (**Figure 2a–b**). Next, we tested whether two clinically-used anti-obesity drugs that are known GLP1RAs, liraglutide or semaglutide (O’Neil *et al*., 2018), could trigger a response from the sensor. As expected, GLPLight1 responded to both GLP1RAs with almost maximal activation, on par with GLP1 (**Figure 2a**). These results indicate that GLPLight1 can serve as a direct readout of pharmacological drug action on the hmGLP1R with higher temporal resolution than previously available approaches, such as downstream signaling assays (Zhang *et al*., 2020).

**Figure 2:**
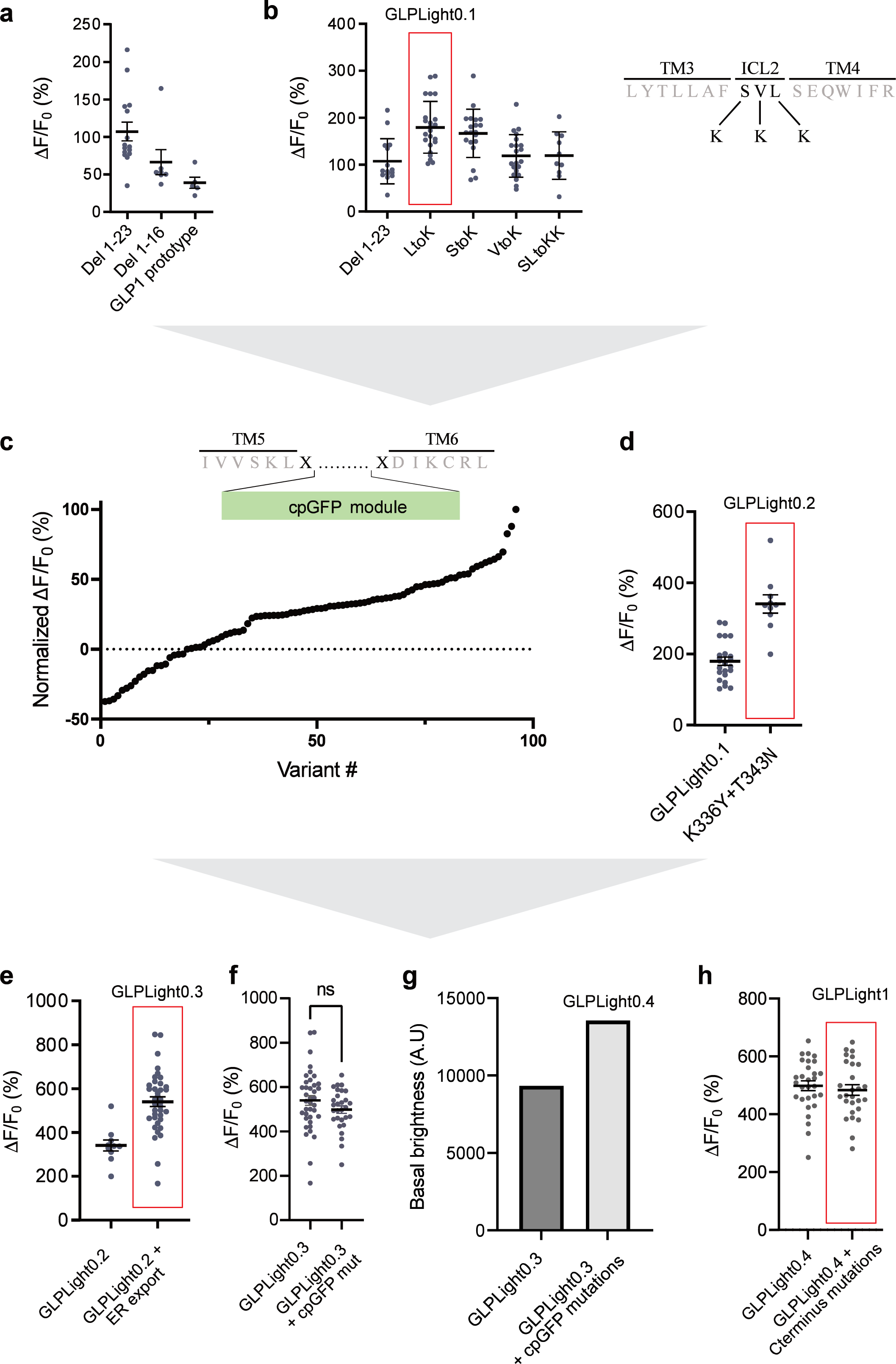
Pharmacological characterization of GLPLight1. **a**) Absolute ΔF/F_0_ responses of GLPLight1 expressing HEK293T cells to various GLP-1-agonists, antagonist, or to other class-B1 neuropeptide ligands applied at 1 μM final (unless stated otherwise). n = 30 cells from three independent experiments for all conditions. **b**) ΔF/F_0_ responses from time-lapse imaging experiments in which 1 μM GLP-1 and 10 μM exendin-9 (Ex-9, a peptide antagonist of GLP1R) were subsequently bath-applied onto GLPLight1-expressing HEK293T cells. n = 30 cells from three independent experiments. **c**) Normalized dose-response curves showing the fluorescent responses of GLPLight1-expressing HEK293T cells and primary cortical neurons to GLP-1 (dark green and light green respectively) or GLPLight1-expressing HEK293T cells to glucagon (purple). The curves fit were performed using a 4-parameter equation and the mean EC_50_ values determined are shown next to the traces in the corresponding color. n=3,6 and 3 independent experiments for GLP-1(7**–**37) in neurons, GLP-1(7**–**37) and glucagon in HEK293T cells, respectively. **d**) Dose-response curves showing the fluorescent responses of GLPLight1-expressing HEK293T cells to alanine mutants of the GLP-1 peptide normalized to the maximum mean fluorescence response (FITC intensity) obtained for the WT GLP-1 peptide. n = 3 independent experiments for each peptide. All data are displayed as mean ± SEM.

Knowing that GLP-1 is produced along with GLP-2 and glucagon via proteolytic processing of a common preproglucagon precursor protein (Drucker, 2001), we decided to investigate the specificity of our sensor against these other peptides. While the sensor did not respond with any detectable increase in fluorescence to GLP-2, it responded to glucagon with a ΔF/F_0_ of 324% (61% of maximal response to GLP-1). To further characterize the sensitivity of GLPLight1 to its two endogenous agonists, we performed titrations of GLP-1 and glucagon in HEK293T cells and determined that GLPLight1 had a 94-fold higher affinity for GLP-1 compared to glucagon (EC_50_ = 28 nM for GLP-1, EC_50_ = 2.6 µM for glucagon), in agreement with previously-reported results employing a downstream cAMP readout (Runge *et al*., 2003). Furthermore, the affinity of GLP-1 measured in primary neurons (EC_50_ = 9.3 nM) was comparable to the one in HEK cells (**Figure 2c**). Additionally, GLPLight1 did not respond to a panel of other endogenous class B1 GPCR peptide ligands that were tested at high concentration (1.0 µM), including GIP, CRF, PTH, PACAP or VIP.

The binding of GLP-1 to its receptor occurs via the N-terminus of the peptide, as demonstrated by previous structural (Jazayeri *et al*., 2017) and mutagenesis studies (Longwell *et al*., 2021; Zhang *et al*., 2020). We therefore set out to determine whether the general trends observed by fluorescence response of GLPLight1 is in agreement with the pharmacological readout of GLP1R activation obtained using classical assays (Adelhorst *et al*., 1994). We synthesized four single-residue alanine mutants of GLP-1 at selected N-terminal positions (H1A, E3A, G4A, T5A) using automated fast-flow peptide synthesis (AFPS, see Supplementary Information) (Hartrampf *et al*., 2020; Mijalis *et al*., 2017). All peptides were obtained in good yields and excellent purities after RP-HPLC purification. Titrations of individual GLP-1 mutants on GLPLight1-expressing cells revealed clear effects of the mutations on either the maximal sensor response (E_max_), the potency (EC_50_) of the peptide ligand, or both (**Table 1** and **Figure 2d**). In particular, the critical role of H1 and G4 for both binding and activation has been reported in the literature several times (Manandhar and Ahn, 2015). In agreement with these results, we observed a significant reduction of E_max_ and EC_50_ for H1A (56% and 1300 nM, respectively) (**Table 1**, **Entry a**) and G4A (14% and 993 nM, respectively) (**Table 1**, **Entry c**), compared to WT GLP-1 using GLP1Light as a readout. Furthermore, position E3 was reported to be critical for binding, but not for activation. Here, we determined an E_max_ of 96% compared to WT GLP-1 (**Table 1**, **Entry b**), as well as a reduced EC_50_ (757 nM) for E3A, which is in agreement with the literature (Manandhar and Ahn, 2015) (**Table 1**). Finally, T5 has been reported as less important for GLP1R binding and activation than the other investigated peptide positions (Adelhorst *et al*., 1994). Accordingly, our experiments with GLPLight1 T5A showed the highest E_max_ (100%) and EC_50_ (188 nM) (**Table 1**, **Entry d**) of all alanine mutants investigated herein. Overall, we conclude that fluorescence response of GLPLight1 can be used to study the relative trends for E_max_ and potency of GLP1R ligands.

**Table 1:** Titration parameters of alanine scanned variants of GLP-1 peptide. The E_max_ and pEC_50_ values were derived from the four-parameter non-linear fit for each peptide and the EC_50_ shift by comparison against WT GLP-1 peptide measured alongside.

| Entry | GLP-1 Variant | $E_{\max}$ (% WT GLP-1) | $pEC_{50}$ (M) | Fold-reduction $EC_{50}$ vs. WT GLP-1 |
| --- | --- | --- | --- | --- |
| a | H1A | $56 \pm 4$ | $5.89 \pm 0.05$ | $\approx 63$ |
| b | E3A | $96 \pm 3$ | $6.10 \pm 0.03$ | $\approx 37$ |
| c | G4A | $14 \pm 3$ | $6.00 \pm 0.13$ | $\approx 48$ |
| d | T5A | $100 \pm 4$ | $6.73 \pm 0.04$ | $\approx 9$ |
| n/a | WT GLP-1 | 100 | $7.69 \pm 0.04$ | 0 |

State-of-the-art techniques for detecting endogenous GLP-1 or glucagon release in vitro from cultured cells or tissues consist of costly and time-consuming antibody-based assays (Kuhre *et al*., 2016) or analytical chemistry procedures (Amao *et al*., 2015). Given the genetically-encoded nature and the fast optical readout of GLPLight1, this tool has the potential to facilitate studies investigating the physiological regulation of GLP-1 release in vitro. To establish whether GLPLight1 could be sensitive enough to detect endogenous GLP-1 release in an in vitro setting, we cultured sensor-expressing HEK293T cells in the presence or absence of a GLP-1/glucagon-producing immortalized enteroendocrine cell line (GLUTag cells (Brubaker *et al*., 1998)). To distinguish the two cell types in the co-culture system, the HEK239T cells were co-transfected with a cytosolic red fluorescent protein (mKate2). To detect whether the GLPLight-expressing cells had detected endogenous GLP-1 release by the enterocrine cells, we bath-applied GLP-1 to cause full activation of the sensor. We observed that the response to GLP-1 of sensor-expressing cells cultured in the presence of GLUTag cells was significantly lower than that of cells cultured in their absence (**Figure 2-figure supplement 1**). These results indicate that GLPLight1 was partially pre-activated by endogenous GLP-1 secreted by the enterocrine cells present in the dish. The detection of endogenous GLP-1 by the sensor opens the possibility to use it as a screening tool for studying intrinsic/extrinsic factors that regulate GLP-1 release from enterocrine cells in vitro.

### Development and in vitro characterization of photo-GLP1

To investigate the spatiotemporal activation of GLP1R and GLPLight1, a photocaged derivative of GLP-1 was envisioned. To ensure that the photocaged GLP-1 derivative does not activate GLP1R or GLPLight1 prior to uncaging (i.e. in the dark), the photocage must be located on or near GLP-1 regions that are essential for binding. Photocaging of peptides can be achieved by the attachment of a photocaging molecule at a side-chain functionality, backbone amide, or at the C- or N-terminus of the peptide. Recently, we reported the optical control of orexin-B using a UV-visible light-sensitive C-terminal photocage (Duffet *et al*., 2022b). As opposed to orexin-B, GLP-1 primarily binds via its N-terminus to GLP1R (Jazayeri *et al*., 2017). We therefore explored an N-terminal caging strategy to generate a photocaged GLP-1 derivative (photo-GLP1, Figure 3a). GLP-1 was prepared by solid-phase peptide synthesis (SPPS) utilizing AFPS (Hartrampf *et al*., 2020; Mijalis *et al*., 2017). Before cleavage of the peptide from the resin, photocaging of the GLP-1 N-terminal amine was carried out by treating the resin-bound peptide with an active ester (*N*-hydroxysuccinimide [NHS] ester) form of the nitrobenzene-type photocage (see **Supplementary Data**). Cleavage of the resulting photocaged peptide from the resin followed by RP-HPLC purification successfully provided photo-GLP1 in 5% overall yield with >95% purity. To confirm the release of WT GLP-1 upon treatment of photo-GLP1 with UV light, photo-GLP1 (80 µM in HBSS) was irradiated under LED light (λ = 370 nm, 0.64 mW/mm^2^) for 20 min with air cooling. Subsequent LCMS and UHPLC analysis demonstrated complete uncaging of photo-GLP1 to afford WT GLP-1, confirmed by co-injection of a standard sample of WT GLP-1 (80 µM in HBSS) (**Figure 3-figure supplement 1**).

**Figure 3:**
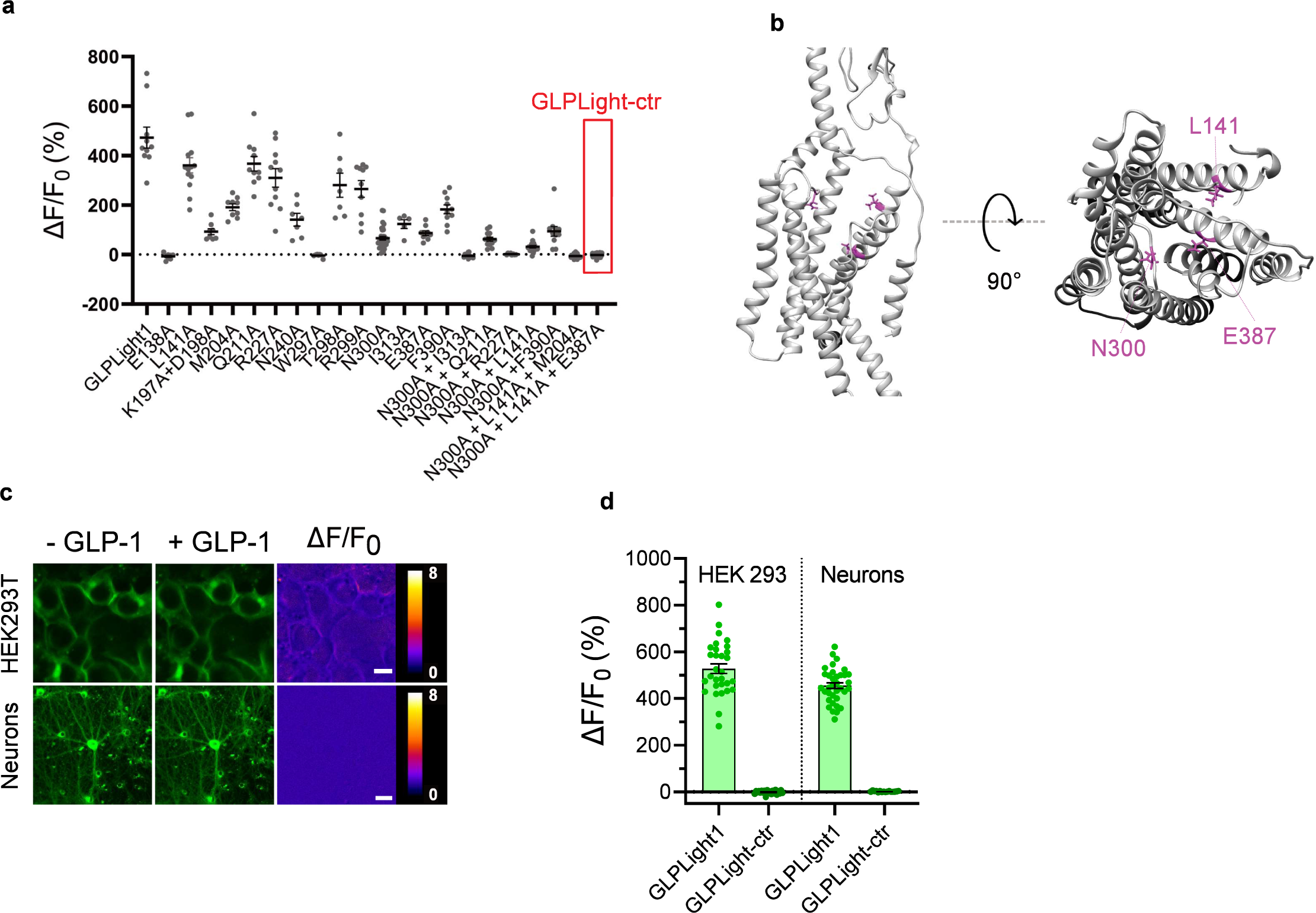
All-optical visualization and control of human GLP1R activation. **a**) Schematic representation of the N-terminal chemical caging strategy used to generate photo-GLP1. The peptide product (native GLP-1) after optical uncaging is shown. **b**) Time-lapses of fluorescence response for GLPLight1 or GLPLight-ctr expressing HEK293T cells before and after optical uncaging (purple vertical bar, 405 nm laser, scanning rate of 0.8 Hz and variable durations as specified below the graph). The values were normalized to the maximal response of GLPLight1 between t = 150 and 200 sec to 10 μM GLP-1 (purple trace). In all cases, cells were pre-incubated for 1**–**2 minutes with 10 μM photo-GLP1 before imaging and optical uncaging started. All fluorescent signals were analyzed within 20 μm distance from the uncaging area. n=11 to 19 cells from three independent experiments. **c)** Quantification of the normalized average fluorescence from **b** between t = 25 to 75 sec for uncaging experiments, and t = 150 to 200 sec for GLP-1 application experiments. All uncaging experiments on GLPLight1 were compared to the one with pre-incubation with Exendin-9 (Ex-9, see Figure 3-figure supplement 2) using Brown-Forsythe ANOVA test followed by Dunnett’s T3 multiple comparison. p = 0.0061; 0.0001; 0.0026 and 1,293 x 10^-6^ for 10, 20, 50 and 100 sec uncaging events, respectively. **d)** Fluorescence response of GLPLight1-expressing HEL293T cells to 10 μM GLP-1 either after pre-incubation with 10 μM photo-GLP1 (magenta) or in the absence of it (blue). The data were normalized to the maximal response of GLPLight1-expressing cells in the absence of photo-GLP1 and fitted with a non-linear mono exponential fit to determine τ values. Statistical analysis was performed using the extra sum-of-squares F test; ****p< 0.0001; n=18 and 17 cells from three independent experiments in the absence or presence of photo-GLP1 respectively. All data are displayed as mean ± SEM. **e**) Kymographs of representative cells from **d** showing the fluorescence intensity of a line drawn across a cell membrane over time in the absence (top) or presence (bottom) of photo-GLP1 (10 μM) in the bath. The timepoint of GLP-1 application is shown by the red dotted line. **f**) Representative images of multiple uncaging events performed at different locations across the field of view. Images show the basal fluorescence of GLPLight1-expressing HEK cells (left) in the presence (top) or absence (bottom) of 10 μM photo-GLP1, as well as the corresponding pixel-wise heatmap of SNR post-uncaging. Localized functional sensor responses to optical uncaging of photo-GLP1 are indicated by white arrows. Uncaging was performed for a duration of 40 sec in total for all the three areas shown as white squares using a scanning rate of 1.5Hz. Scale bars: 20 μm. **g**) Quantification of the timelapse of fluorescence response of GLPLight1 from **f** inside the uncaging areas. **h**) Same as **f** but with a sub-cellular localized uncaging region selected on the membrane of a GLPLight1-expressing cell with 1,5 s uncaging duration and a 25 Hz scanning rate. Scale bar 10 μm.

We then leveraged on GLPLight1 to establish an all-optical assay for characterizing photo-GLP1 uncaging in vitro. We bath-applied photo-GLP1 (10 µM) onto GLPLight1-expressing HEK293T cells and performed optical uncaging by exposing a defined area directly next to the cells to 405 nm laser light (UV light) for defined periods of time, while the sensor fluorescence was imaged using 488 nm laser light. Application of photo-GLP1 by itself failed to trigger any response from GLPLight1, indicating a lack of functional activity in the absence of UV light (**Figure3-figure supplement 2a**). On the contrary, after photo-GLP1 was added to the bath, the fluorescence of GLPLight1 visibly increased upon 10 seconds of UV light exposure, indicating that GLP-1 could successfully be uncaged and activated the sensor on the cells. Higher durations of UV light exposure led to a higher degree of GLPLight1 responses, and the maximal uncaging duration tested (100 sec) triggered approximately 30% of the maximal response of the sensor, as assessed in the same assay by bath-application of a saturating GLP-1 concentration (10 µM) (**Figure 3b–c**). Importantly, to show that the sensor signals are not due to UV light-induced artifacts, we reproduced the maximal (100 sec) uncaging protocol on GLPLight-ctr-expressing HEK293T cells and confirmed that in this case no sensor response could be observed. Furthermore, pre-treatment of the cells with the GLP1R antagonist Ex-9 significantly blunted the sensor response evoked by the optical uncaging (100 sec) (**Figure 3c, Figure3-figure supplement 2b**). These results indicate that photo-GLP1 can be effectively uncaged in vitro using 405 nm light to control hmGLP1R activation.

### High-resolution all-optical visualization and control of GLP1R activity

Upon performing the uncaging experiments, we noticed that the profile of the sensor response to bath-applied GLP-1 differed, depending on whether or not photo-GLP1 was present in the bath. To investigate this phenomenon more in detail, we measured and compared the sensor activation kinetics when GLPLight1 was activated by direct bath application of GLP-1 in the presence or absence of an equimolar concentration of photo-GLP1 in the bath. The sensor response was strikingly different in the two conditions, and exhibited an approximate 14-fold reduction in the speed of activation in the presence of photo-GLP1 (τ_ON_ without photo-GLP1 = 4,7 sec; τ_ON_ with photo-GLP1 = 68,1 sec; **Figure 3d–e**). These results indicate that photo-GLP1, in the dark (i.e. with an intact photocage), can affect the kinetics of GLP1R activation, and this is likely mediated by its binding to the receptor extracellular domain (ECD), which competes for the functionally-active GLP-1. In fact, since the GLP1R belongs to class-B1 GPCRs, the binding of GLP-1 is known to involve an initial step where the peptide C-terminus is recruited to the ECD, followed by a second step involving insertion of the peptide N-terminus into the receptor binding pocket (Wu *et al*., 2020). Given that our photocage was placed at the very N-terminus of photo-GLP1, our results show that this caging approach prevents the peptide’s ability to activate GLPLight1 but, at the same time, preserves its ability to interact with the ECD.

We next asked whether we could leverage GLPLight1 to obtain spatial information on the extent of GLP1R activation in response to photo-GLP1 uncaging. To do so, we performed optical photo-GLP1 uncaging on three separate areas of about 400 µm^2^ placed at different locations in a large field of view (FOV, approximately 40,000 µm^2^). UV light was applied for a total of 40 seconds on the three uncaging regions during the imaging session. GLPLight1 shows a fluorescent response in all three uncaged areas, while its fluorescence remained unaltered throughout the rest of the FOV, indicating high spatial localization of the response to GLP-1 (**Figure 3f**). As a control, the omission of photo-GLP1 in the cell bath led to no sensor response upon uncaging (**Figure 3g**). Additionally, the same session was repeated on GLPLight-ctr-expressing cells. Also in this case, no response to uncaging could be observed (**Figure 3f**). To determine whether the sensor readout in this assay could report GLP1R activation with even subcellular resolution, we repeated the uncaging experiment by selecting an uncaging area of approximately 16 µm^2^ directly on a cell membrane. In this case, the application of UV light led to localized activation of GLPLight1 that was limited to a portion of the cell membrane and did not spread to neighboring cells (**Figure 3h**). These results demonstrate that the optical nature of the GLPLight1 readout makes it possible to determine the spatial extent of GLP1R activation with very spatial high-resolution, down to sub-cellular domains.

Finally, we tested whether uncaging of photo-GLP1 could be used to control functional signaling downstream of hmGLP1R activation. To this aim, we employed a recently developed genetically encoded sensor for cyclic-AMP (G-Flamp1) (Wang *et al*., 2022), which is the main second messenger involved in cellular signaling downstream of GLP1R activation (Holz *et al*., 2015). We imaged a field of HEK293T cells co-transfected with the hmGLP1R and G-Flamp1 (**Figure 4a**) during application of photo-GLP1 to the cells and after optical uncaging of photo-GLP1 (2 sec, 1 nM) within a limited area of about 70 µm^2^ located directly above a single HEK293T cell. As a result of uncaging, the signal from the cAMP sensor increased visibly and significantly only in the cell directly underneath the uncaged area (**Figure 4b**). The same uncaging protocol applied in the absence of photo-GLP1 on the cells failed to trigger any response from the cAMP sensor (**Figure 4-figure supplement 1**), indicating that the sensor signals reliably reported intracellular cAMP signaling triggered by uncaged GLP-1. Furthermore, as a positive control, we bath-applied the same concentration of GLP-1 (1 nM) at the end of each recording to stimulate simultaneously the activation of the receptor on all cells. Indeed, this could elicit a response in all the imaged cells that did not respond to the previous uncaging protocol (‘distant cells’) (**Figure 4b–d**). As part of our observations, we observed a small dip of the G-Flamp-1 signal in response to photo-GLP1 bath application. To assess whether this signal drop was caused by the signaling activity of the photo-GLP1 or was an artifact from G-Flamp-1 imaging, we repeated the measurement by applying HBSS to the cells. The small signal drop could be detected also in these experiments (**Figure 4-figure supplement 1**), demonstrating that the initial dip in G-Flamp-1 signal was artefactual, possibly due to temperature or pressure changes onto the cells. Overall, our results demonstrate that uncaging of photo-GLP1 can be used to achieve optical control of GLP1R signaling activation with high spatiotemporal resolution.

**Figure 4:**
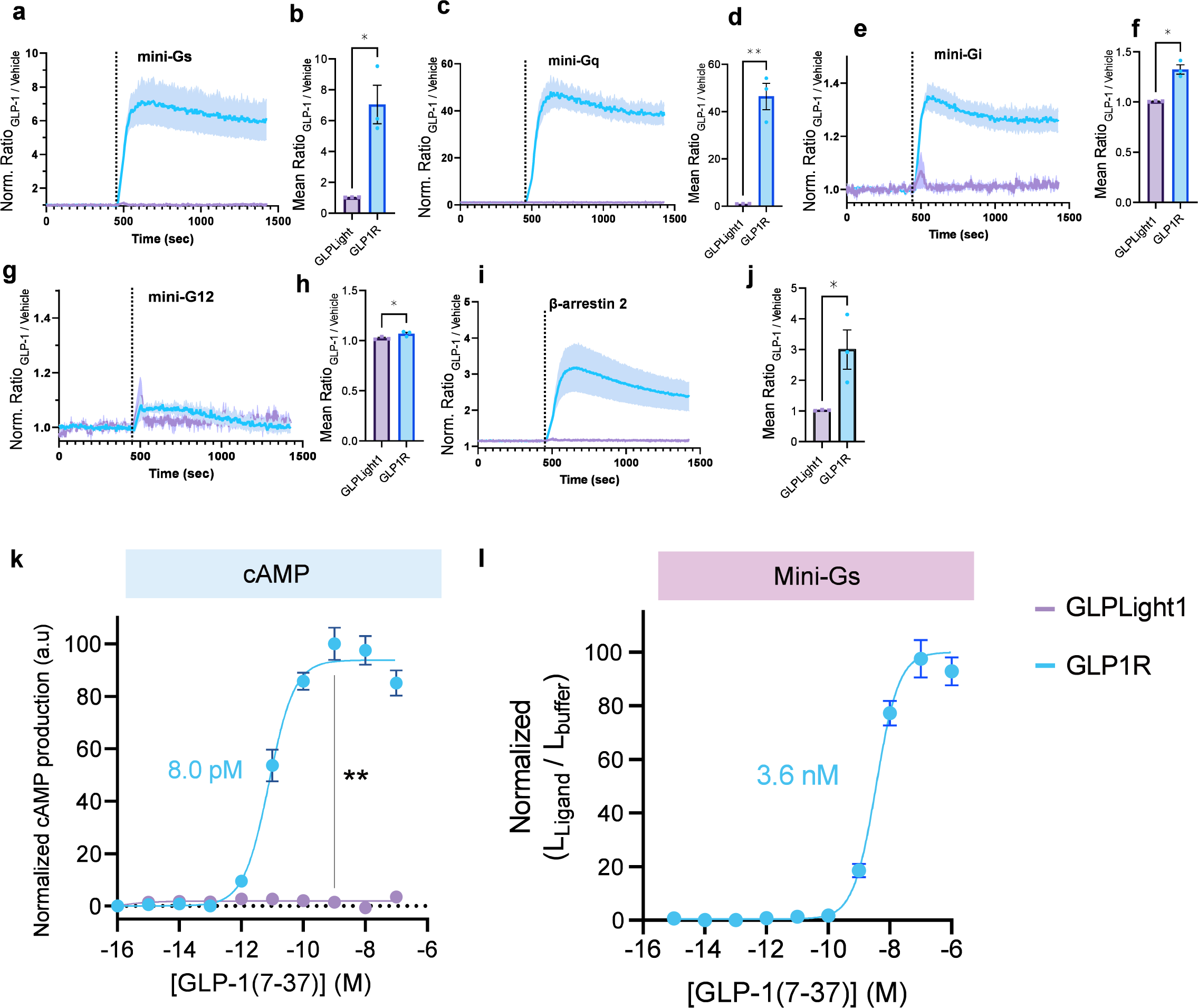
Effect of photo-GLP1 uncaging on intracellular signaling. **a)** Representative images of HEK293T cells used in **c,d**. hmGLP1R expression was visualized using an Alexa-647 conjugated anti-FLAG antibody. The uncaging region is represented by the white square in the lower right area. **b**) Representative images from the pixel-wise fluorescence response from G-Flamp1 after uncaging (left) and bath-application of 1 nM WT GLP-1 (right). The white arrow indicates the localized area of uncaging. **c**) Fluorescence change during timelapse imaging of G-Flamp1/GLP1R co-expressing cells after addition of 1 nM photo-GLP1 and localized uncaging (405 nm, 2 sec and 32 Hz scanning rate), followed by bath-application of 1 nM WT GLP-1. The timepoints of ligand addition are represented using grey rectangles and the uncaging bout by the vertical black line. Quantification of the fluorescence response is shown separately for ‘target cells’ (blue, cells within the uncaging area) and for ‘distant cells’ (grey, cells positioned at least 10 μm away from the uncaging area). The fluorescent responses from G-Flamp1 were normalized to the maximal activation after addition of WT GLP-1. **d**) Quantification from the average normalized fluorescence in **c** between t=400 and t=450 sec using the 10 frames before uncaging as a baseline for each condition. n = 3 ‘target cells’ and 15 ‘distant cells’ from 3 independent experiments. *P = 0.03061 using a one-tailed Student’s t-test with Welch’s correction. All scale bars: 20 μm and all data displayed as mean ± SEM.

## Discussion

Here, we report the first genetically encoded sensor engineered based on cpGFP and the human GLP1R. We show that this tool can directly report ligand-induced conformational activation of this receptor with the high sensitivity and spatiotemporal resolution typical of GPCR-based sensors. Using this new probe, we found that ligand-induced conformational activation of the human GLP1R occurs on slower timescales compared to the reported kinetics of other similarly-built GPCR-sensors (Labouesse and Patriarchi, 2021). This new insight is not surprising given that previously developed sensors were built from class-A GPCRs (Labouesse and Patriarchi, 2021), while GLP1R belongs to a different class of GPCRs (class B1) that is characterized by a distinct ligand-binding mechanism that involved initial ligand ‘capture’ by the receptor’s ECD, followed by ligand insertion into the receptor binding pocket for initiating the transduction of signaling (Zhang *et al*., 2020). As a reference, other previously-characterized class-A GPCR-based neuropeptide biosensors showed sub-second activation kinetics (Duffet *et al*., 2022a; Ino *et al*., 2022). Accordingly, our observations show that the receptor activation kinetics can be largely influenced by pre-incubation with an inactive form of the GLP-1 peptide (photo-GLP1), likely because the inactive peptide interacts with and occupies the receptor’s ECD.

We showcased the sensitivity and utility of GLPLight1 as a pharmacological tool to aid drug screening and development efforts by characterizing its response to various naturally occurring peptide ligands, as well as clinically-used agonists and peptide-derivatives with diverse pharmacological actions on GLP1R. Besides its applications in pharmacology and drug discovery, given the high sensitivity and lack of interference with intracellular signaling of GLPLight1 it might be possible to employ this tool to investigate the dynamics of endogenous GLP-1 and/or glucagon directly in living systems (in vivo), although based on the evidence provided in this study the in vivo utilization of the sensor is not guaranteed to succeed.

The apparent EC_50_ of GLPLight1 fluorescence response to GLP-1 is very similar to that measured for mini-Gs recruitment to the hmGLP1R, while it is approximately three orders of magnitude lower than that of the cAMP response downstream of hmGLP1R. This discrepancy might be due to intrinsic differences of the assays used or to intrinsic differences in the distinct aspects of the signaling pathway investigated (i.e. direct recruitment of mini-Gs versus enzymatically-amplified cAMP signals). This raises the interesting possibility that under physiological conditions GLP-1 might elicit different functional responses based on the location of its action and on the spatial concentration gradients on target cells/tissues.

Given that GLPLight1 produces a fluorescence readout that is more representative in terms of sensitivity to that measured by direct recruitment of mini-Gs proteins to the hmGLP1R, the characteristics of this sensor appear not suitable to detect the concentration range achieved by GLP-1 in the periphery through endocrine signaling (picomolar levels). Nevertheless, it is conceivable that under specific circumstances, for example in specific brain areas or in close proximity to enteroendocrine cells in the gut, levels of GLP-1 release might reach high-enough levels that could be detected by GLPLight1. Future studies could attempt in vivo use of the sensor to further explore this interesting direction, for example by leveraging on AAV-mediated expression of GLPLight1 in living tissues or animals for implementing its use through in vivo imaging techniques, such as fiber photometry (Gunaydin *et al*., 2014), mesoscopy (Cardin *et al*., 2020) or two-photon microscopy (Helmchen, 2009). Through such efforts, GLPLight1 might be helpful to shine new light on the hidden mechanisms of GLP-1 and/or glucagon release dynamics in relation to physiological or pathological conditions.

Finally, we leveraged GLPLight1 to characterize the uncaging of the photocaged GLP-1 derivative (photo-GLP1) described for the first time in this work. Optical tools to selectively activate GLP1R could contribute to mechanistic studies (Chen *et al*., 2022; Frank *et al*., 2018), and the photoswitchable GLP-1 LirAzo was recently used to optically control insulin secretion and cell survival (Broichhagen *et al*., 2015). As opposed to photoswitchable peptides, in which the side chain or part of the peptide backbone is replaced by a photoswitchable moiety such as an azobenzene, photo-GLP1 releases native GLP-1 upon optical uncaging. A drawback of a photocaging strategy, on the other hand, is that it is an irreversible transformation, unlike photoswitchable derivatives. By deploying GLPLight1 and photo-GLP1 in concert in an all-optical assay, we determined that the spatial spread of GLP1R activation in response to GLP-1 release can be localized to single-cells or even sub-cellular domains. Furthermore, by combining a state-of-the-art cAMP sensor with photo-GLP1, we demonstrated the optical control of hmGLP1R-dependent downstream cellular signaling with single-cell resolution, opening exciting new opportunities for investigating the spatial regulation of this signaling pathway. Since we photocaged native GLP1, it is important to note, that the photo-GLP1 might still be susceptible to DPPIV-mediated degradation when used in *in vivo* applications. We envisage that our photo-GLP1 will nonetheless find applications in neurobiological *in vivo* studies in brain tissue, as DPPIV-levels in the brain are significantly lower than in peripheral organs.

In summary, we developed and utilized a new all-optical toolkit to unveil a previously inaccessible spatial dimension of the GLP-1/GLP1R system. These tools may thus be readily implemented in a variety of applications, some of which are showcased as part of this study, to advance our understanding of the roles of GLP-1/glucagon/GLP1R signaling system in physiology, or to foster the drug screening and development process targeting the GLP1R pathway.

## DATA AVAILABILITY

DNA plasmids used for viral production have been deposited both on the UZH Viral Vector Facility (https://vvf.ethz.ch/) and on AddGene (plasmid numbers: 187466-187468). Plasmids and viral vectors can be obtained either from the Patriarchi laboratory, the UZH Viral Vector Facility, or AddGene.

Source data are provided with the manuscript.

## ACKNOWLEDGEMENTS

The results are part of a project that has received funding from the European Research Council (ERC) under the European Union’s Horizon 2020 research and innovation program (Grant agreement No. 891959) (T.P.). We also acknowledge funding from the University of Zürich and the Swiss National Science Foundation (Grant No. 310030_196455 and 310030L_212508) (T.P.) and (Grant No. 200021_200865) (N.H.). We would like to thank Jean-Charles Paterna and the Viral Vector Facility of the Neuroscience Center Zürich (ZNZ) for the help with virus production. All plasmids encoding miniG proteins-LgBit were a kind gift from Nevin A. Lambert (University of Augusta). The plasmids encoding Beta2AR-SmBit and Beta-Arrestin-LgBit, as well as the Alexa-647 labelled M1 anti-FLAG antibody, were a kind gift from Miriam Stoeber (University of Geneva). The GLUTag enterocrine cell line was a kind gift from Daniel J Drucker (University of Toronto).

## AUTHOR CONTRIBUTIONS

T.P. conceived the development and applications of GLPLight1. N.H. conceived the development of photo-GLP1 and GLP-1 derivatives. T.P. and N.H. co-supervised the study. L.D. and A.G. performed molecular cloning and *in vitro* sensor library screening under the supervision of T.P. L.D. performed all sensor imaging and characterization experiments in HEK293 cells and neurons, including kinetics, signaling, FACS titrations, and all-optical assays, and analyzed data under the supervision of T.P. M.A.B. prepared cortical neuronal cultures under the supervision of D.B. E.T.W. designed, synthesized, and characterized photo-GLP1 under the supervision of N.H. S.C. synthetized the GLP-1 alanine mutants under the supervision of N.H. T.P., L.D., N.H. and E.T.W. wrote the manuscript with contributions from all authors.

## Material and methods

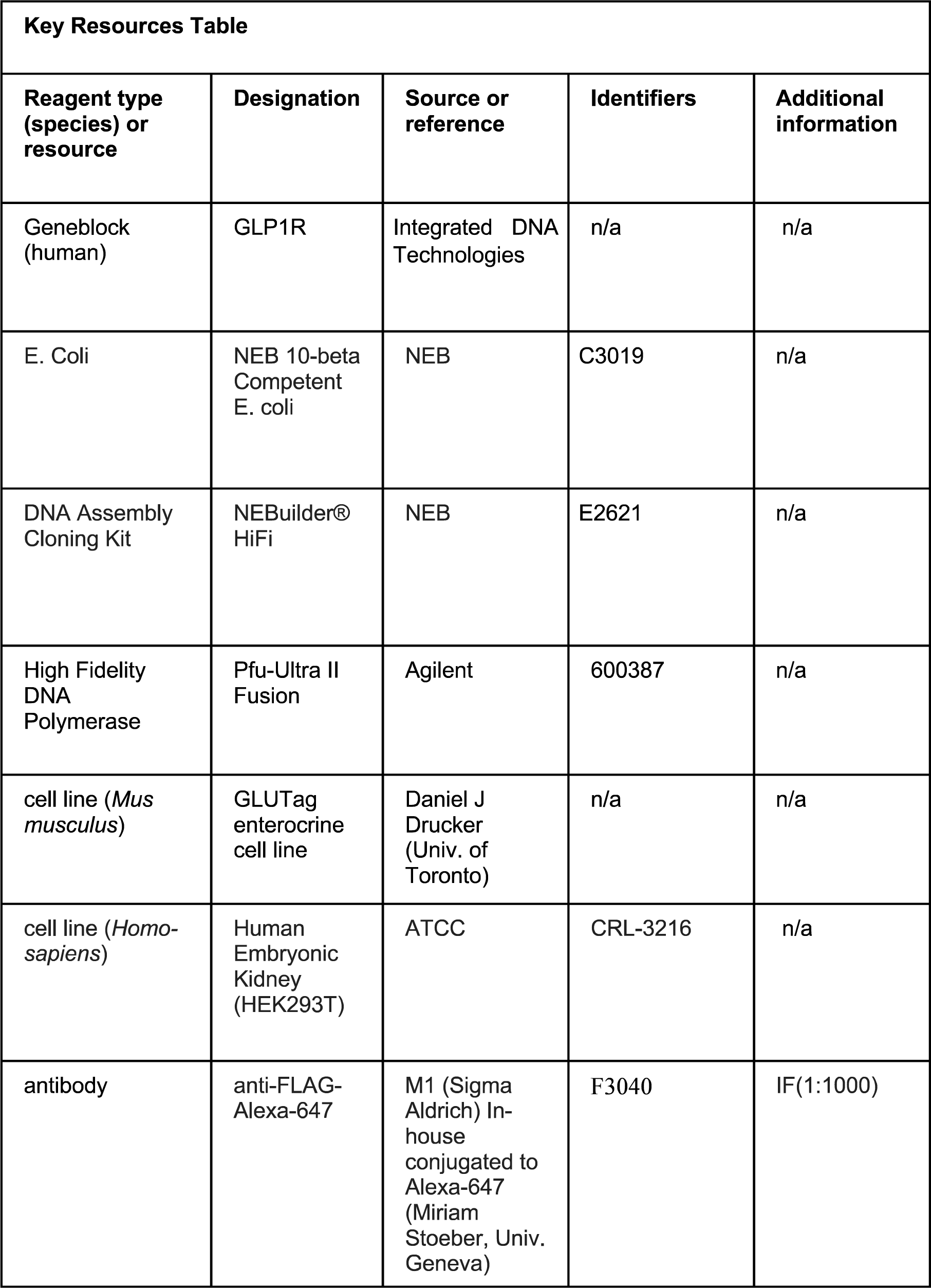

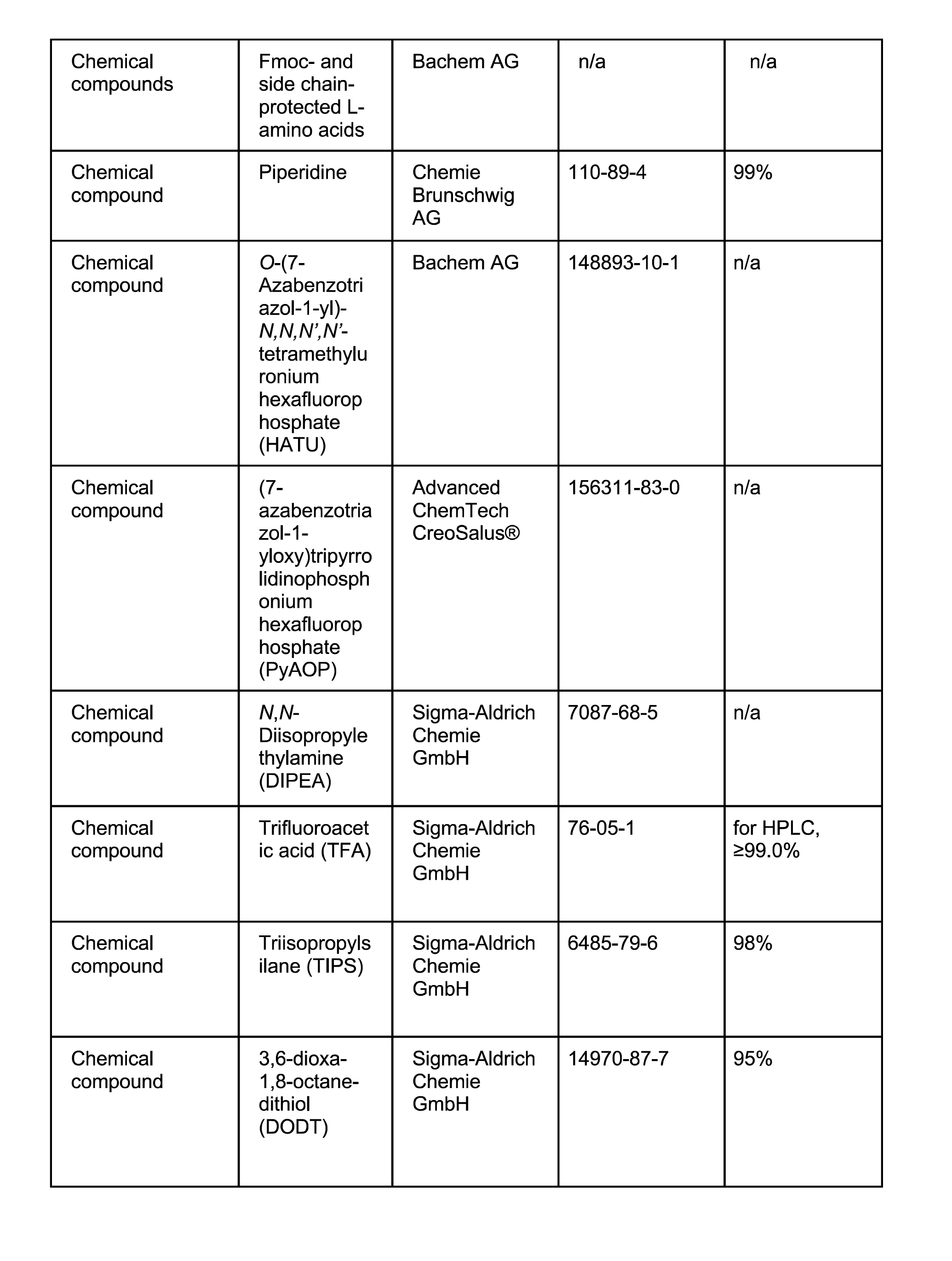

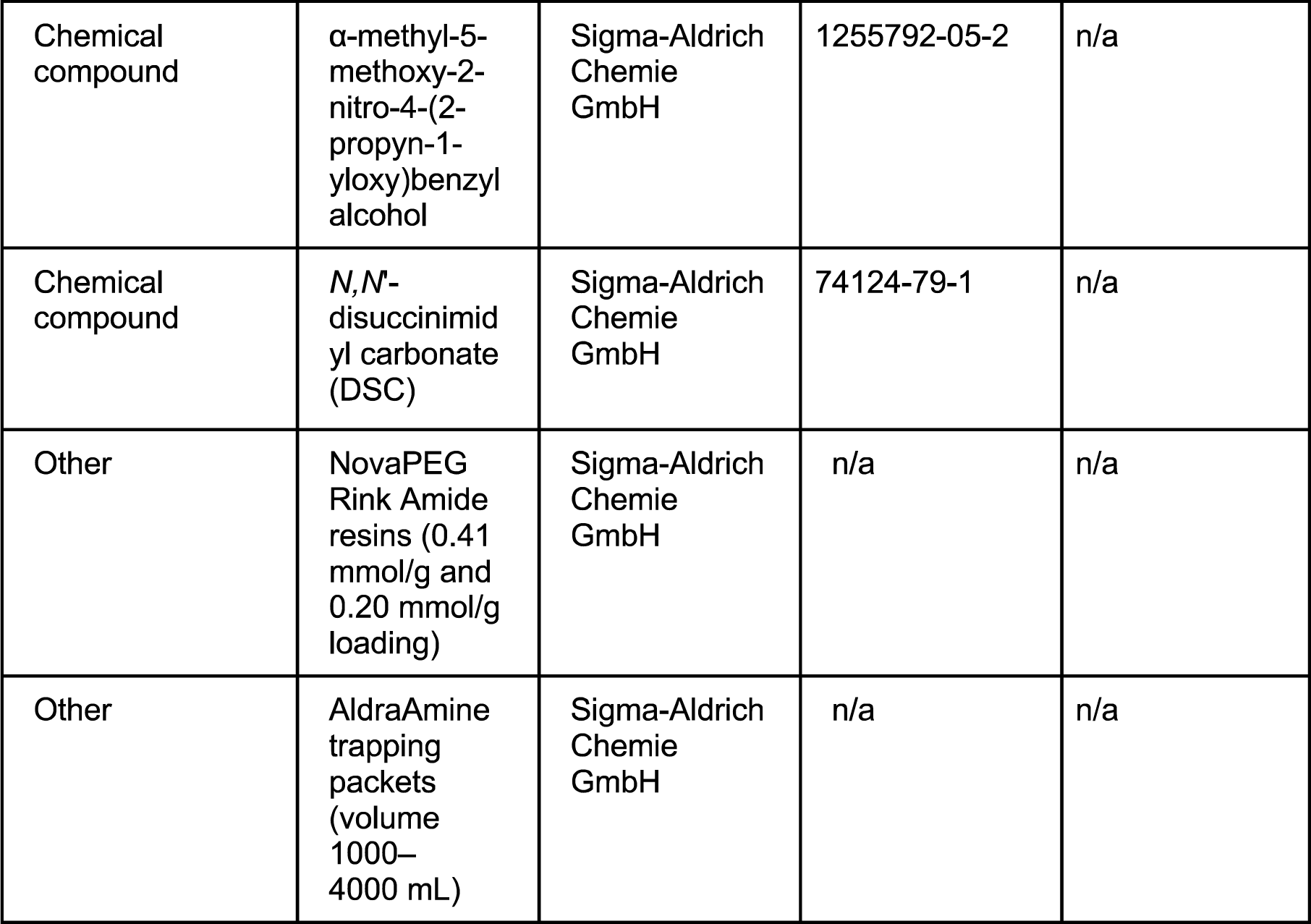

### Molecular cloning

The sequence coding for human GLP1R was ordered as a synthetic DNA geneblock (Integrated DNA Technologies) bearing HindIII and NotI restriction site for cloning into a CMV-promoter plasmid (Addgene #60360). Sequences coding for the hemagglutinin secretion motif and a FLAG Tag were added to the N-terminus of the GLP1R open reading frame to increase plasma membrane expression and enable receptor labelling, respectively. Sensor variants were obtained either using Gibson assembly (NEBuilder® HiFi DNA Assembly Cloning Kit) (Gibson *et al*., 2009). Site-saturated mutagenesis was performed by PCR using primers bearing randomized codons at specified locations (NNK). For luminescence-based characterization of G protein and Beta-arrestin coupling, the small subunit (ie. smBit) of NanoLuc (Cannaert *et al*., 2016) was PCR-amplified from a Beta2AR-SmBit donor plasmid and cloned at the C-terminal end of the GLP1R and GLPLight1 using Gibson assembly. PCR reactions were performed using a Pfu-Ultra II Fusion High Fidelity DNA Polymerase (Agilent). All sequences were verified using Sanger sequencing (Microsynth). For cloning GLPLight1 and GLPLight-ctr into the viral vector, BamHI and HindIII restriction sites were added flanking the sensor coding sequence by PCR-amplification, followed by restriction cloning into pAAV-hSynapsin1-WPRE, obtained from the Viral Vector Facility of the University of Zürich.

### Structural modelling

The structural model of GLPLight1 was obtained using ColabFold (Mirdita *et al*., 2022) using pdb70 as a template mode. The best prediction was selected manually and edited using Chimera.

### Peptide synthesis and biochemical characterization

GLP-1, photo-GLP1 and all alanine scan peptides were synthesized on an automated fast-flow peptide synthesizer (AFPS) using a recently developed protocol (Hartrampf *et al*., 2020). A detailed description of the synthetic procedures and all analytical data can be found in the Supplementary Information.

### Cell culture, imaging and quantification

Mammalian HEK293T cells (CRL-3216 from ATCC) were authenticated by the vendor and tested negative for mycoplasma. They were cultured in DMEM medium (Thermo Fisher) supplemented with 10% FBS (Thermo Fisher) and 1X final Antibiotic-Antimicotic (Thermo Fisher) and incubated at 37 °C in 5% CO_2._ The cells were transfected using Effectene transfection kit for individual dishes or 24 well plates (Qiagen) or Linear PEI (Sigma Aldrich) for T75 flask transfection following the manufacturer’s instructions and imaged 24-48h after transfection. GLUTag enterocrine cells were authenticated by the provider and tested negative for mycoplasma. They were cultured on plates coated with 0.1% gelatine (Sigma Aldrich) in low-glucose DMEM medium (1g/L glucose) supplemented with L-glutamine (4 mM) and pyruvate (1 mM), 10% FBS and 1% Pen/Strep (Thermo Fisher). Primary cortical neurons were prepared as follows: the cerebral cortex of 18 days old rat embryos were carefully dissected and washed with 5 mL sterile-filtered PBGA buffer (PBS containing 10 mM glucose, 1.0 mg/mL bovine serum albumin and antibiotic-antimycotic 1:100 (10,000 units/mL penicillin; 10,000 μg/mL streptomycin; 25 μg/mL amphotericin B)) (ThermoFisher Scientific). Cortices were cut into small pieces and digested in 5.0 mL sterile filtered papain solution for 15 minutes at 37 °C. Tissues were then washed with complete DMEM medium containing 10% Fetal Calf Serum and penicillin/streptomycin (1:100), triturated and filtered through a 40 μm cell-strainer. Neurons were plated at a concentration of 40,000– 50,000 cells per well onto poly-L-lysine (50 μg/mL in PBS, ThermoFisher Scientific) coated dishes and kept in NU-medium (Minimum Essential Medium (MEM) with 15% NU serum, 2% B27 supplement, 15 mM HEPES, 0.45% glucose, 1.0 mM sodium pyruvate, 2.0 mM GlutaMAX). The cultures were virally transduced after 4–6 days with adeno-associated viruses at a 1 × 10^9^ GC/mL final titer and kept for 12– 16 days in vitro. The HEK293T cells or neurons were rinsed with HBSS (Hank’s Balanced Salt Solution, Life Technologies) and kept in a final volume of HBSS being either 100 µL for individual 15 mm glass bottom insert dish or 500 µL for 24-well plates. Time lapse recordings were performed at room temperature (approx. 20 °C) on a Zeiss LSM 800 inverted confocal microscope controlled by Zeiss Zen Blue 2018 v2.6 software using either a 40X oil-based objective (individual dishes) or 20X air objective (24-well plates). The probes were excited using the following laser lines: 488 nm for GLPLight1 and GLPLight1-ctr. The ligands were all added in bolus before or during the time-lapse recording using a micropipette to reach the desired final concentration once mixed with HBSS imaging media. Optical uncaging was performed using a 40x Plan-Apochromat oil-based objective (N/A = 1,4; 69% transmittance at 405 nm from manufacturer’s datasheet) over specified surface areas with various scanning rates (described in each legend) and a pixel dwell time of 1.52 µsec. The average intensity of laser light used for uncaging was measured using a S120C Photodiode Power Sensor from Thorlabs and was kept at 0.38 mW. Image quantification was performed after manual selection of the regions of interest (ROI) corresponding to the cell membrane using the thresholding function from Fiji. The sensor response (ΔF/F_0_) was calculated as followed: F_t_-F_0_)/F_0_ with F_t_ being the fluorescence intensity of the ROI at each time point t, and F_0_ being the mean fluorescence intensity of the 10 time points before ligand addition for each ROI. ΔF/F_0_ values were calculated using a custom-made MatLab script and plotted in GraphPad Prism. The ΔF/F_0_ images were obtained by dividing pixel-wise fluorescence intensities prior and post ligand addition using a separate MatLab script and displayed as a color-coded RGB image.

### Plate reader-based imaging

The spectral characterization of the sensor was performed using GLPLight1 transfected HEK293T cells pre and post 10 µM GLP-1(7**–**37) addition. The excitation and emission spectra were measured at λ_em_ = 560 nm and λ_ex_ = 470 nm, respectively, on a TECAN M200 Pro plate reader at 37 °C. Transfected or untransfected cells were lifted using Versene^®^ (Thermo Fisher Scientific) and resuspended in PBS at a concentration of 3.3 million cells per mL. For each condition, 300 µL of the cell suspension or PBS was transferred per individual wells of a black bottom 96-well plate. Untransfected cells were used to correct for autofluorescence whereas PBS alone was used to subtract the buffer Raman bands. Intracellular cAMP production was assessed with the GloSensor™ cAMP assay. HEK293T cells were co-transfected with the pGLO20F and either human GLP1Ror GLPLight1 in separate T75 flasks. Note that the endogenous signal peptide (amino acids 1**–**23) from GLP1R WT was deleted to maintain a similar membrane expression compared to GLPLight1 for all signaling assays. Cells were lifted 24 h after transfection using Versene^®^ and re-suspended at a final concentration of 1,500,000 cells per mL in DMEM without phenol red + 15 mM HEPES (Thermo Fisher Scientific). 100 µL of the cell suspension was dispensed per well in a 96-well white plate (Corning) and incubated with 2.0 mM of Luciferin potassium salt in 10 mM HEPES (pH 7.4) for 45–60 minutes. The cells were then imaged right after addition of 50 µL of ligand to reach the desired final concentration using a Cytation C10 (Biotek) plate reader in kinetic luminescence mode at 37 °C. Positive (2.5 mM Forskolin) and negative controls (assay medium) were always included in triplicate alongside the constructs to be tested. The dose-response curves were obtained using the average luminescence value of the 5 timepoints after the peak of cAMP production of the positive control. Luminescence complementation assays were conducted using HEK293T cells co-transfected with GLPLight1-SmBit or GLP1R-SmBit along with either miniGs-LgBit, miniGi-LgBit, miniGq-LgBit, miniG12-LgBit or Beta-arrestin-2-LgBit. After transfection, cells were seeded in a 96-well Optiplate^®^, using 10,000 cells per well for the miniGs-LgBit condition and 50,000 cells for all others. Cells were then incubated for 45–60 minutes at 37 °C with the NanoGlo^®^ live cell reagent according to the manufacturer’s instructions. The baseline luminescence was recorded for 100 cycles (approx. 460 sec), paused for manual addition of the ligand or the vehicle and resumed for another 200 cycles (approx. 920 sec). The ΔR/R_0_ values were calculated by dividing the raw luminescence intensities after GLP-1(7-37) addition by the ones after vehicle addition. This ratio was then normalized using the average luminescence intensity before addition as a baseline for both GLPLight1 and GLP1R conditions. The quantification of the maximal recruitment was calculated using the average ΔR/R_0_ between t = 600 sec and t = 700 sec for the time-lapses and t = 1600 sec and t = 1800 sec for the GLP-1 titration of miniGs recruitment to GLP1R.

### Flow cytometry

After transfection, HEK293T cells were harvested using Versene^®^. After resuspension in FACS buffer (1 × PBS, 1.0 mM EDTA, 25 mM HEPES pH 7.0, 1% FBS) 300,000 cells were dispensed in each well of a 96-well plate, mixed with an equivalent volume of ligand to reach the desired concentration, and were incubated for 30 minutes at room temperature before the start of the measurement. Transduced neurons were washed once using the FACS buffer, gently mechanically lifted using a cell scraper and homogenized by repeated up and down pipetting in FACS buffer. They were then incubated for 15 minutes on ice to minimize cell death before measurement. All flow cytometry experiments were performed on a FACS Canto II 2L using a high-throughput sampler (HTS). Forward scattering (FSC), side-scattering (SSC) and 488 nm-excited fluorescence (FITC) data were acquired for a total of 50,000– 100,000 events per well. The cells were manually gated using non-expressing cells as comparison, to define the FITC-positive population. Within this subgroup, the mean FITC intensity was calculated for each condition and normalized to the maximum FITC signal.

### Virus production

The AAV biosensors constructs used in this study were cloned by the Patriarchi laboratory. The VVF provided the backbone AAV constructs and produced the viruses. The titer of the viruses used were: AAVDJ.hSynapsin1.GLPLight1, 3.7 × 10^12^ VG/mL; AAVDJ.hSynapsin1.GLPLight-ctr, 3.4 × 10^12^ VG/mL.

### Animals

Animal procedures were performed in accordance to the guidelines of the European Community Council Directive or the Animal Welfare Ordinance (TSchV 455.1) of the Swiss Federal Food Safety and Veterinary Office and were approved by the Zürich Cantonal Veterinary Office. Rat embryos (E17) obtained from timed-pregnant Wistar rats (Envigo) were used for preparing primary cortical neuronal cultures.

### Statistical analyses

For in vitro analysis of sensor variants, where relevant the statistical significance of their responses was determined using a two-tailed unpaired Student’s t-test with Welch’s correction. For comparison of uncaging events in the presence or absence of antagonist statistical analysis was performed using Brown-Forsythe ANOVA test followed by Dunnett’s T3 multiple comparison. For comparison of kinetic measurements, statistical analysis was performed using the extra sum-of-squares F test. All numbers of experimental repeats and p values are reported in the figure legends. Error bars represent mean ± standard error of the mean (SEM).

**Figure 1 – figure supplement 1.**
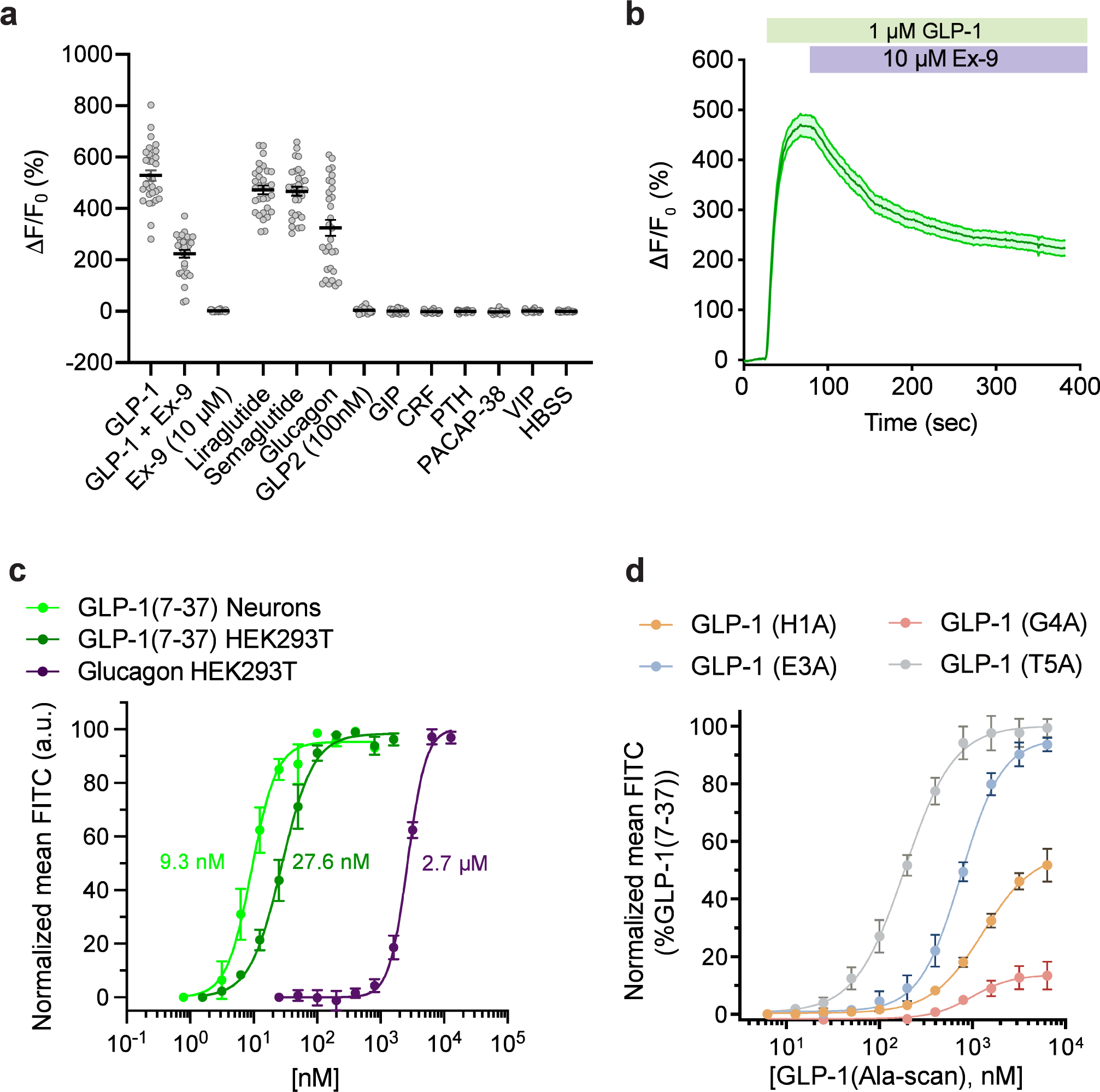
Screening process for the development of GLPLight1. **a)** Maximal fluorescence response to 10 μM GLP-1 of the prototype GLP-1 sensor and the N-terminus deletion variants (16 or 23 first amino acids). **b**) Maximal fluorescence response to 10 μM GLP-1 of the ICL2 lysine variants compared to GLP-1 sensor prototype with deletion of the first 23 amino acids. The variant with the best response containing the mutation L260K was named GLPLight0.1. The top-right insert depicts the site that were mutated on the sensor’s ICL2 in black letters. **c**) Overview of the randomized screening of residues K336 and T343 on GLPLight0.1 backbone after addition of 10 μM GLP-1. The data were normalized to the variant showing the highest response. The two randomized residues are shown above in the diagram as the black letters: x. **d**) Comparison of the maximal response to 10 μM GLP-1 of GLPLight0.1 to the best variant from **c,** subsequently called GLPLight0.2. **e**) Comparison of the maximal response to 10 μM GLP-1 of GLPLight0.2 constructs with or without addition of an ER export sequence at the C-terminus. The construct containing the ER export sequence was called GLPLight0.3. **f**) Comparison of the maximal response to 10 μM GLP-1 of GLPLight0.3 with or without the cpGFP mutations as well as their basal brightness in **g**. The construct containing the mutations was called GLPLight0.4. **h**) Comparison of the maximal response to 10 μM GLP-1 of GLPLight0.4 with or without the alanine mutations of the C-terminal phosphorylation sites. The final version of the sensor was called GLPLight1. All data were acquired in HEK293T cells with multiple repeats shown as mean ± SEM.

**Figure 1 – figure supplement 2.**
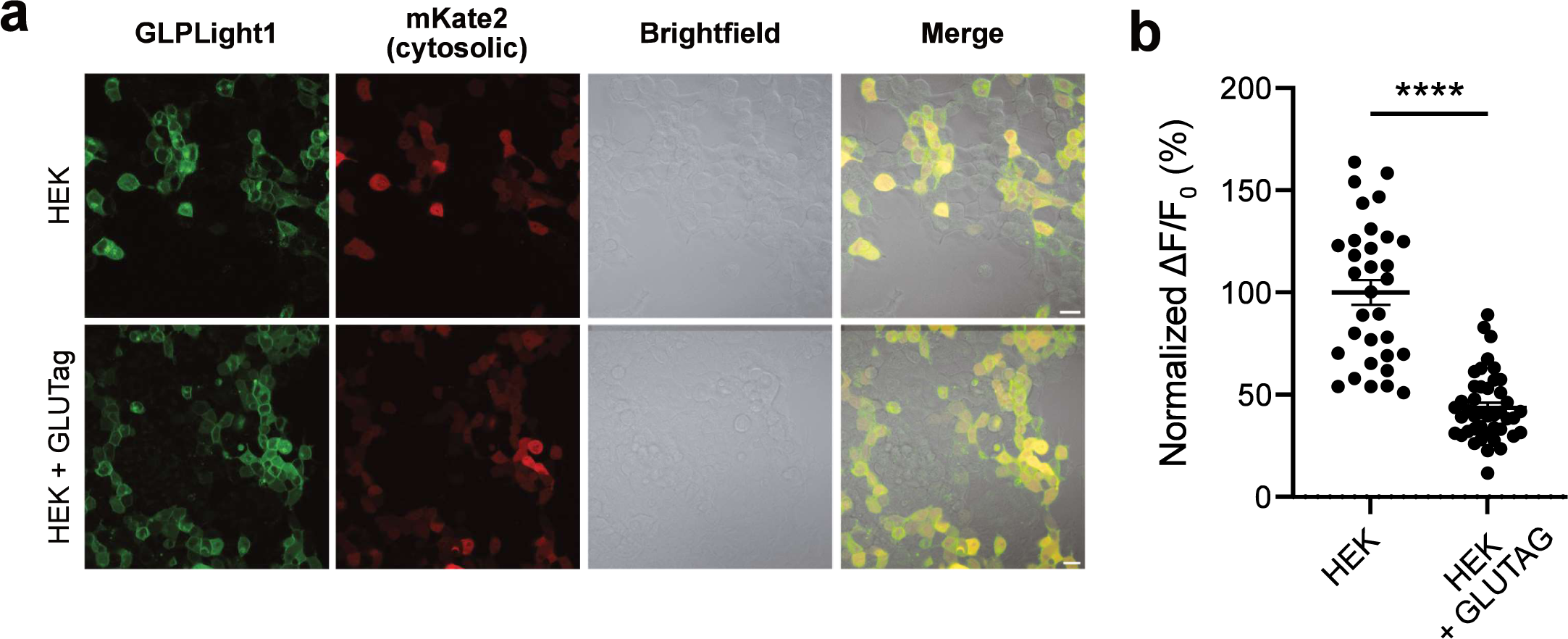
Development of the control sensor GLPLight-ctr. **a)** Maximal response of GLPLight1 binding pocket variants to 1 μM GLP-1(7**–**37). **b**) Structural model of GLPLight-ctr shown from the side (left) or top (right) perspective. The residues mutated to alanine compared to GLPLight1 are shown in magenta. **c**) Representative images from GLPLight-ctr expression and fluorescence intensity change before (left) and after (center) addition of 10 μM GLP-1(7**–**37) as well as the respective pixel-wise ΔF/F_0_ images in HEK293T cells (top) and primary cortical neurons (bottom). Scale bars: 10 μm for HEK cells and 20 μm for neurons. d) Maximal response of either GLPLight1 or GLPLight-ctr in HEK293T cells and primary cortical neurons after addition of 10 μM GLP-1(7**–**37).

**Figure 1 – figure supplement 3.**
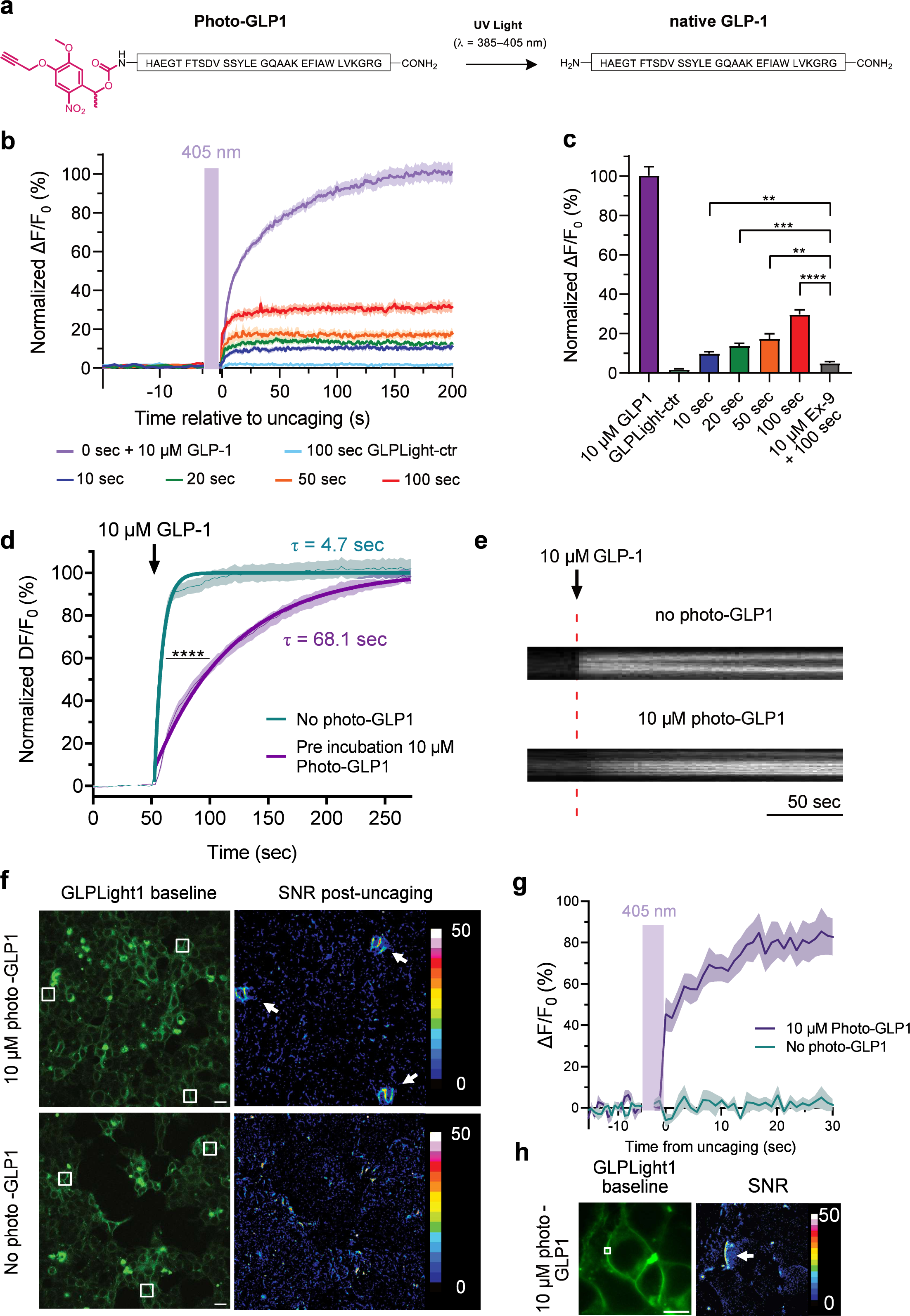
Intracellular signaling characterization of GLP1R and GLPLight1. Normalized timelapse recording of agonist induced miniG protein or beta arrestin recruitment in GLP1R or GLPLight1 expressing HEK293T cells along with quantification of the mean signal between t = 600**–** 700 sec for mini-Gs in **a–b**, mini-Gq in **c–d**, mini-Gi in **e–f**, mini-G12 in **g–h**, and β-arrestin-2 in **i–j**. The dashed line corresponds to the addition of 100 nM GLP-1(7-37). One-tailed unpaired t-test with Welch’s correction, mini-Gs: p = 0,0202; mini-Gq: p = 0,0074; mini-Gi: p = 0,0102; mini-G12: p = 0,0356 and β-arrestin-2: p = 0,0452. **k**) Titration of GLP-1 induced intracellular cAMP in HEK293T cells co-expressing GLO20F cAMP sensor and GLP1R or GLPLight1. The data were normalized to the maximum signal observed for GLP1R. Unpaired two-tailed t-test with Welch’s correction for both constructs after 100 nM GLP-1. P = 0,0041. **l**) Titration of GLP-1 on miniGs-recruitment assay in GLP1R expressing HEK293T cells. The curve fit was performed using a four parameter non-linear fit and the EC_50_ value is shown next to the trace. All data obtained from three independent experiments.

**Figure 2 – figure supplement 1.**
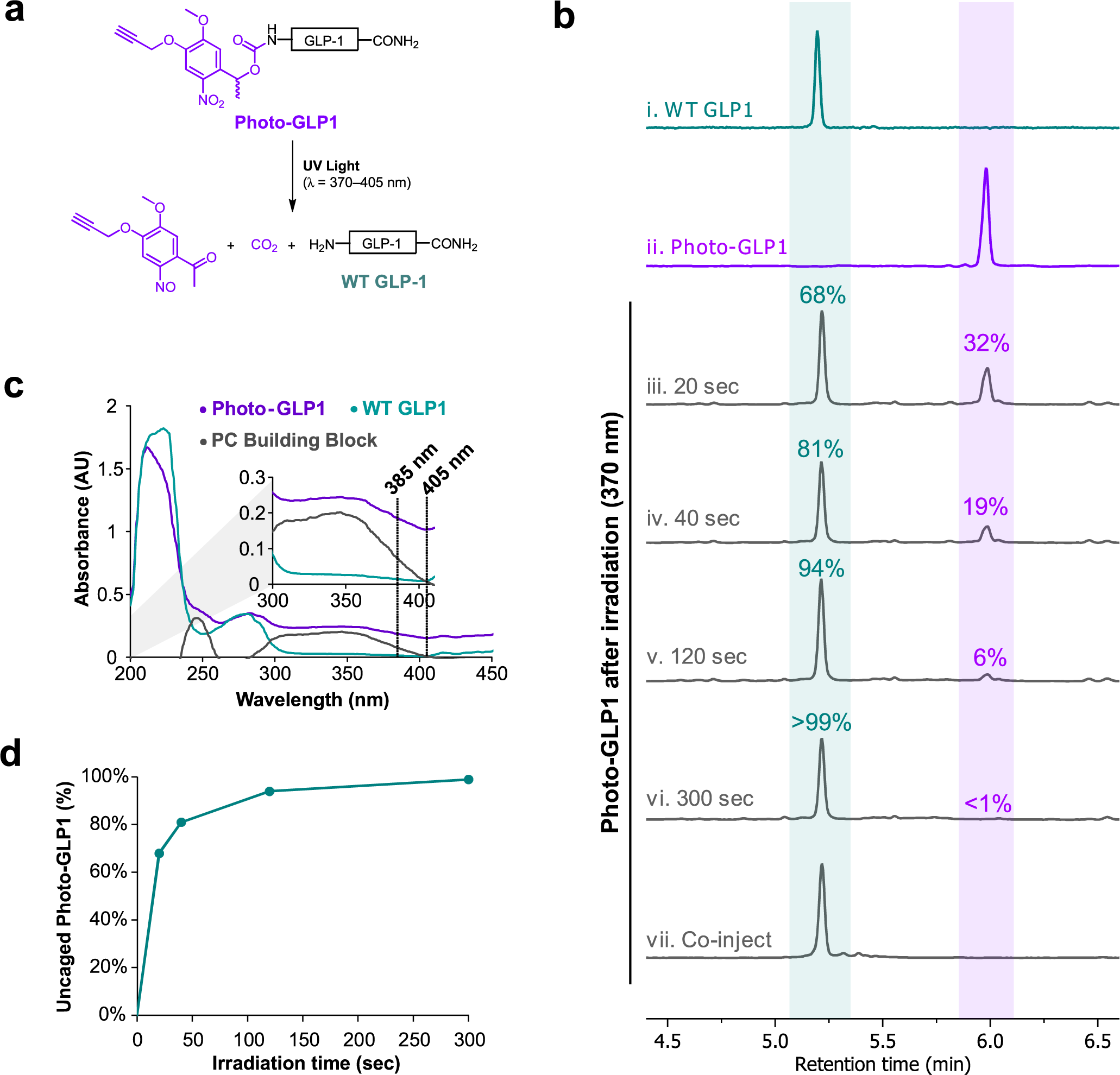
Detecting endogenous GLP-1 release from enterocrine cells using GLPLight1. **a)** Representative fluorescence and brightfield images of HEK293T co-transfected with GLPLight1 and cytosolic mKate2 cultured overnight in the absence (top row) or presence (bottom row) of GLUTag cells. **b)** Fluorescence response of GLPLight1 expressing HEK cells cultured with (right) or without (left) GLUTag cells. The data were normalized to the average response of GLPLight1 from the HEK cells only population. Statistical analysis was done using a two-tailed unpaired t-test, ****P = 3.399 × 10^-14^; n = 32 and 43 cells from three independent experiments for HEK cells only or HEK+GLUTag cells, respectively.

**Figure 3 – figure supplement 1.**
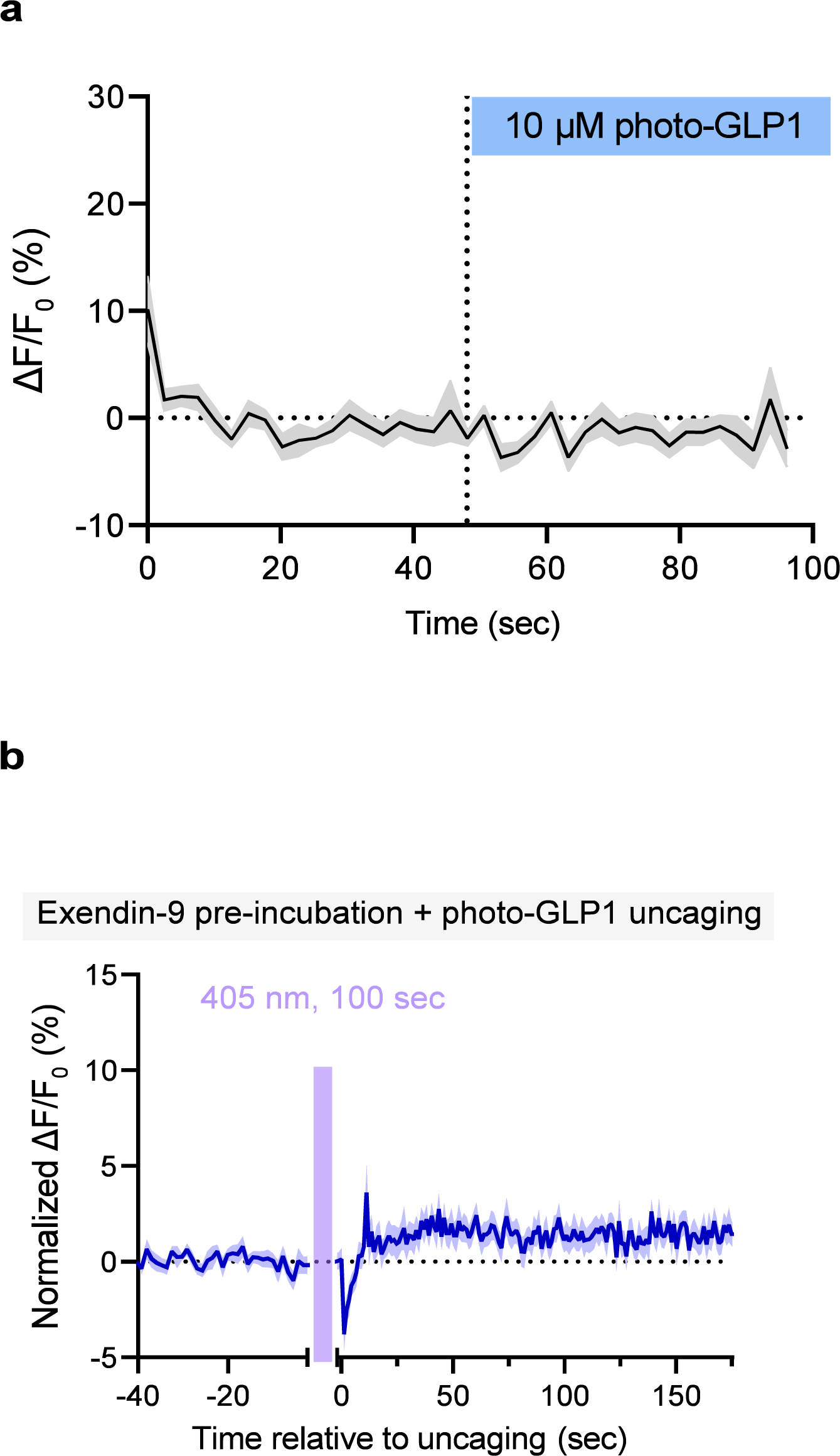
Biochemical characterization of photo-GLP1. **a)** Scheme illustrating the uncaging reaction of photo-GLP1 when exposed to UV light at 370–405 nm, producing WT GLP-1. **b**) LCMS chromatographic traces of (i) pure WT GLP-1, (ii) pure photo-GLP1, photo-GLP1 after irradiation at 370 nm for (iii) 20 sec, (iv) 40 sec, (v) 120 sec, (vi) 300 sec, and (vii) co-injection of photo-GLP1 after irradiation for 300 sec (50 µL, 80 µM) with pure WT GLP-1 (50 µL, 80 µM). Chromatographic peaks corresponding to WT GLP-1 are highlighted in blue/green, peaks corresponding to photo-GLP1 are highlighted in purple. **c**) UV-Vis spectra (λ = 200–450 nm) of photo- GLP1 (purple, 80 µM), WT GLP-1 (blue/green, 80 µM), and α-methyl-5-methoxy-2-nitro-4-(2-propyn-1- yloxy)benzyl alcohol (PC Building Block, grey, 80 µM). All UV-Vis measurements were carried out in HBSS buffer and processed with blank (HBSS buffer) subtraction. **d**) Amount of WT GLP-1 produced by irradiation of photo-GLP1 at 370 nm over time. Amount of WT GLP-1 at each time-point was estimated using area under the curve (AUC) of the corresponding chromatographic peak and given as a percentage of the total AUC value of peaks corresponding to photo-GLP1 and WT GLP-1.

**Figure 3 – figure supplement 2.**
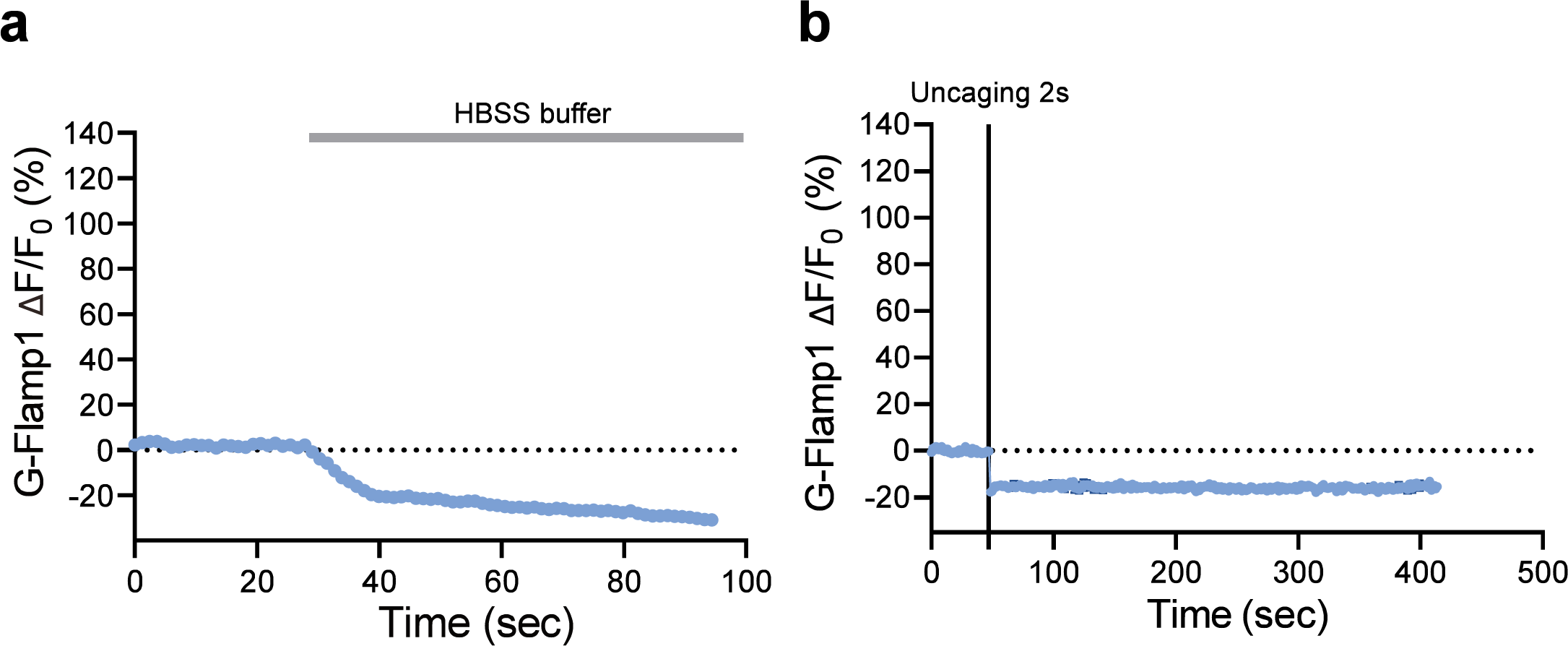
Further characterization of photo-GLP1 uncaging. **a)** Fluorescence response (ΔF/F_0_) of in GLPLight1 expressing HEK293T cells after bolus addition of 10 μM photo-GLP1. The time of addition is represented by the blue rectangle above; n = 16 cells. **b**) Fluorescence response of GLPLight1 expressing HEK293T cells normalized to the maximal response to 10 μM WT GLP-1 after 2 minutes pre-incubation with a mix of 10 μM Exendin-9 + 10 μM photo-GLP1 and following optical uncaging for 100 seconds with a 405 nm laser (represented by the magenta shaded area); n = 17 cells. All data shown as mean ± SEM and acquired from 3 independent experiments.

**Figure 4 - figure supplement 1.**
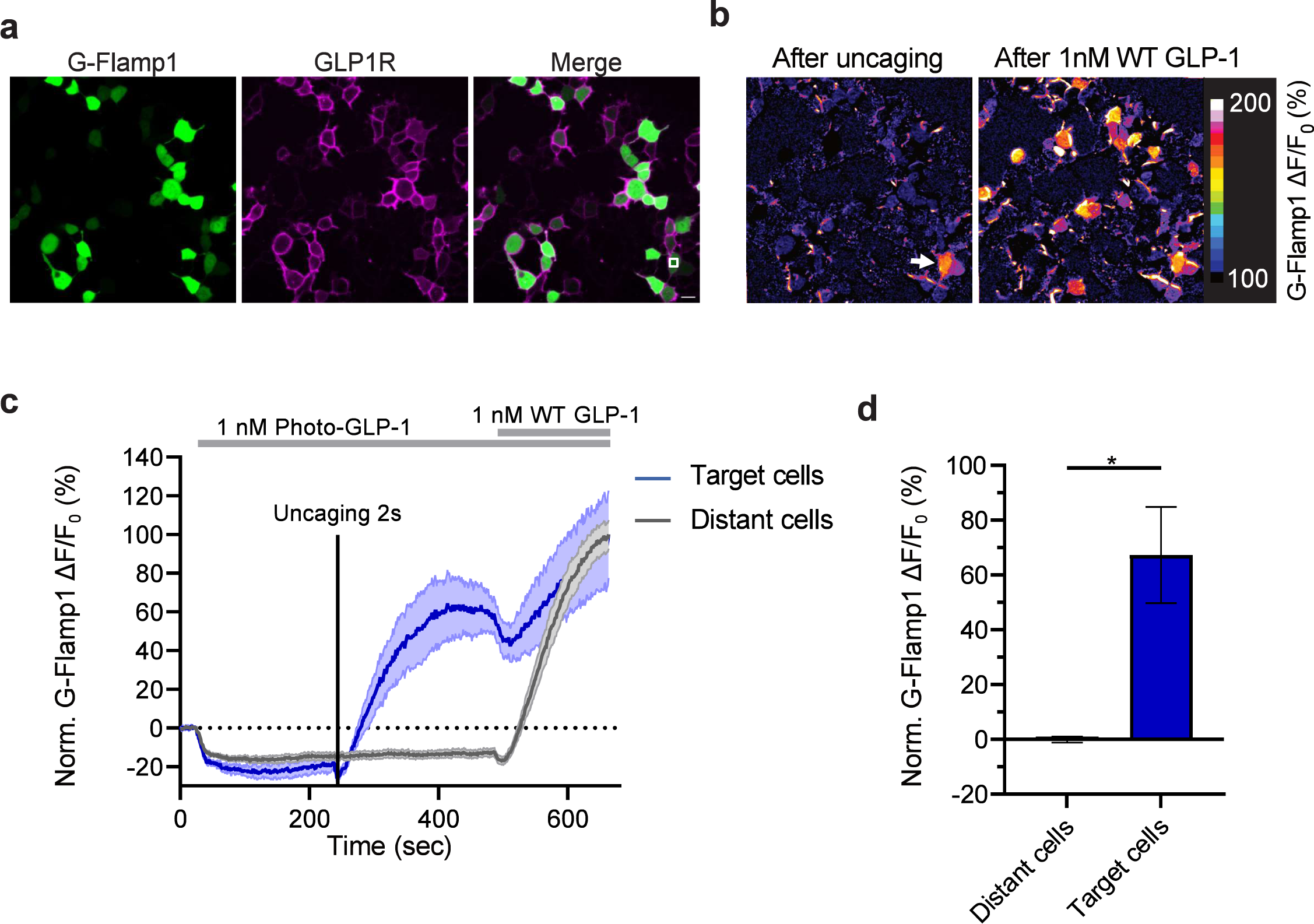
Control experiment for intracellular signaling characterization. **a)** Timelapse imaging of G-Flamp1 response in HEK293T cells co-expressing GLP1R after addition of HBSS buffer (used for imaging and dilution of the ligands). The time point of buffer addition is indicated by the grey rectangle above. n = 15 cells from three independent experiments. **b**) Timelapse imaging of G-Flamp1 response in HEK293T cells co-expressing GLP1R after 2 seconds photo-activation (represented by the black vertical line) in the absence of photo-GLP1. n = 3 cells from three independent experiments.

## Synthesis and Characterization of Building Blocks and Peptides

### 1 Materials and Equipment

Unless otherwise noted, all reactions were performed under an atmosphere of inert gas (N_2_). The reactions were carried out in oven-dried glassware using dry solvents, unless otherwise noted. Room temperature is defined as a range between 20–25 °C.

#### 1.1 Chemicals and Solvents

All chemicals and solvents were used as supplied, unless otherwise stated. Fmoc- and side chain-protected L-amino acids (Fmoc-Ala-OH, Fmoc-Arg(Pbf)-OH, Fmoc-Asn(Trt)-OH, Fmoc-Asp(*t*Bu)-OH, Fmoc-Cys(Trt)-OH, Fmoc-Gln(Trt)-OH, Fmoc-Glu(*t*Bu)-OH, Fmoc-Gly-OH, Fmoc-His(Trt)-OH, Fmoc-Ile-OH, Fmoc-Leu-OH, Fmoc-Lys(Boc)-OH, Fmoc-Met-OH, Fmoc-Phe-OH, Fmoc-Pro-OH, Fmoc-Ser(*t*Bu)-OH, Fmoc-Thr(*t*Bu)-OH, Fmoc-Trp(Boc)-OH, Fmoc-Tyr(*t*Bu)-OH, Fmoc-Val-OH) were purchased from Bachem AG or the Novabiochem-line from Sigma-Aldrich Chemie GmbH (Pbf = 2,2,4,6,7- pentamethyldihydrobenzofuran-5-sulfonyl, Trt = trityl, *t*Bu = *tert*-butyl, Boc = *tert*-butoxycarbonyl). Piperidine (99%) was purchased from Chemie Brunschwig AG. *O*-(7-Azabenzotriazol-1-yl)-*N,N,N’,N’*- tetramethyluronium hexafluorophosphate (HATU) and (7-azabenzotriazol-1- yloxy)tripyrrolidinophosphonium hexafluorophosphate (PyAOP) were purchased from Bachem AG and Advanced ChemTech CreoSalus®, respectively. *N*,*N*-Diisopropylethylamine (*i*Pr_2_NEt, DIPEA, 99.5%), trifluoroacetic acid (TFA, for HPLC, ≥99.0%), triisopropylsilane (TIPS, 98%), 3,6-dioxa-1,8-octane-dithiol (DODT, 95%), α-methyl-5-methoxy-2-nitro-4-(2-propyn-1-yloxy)benzyl alcohol, and acetonitrile (MeCN, HPLC gradient grade, ≥99.9%) were purchased from Sigma-Aldrich Chemie GmbH. *N,N*’-disuccinimidyl carbonate was purchased from the Novabiochem-line from Sigma-Aldrich Chemie GmbH. *N,N*-Dimethylformamide (DMF) was purchased from the Supelco-line from VWR International GmbH. Dichloromethane (DCM, ≥99.8%) was purchased from Fischer Scientific Inc. Diethyl ether was purchased from Honeywell Riedel-de Haën. Dry MeCN (99.9%, AcroSeal®, stored over molecular sieves) was purchased from Acros Organics. Hexanes (technical grade) and EtOAc (technical grade) used for flash column chromatography were purchased from Thommen-Furler AG and distilled under reduced pressure prior to use. NovaPEG Rink Amide resins (0.41 mmol/g and 0.20 mmol/g loading) were purchased from the Novabiochem-line from Sigma-Aldrich Chemie GmbH. Et_3_N (anhydrous) was purchased from Fluorochem Ltd. AldraAmine trapping packets (volume 1000–4000 mL) were purchased from Sigma-Aldrich Chemie GmbH.

#### 1.2 Chromatography

Analytical thin-layer chromatography (TLC): Merck TLC plates (silica gel 60) on glass with the indicated solvent system. TLC spots were visualized by UV light (254 nm) and by stains of KMnO_4_ or Ninhydrin. Flash column chromatography was performed using Merck silica gel 60 (40−63 μm particle size) with the indicated solvent system.

#### 1.3 UV-Vis Spectroscopy

UV-Vis spectra were acquired using a Multiskan SkyHigh Microplate Spectrophotometer (ThermoFisher Scientific Inc.) in the range of 200–450 nm with 1 nm steps and quartz cuvettes (10 mm pathlength, Hellma Schweiz AG).

#### 1.4 NMR Spectroscopy

Nuclear magnetic resonance (NMR) spectra were recorded in deuterated chloroform (CDCl_3_) or D_2_O in 5 mm tubes on a Bruker AV2-401 (400 MHz) spectrometer equipped with a BOSS-I shim system, a digital lock control unit, an AMOS Control System, a DQD unit, a BVT3200 with a BCU05 cooling unit, GRASP Level II for gradient spectroscopy. Chemical shifts (δ scale) are expressed in parts per million (ppm) and are calibrated using residual protic solvent as an internal reference (CHCl_3_: δ = 7.26 ppm). Data for ^1^H NMR (400 MHz) spectra are reported as follows: chemical shift (δ ppm) (multiplicity, coupling constants (Hz), integration). Couplings are expressed as: s = singlet, d = doublet, t = triplet, q = quartet, m = multiplet or combinations thereof. ^13^C NMR spectra were recorded at 126 MHz. Carbon chemical shifts (δ scale), expressed in parts per million (ppm), are referenced to the central carbon resonances of the solvents (CHCl_3_: δ = 77.2 ppm).

#### 1.5 Analytical LC-HR-ESI-MS (LCMS)

Liquid chromatography high resolution electrospray ionization mass spectrometry (LCMS): Acquity UPLC (Waters, Milford, USA) connected to an Acquity eλ diode array detector and a Synapt G2HR-ESI-QTOF-MS (Waters, Milford, USA); injection of 10 μL sample (c = *ca.* 10–100 μg/mL in the indicated solvent); Acquity BEH C8 HPLC column (1.7 μm particle size, 2.1 × 100 mm, Waters) kept at room temperature; elution at a flow rate of 0.3 mL/min with A: H_2_O + 0.02% HCO_2_H + 0.04% CF₃CO₂H and B: CH_3_CN + 0.04% HCO_2_H + 0.02% CF₃CO₂H, isocratic 5% B for 1 min; then 5–95% B over 9 min. UV-Vis spectra recorded in the range of 190–300 nm at 1.2 nm resolution and 20 points s^−1^; ESI: positive ionization mode, capillary voltage 3.0 kV, sampling cone 40 V, extraction cone 4 V, N_2_ cone gas 4 L/h, N_2_ desolvation gas 800 L/min, source temperature 120 °C; mass analyzer in resolution mode: mass range 150–3000 m/z with a scan rate of 1 Hz; mass calibration to <2 ppm within 50–2500 m/z with a 5 mM aq. soln. of HCO_2_Na, lock masses: m/z 195.0882 (caffeine, 0.70 ng/mL) and 556.2771 (Leucine-enkephalin, 2.0 ng/mL). All mass spectra are integrated across Rt = 4–6 min of the total ion count (TIC). The areas indicated in grey are excluded from integration, due to presence of the injection peak and solvent-mixing baseline perturbation. For purity of the final peptides, please refer to the UHPLC spectra. Deconvoluted masses from the raw m/z values are calculated using Mestrelab Research S.L.© MestReNova v. 14.1 Mnova MS Suite.

#### 1.6 Analytical Ultra High-Performance Liquid Chromatography (UHPLC)

For determination of purity by UHPLC, the filtered peptide solution was diluted in 10–50% acetonitrile (MeCN) in water with 0.1% TFA (500 μL) to a final concentration of approximately 0.5 mg/mL. The samples were measured on Agilent 1290 Infinity II Series UHPLC with UV detection at 214 nm and analyzed using Agilent OpenLab CDS and ChemStation software. Where specified, analytical UHPLC spectra were acquired wherein eluent A = MeCN/H_2_O (5:95) with 0.1% TFA, and eluent B = MeCN/H_2_O (95:5) with 0.1% TFA using either (i) an Agilent Zorbax 300SB-C18 Narrow-Bore column (2.1 mm × 150 mm, 5 µm particle size) kept at 40 °C, at a flow rate of 1.5 mL/min, with a gradient of 0% B for 3 min, followed by 0–100% B over 30 min (ca. 3% MeCN/min), then 0% B for a further 2 min (total method time was 35 min), or (ii) an Agilent Zorbax Eclipse Plus C18 Rapid Resolution HD column (2.1 × 50 mm, 1.8 µm particle size), with a gradient of 0% B for 1 min, followed by 0–100% B over 9 min (ca. 10% MeCN/min), then 0% B for a further 1 min (total method time was 11 min). Purities of the final peptides were calculated by integration of the Area Under the Curve (AUC) of desired product peak (detected at λ = 214 nm) as a percentage of the AUC of all peaks between the indicated timeframe.

### 2 General Procedures

#### 2.1 Solid-Phase Automated Fast-flow Peptide Synthesis (AFPS)

Peptides were synthesized on an automated-flow system, which was built in the Hartrampf lab, based on the published AFPS system.^[1]^ All peptides were prepared by AFPS on NovaPEG Rink Amide resin (0.41 mmol/g or 0.20 mmol/g, as specified) to afford C-terminally amidated peptides. All peptides were synthesized from the C- to N-terminus. Standard Fmoc/*t*Bu protected amino acids (0.40 M in DMF) were coupled to the solid support using HATU (0.38 M in DMF) or PyAOP (0.38 M in DMF) with DIPEA (neat, 3.0 mL/min) at an overall flow rate of 20 mL/min. Amino acids A, D, E, F, G, I, K, L, M, P, S, W, and Y were coupled using HATU. Amino acids N, Q, R, V, T, C, and H were coupled using PyAOP. All amino acids except C and H were preheated at 90 °C during the activation step with HATU or PyAOP, whereas C was preheated at 60 °C with PyAOP, and H was preactivated with PyAOP at room temperature. For amino acids D, E, F, G, I, K, L, M, P, S, W, and Y, each coupling was carried out for 19 sec at 20 mL/min. For amino acids A, C, H, N, Q, R, S, T, and V, couplings were carried out for 31 sec at 20 mL/min. Removal of the N^α^-Fmoc group was achieved using 20% piperidine with 1% formic acid in DMF at a flow rate of 20 mL/min at 90 °C for 19 sec. Between each coupling and deprotection step, the resin was washed with DMF (32 mL) at 90 °C with a flow rate of 40 mL/min.

#### 2.2 Cleavage of Peptidyl-Resins

After synthesis, the peptidyl-resin was washed with DCM (3 × 5 mL) and dried under reduced pressure. To the resin (amount as specified) was added cleavage solution (1.5–3.0 mL, TFA/TIPS/DODT/H_2_O 94/1/2.5/2.5, *v/v/v/v*) and the reaction was allowed to proceed at room temperature for 2 h. Subsequently, the supernatant was collected by filtration and concentrated under a light stream of N_2_. Thereafter, ice-cold diethyl ether (14 mL) was added to the concentrated supernatant, and the resulting precipitate collected as a pellet by centrifugation. The pellet was then suspended in a second portion of ice-cold diethyl ether (14 mL) and collected again by centrifugation. Residual ether was allowed to evaporate, and the peptide pellet was dissolved in 10–50% acetonitrile in water containing 0.1% TFA and lyophilized. Photocaged peptides were protected from light during all steps and handling.

#### 2.3 Semi-Preparative Reverse-Phase High-Performance Liquid Chromatography (RP-HPLC)

All peptides were purified by semi-preparative RP-HPLC on a Shimadzu prominence HPLC system (Shimadzu Corp., Japan), using an Agilent Zorbax 300SB-C18 or 300SB-C8 Semi-Preparative column (9.4 × 250 mm, 5 µm particle size), wherein eluent A = H_2_O with 0.1% TFA, and eluent B = MeCN with 0.1% TFA, and with UV absorbance detection at λ = 214 nm. Purifications were carried out at room temperature with a flow rate of 4.0 mL/min or 3.5 mL/min using linear gradients of eluents A and B as specified. Fractions (automatically collected) were then analyzed for purity by LCMS and UHPLC, and fractions containing the desired product (≥95% purity) were pooled and lyophilized.

#### 2.4 Yield Calculations

For synthesis of peptides herein, a pre-loaded Novabiochem® NovaPEG Rink Amide resin (0.41 mmol/g or 0.20 mmol/g) was used. Theoretical yield was determined based on weight of the resin, resin loading, and the molecular weight of each purified peptide.

#### 2.5 UV-Vis Spectroscopic Analysis of Peptides and PC Building Block

Individual UV-Vis spectra of Photo-GLP1, WT GLP-1, and α-methyl-5-methoxy-2-nitro-4-(2-propyn-1- yloxy)benzyl alcohol (PC Building Block) (80 µM in HBSS buffer) were acquired in the range of 200–450 nm with 1 nm steps. Absorbance values for each compound were plotted against wavelength.

#### 2.6 Photocleavage and Characterization of Photocleaved Photo-GLP1

Photo-GLP1 (500 µL, 80 µM in HBSS buffer) was irradiated with a UVA-LED (λ = 370 nm, Kessil PR160L-370 nm) at maximal power settings (0.64 mW/mm^2^) under air cooling. Samples (50 µL) were taken from the reaction at the following time-points: 20, 40, 120, and 300 seconds, and analyzed by LCMS. Another sample (50 µL) was taken at 300 seconds and combined with WT GLP1 (50 µL, 80 µM in HBSS buffer) and analyzed by LCMS to confirm identity of the photocleaved product.

### 3 Building Block Synthesis

#### 1-(5-methoxy-2-nitro-4-(prop-2-yn-1-yloxy)phenyl)ethyl *N*-succinimidyl carbonate (SI1)^[2]^

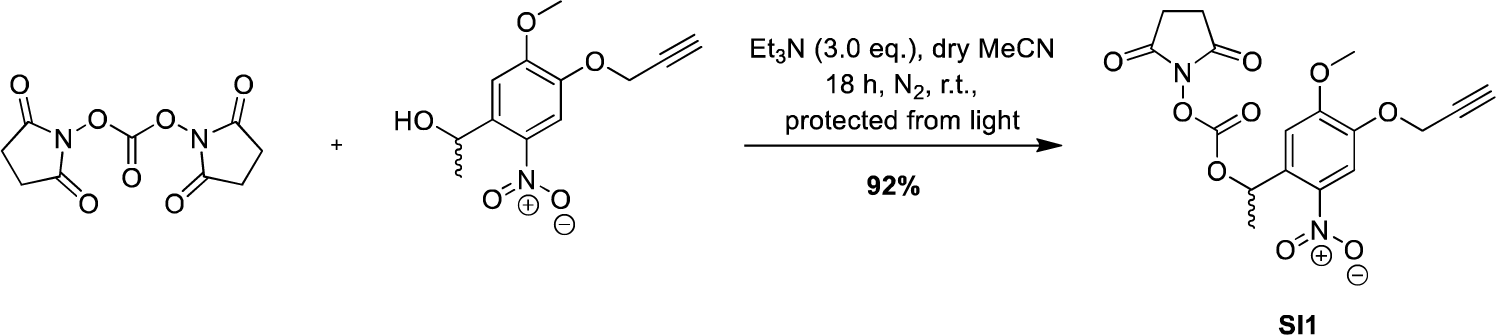

To a solution of dry acetonitrile (50 mL) was added α-methyl-5-methoxy-2-nitro-4-(2-propyn-1-yloxy)benzyl alcohol (0.10 g, 0.39 mmol, 1.0 eq.), *N,N*’-disuccinimidyl carbonate (0.23 g, 0.90 mmol, 2.3 eq.), and Et_3_N (0.16 mL, 1.2 mmol, 3.0 eq.). The reaction was protected from light stirred under N_2_ atmosphere at r.t. for 18 h. The reaction mixture was then concentrated under reduced pressure, and the product was isolated by flash column (3:1 followed by 2:1 hexanes:EtOAc) to yield the *title compound* **SI1** as an off-white solid (0.14 g, 92%). **Rf** 0.31 (1:1 hexanes:EtOAc). **IR** (film, CHCl_3_) 3282.9, 1812.7, 1788.7, 1739.9, 1583.6, 1521.9, 1456.9, 1376.2, 1336.9, 1278.3, 1207.7, 1077.7, 1045.8, 1014.8. **HRMS** (ESI) (m/z): [M+Na]^+^ calcd. for C_17_H_16_N_2_NaO_9_, 415.0748; found 415.0755. **^1^H NMR** (400 MHz, CDCl_3_) δ 7.83 (s, 1H), 7.11 (s, 1H), 6.52 (q, *J* = 6.4 Hz, 1H), 4.83 (dd, *J* = 2.5, 1.4 Hz, 2H), 4.06 (s, 3H), 2.80 (s, 4H), 2.59 (t, *J* = 2.4 Hz, 1H), 1.77 (d, *J* = 6.4 Hz, 3H). **^13^C NMR** (126 MHz, CDCl_3_) δ 168.58, 154.90, 150.73, 146.18, 139.17, 132.36, 110.62, 107.73, 57.15, 56.77, 25.56, 22.10.

Spectroscopic data was in good agreement with the literature.^[3]^

**^1^H NMR (400 MHz, CDCl_3_)** 1-(5-methoxy-2-nitro-4-(prop-2-yn-1-yloxy)phenyl)ethyl *N*-succinimidyl carbonate (**SI1**)

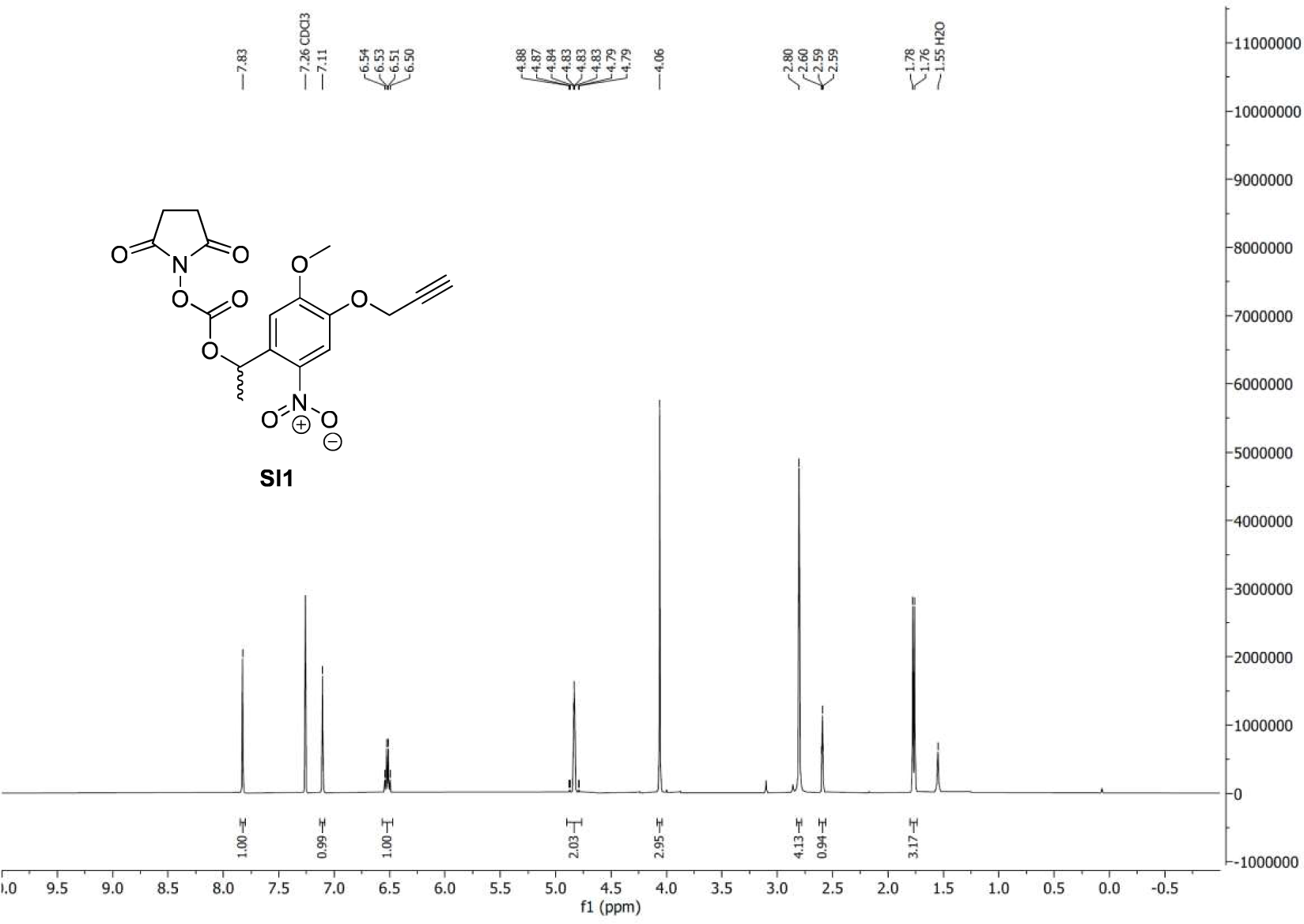

**^13^C NMR (126 MHz, CDCl_3_)** 1-(5-methoxy-2-nitro-4-(prop-2-yn-1-yloxy)phenyl)ethyl *N*-succinimidyl carbonate (**SI1**)

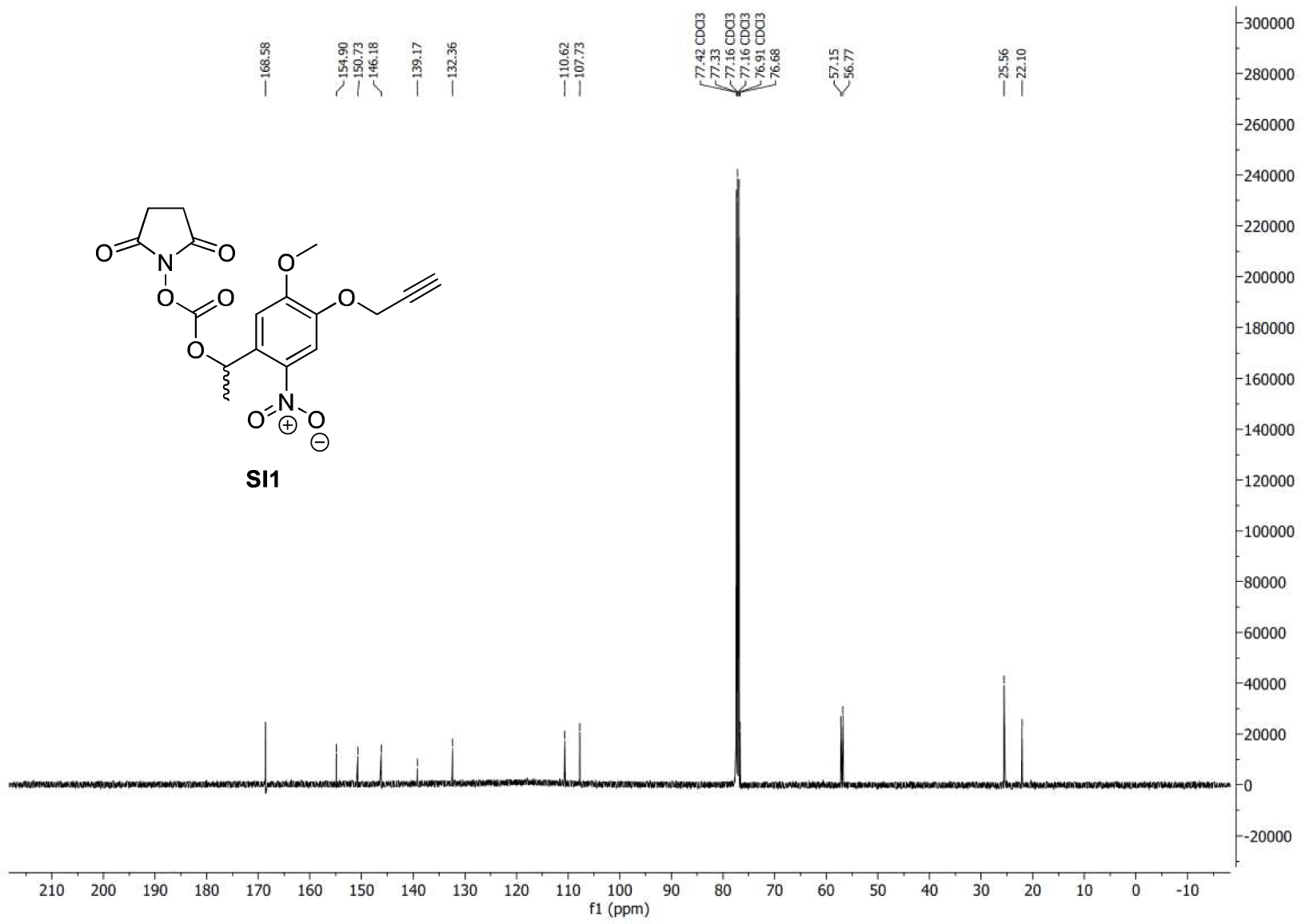

### 4 Peptide Synthesis

#### Synthesis of GLP1[6-30]

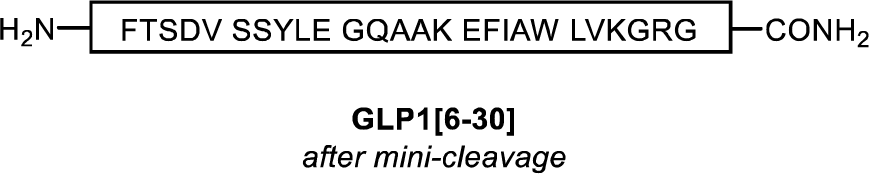

The peptide GLP1[6-30] was prepared via AFPS using Novabiochem® NovaPEG Rink Amide resin (0.41 mmol/g, 0.15 g, 62 µmol) using methods outlined in **General Procedure 2.1**. Total synthesis time to afford resin-bound GLP-1 was approximately 1.5 h. Mini-cleavage of the peptidyl-resin (5.0 mg) was carried out using **General Procedure 2.2** to confirm presence of the crude peptide (67% purity by LCMS, monoisotopic mass calc. 2800.4548, found 2800.7377), **SI Figure 1**. The remaining peptidyl-resin was divided into four equal portions (approx. 16 µmol each) for the synthesis of alanine scan analogues of GLP-1; [H1A]GLP1, [E3A]GLP1, [G4A]GLP1 and [T5A]GLP1.

**SI Figure 1.**
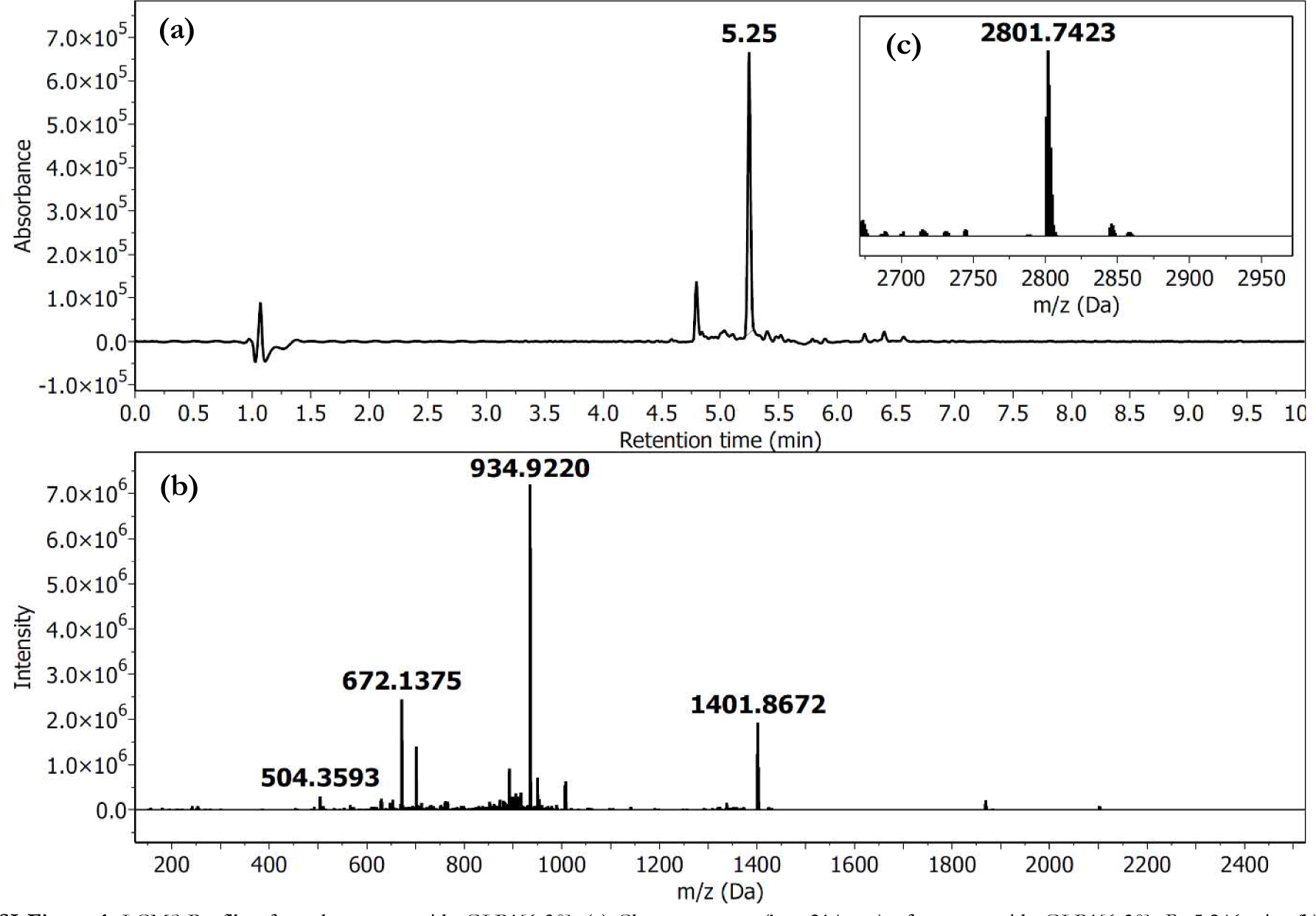
LCMS Profile of crude core peptide GLP1[6-30]. (**a**) Chromatogram (λ = 214 nm) of core peptide GLP1[6-30]; *R*t 5.246 min. (**b**) HRMS (ESI-TOF) spectrum. (**c**) Deconvoluted HRMS; monoisotopic mass calcd. for C_129_H_197_N_33_O_37_ 2800.4548, found 2800.7377.

#### Synthesis of [H1A]GLP1

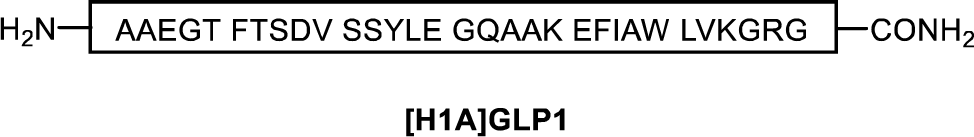

The peptide [H1A]GLP1 was prepared from resin-bound GLP1[6-30] (approx. 16 µmol) via AFPS using methods outlined in **General Procedure 2.1**. Total synthesis time to afford resin-bound [H1A]GLP1 (94 mg final resin weight) from GLP1[6-30] was approximately 20 min. Cleavage of the peptidyl-resin (50 mg) was carried out using **General Procedure 2.2** to afford the crude peptide (34 mg, 58% purity by UHPLC, monoisotopic mass calc. 3229.6408, found 3229.9870), **SI Figures 2 and 4**. The crude peptide (11 mg) was purified by semi-prep RP-HPLC using **General Procedure 2.3** with a gradient of 10–30% B over 20 min (*ca.* 1% B/min), followed by 30–50% B over 40 min (*ca.* 0.5% B/min) on an Agilent Zorbax 300SB-C18 column at room temperature. Fractions were analyzed by LCMS and UHPLC, and fractions containing the correct m/z and high purity (>95% purity by UHPLC) were combined and lyophilized to afford the *title compound* (2.6 mg, >95% purity by UHPLC, 25% overall yield, monoisotopic mass calc. 3229.6408, found 3229.9545), **SI Figures 3 and 5.**

**SI Figure 2.**
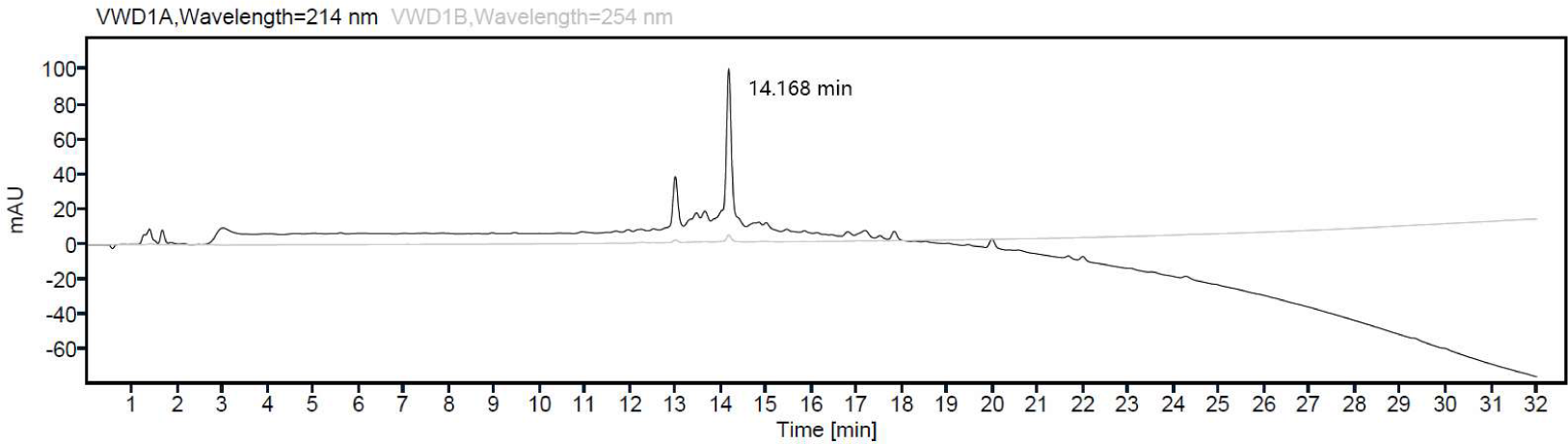
UHPLC profile of [H1A]GLP1 crude product. *R*t 14.168 min (Agilent Zorbax 300SB-C18 column, 5 µm, 2.1 × 150 mm, 5-95% MeCN over 30 min, *ca.* 3% MeCN/min), 58% purity based on Area Under Curve (AUC) at λ = 214 nm, accounting for all peaks between 4.0–29 min.

**SI Figure 3.**
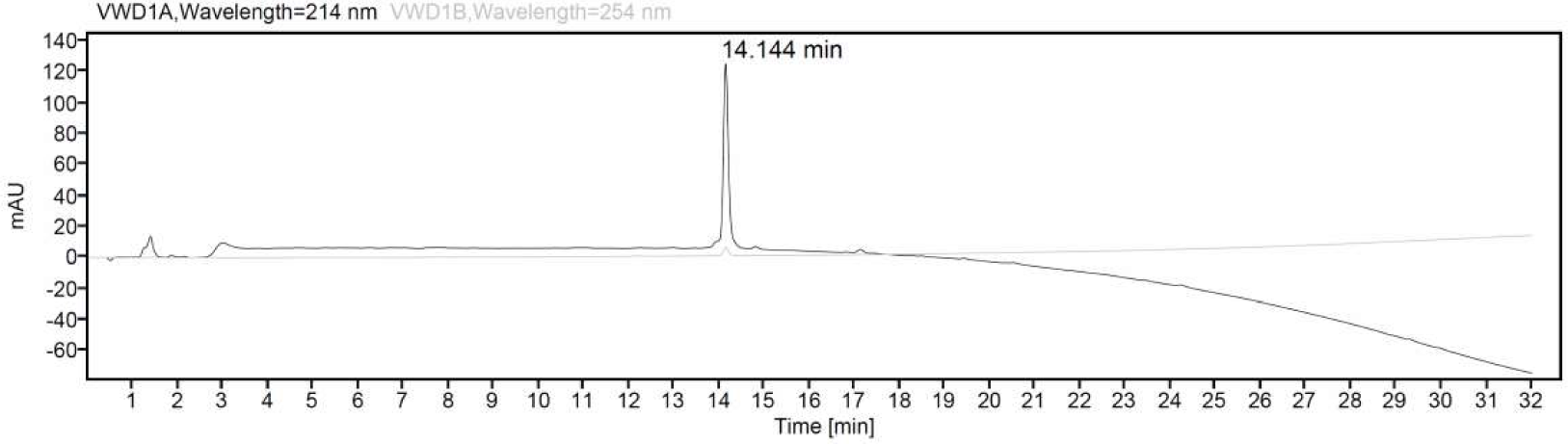
UHPLC profile of [H1A]GLP1 pure product. *R*t 14.144 min (Agilent Zorbax 300SB-C18 column, 5 µm, 2.1 × 150 mm, 5-95% MeCN over 30 min, *ca.* 3% MeCN/min), >95% purity based on Area Under Curve (AUC) at λ = 214 nm, accounting for all peaks between 4.0–29 min.

**SI Figure 4.**
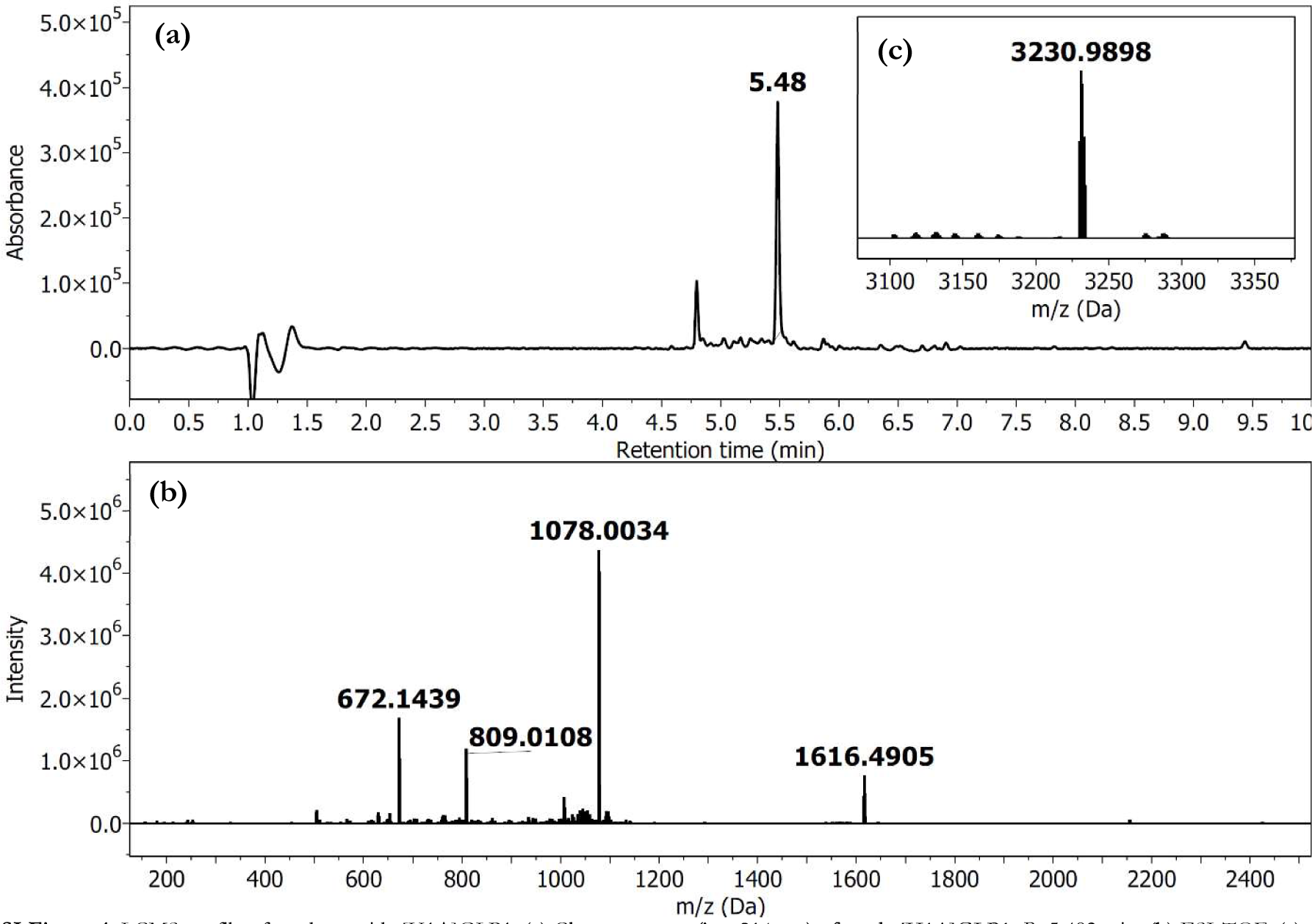
LCMS profile of crude peptide [H1A]GLP1. (**a**) Chromatogram (λ = 214 nm) of crude [H1A]GLP1; *R*t 5.482 min. (**b**) ESI-TOF. (**c**) Deconvoluted HRMS; monoisotopic mass calcd. for C_146_H_224_N_38_O_45_ 3229.6408, found 3229.9870.

**SI Figure 5.**
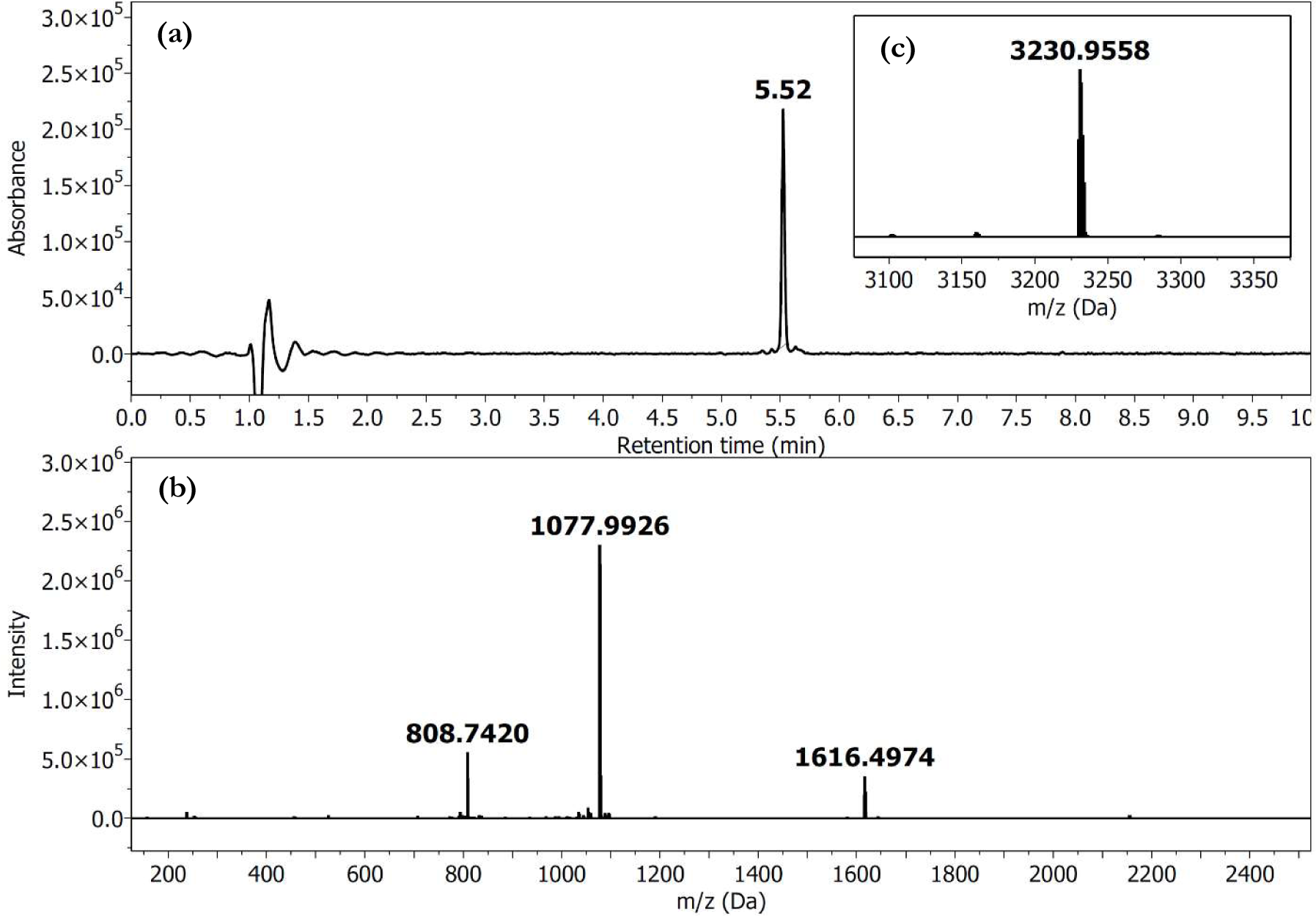
LCMS profile of pure peptide [H1A]GLP1. (**a**) Chromatogram (λ = 214 nm) of pure [H1A]GLP1; *R*t 5.523 min. (**b**) HRMS (ESI- TOF) spectrum. (**c**) Deconvoluted HRMS; monoisotopic mass calcd. for C_146_H_224_N_38_O_45_ 3229.6408, found 3229.9545.

#### Synthesis of [E3A]GLP1

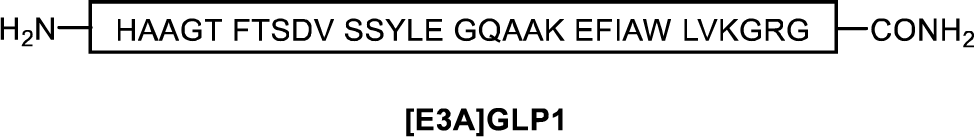

The peptide [E3A]GLP1 was prepared from resin-bound GLP1[6-30] (approx. 16 µmol) via AFPS using methods outlined in **General Procedure 2.1**. Total synthesis time to afford resin-bound [E3A]GLP1 (98 mg final resin weight) from GLP1[6-30] was approximately 20 min. Cleavage of the peptidyl-resin (52 mg) was carried out using **General Procedure 2.2** to afford the crude peptide (35 mg, 54% purity by UHPLC, monoisotopic mass calc. 3237.6571, found 3237.6608), **SI Figures 6 and 8**. The crude peptide (6.6 mg) was purified by semi-prep RP-HPLC using **General Procedure 2.3** with a gradient of 10–30% B over 20 min (*ca.* 1% B/min), followed by 30–50 % B over 40 min (*ca.* 0.5% B/min) on an Agilent Zorbax 300SB-C18 column at room temperature. Fractions were analyzed by LCMS and UHPLC, and fractions containing the correct m/z and high purity (>95% purity by UHPLC) were combined and lyophilized to afford the *title compound* (1.5 mg, >95% purity by UHPLC, 21% overall yield, monoisotopic mass calc. 3237.6571, found 3237.6512), **SI Figures 7 and 9.**

**SI Figure 6.**
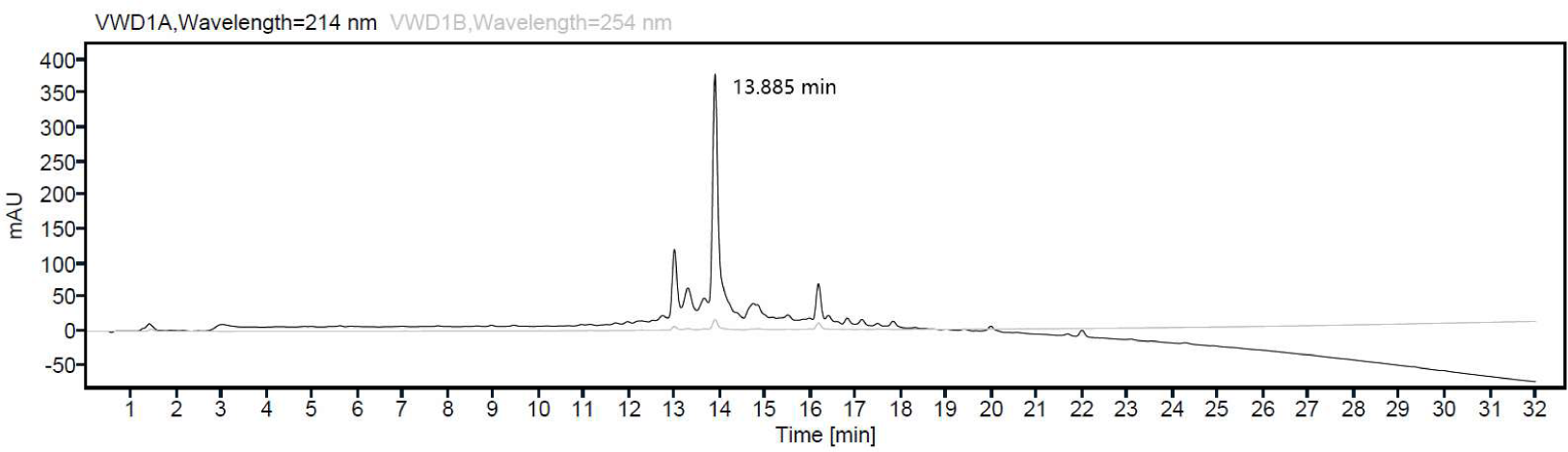
UHPLC profile of [E3A]GLP1 crude product. *R*t 13.885 min (Agilent Zorbax 300SB-C18 column, 5 µm, 2.1 × 150 mm, 5-95% MeCN over 30 min, *ca.* 3% MeCN/min), 54% purity based on Area Under Curve (AUC) at λ = 214 nm, accounting for all peaks between 4.0–29 min.

**SI Figure 7.**
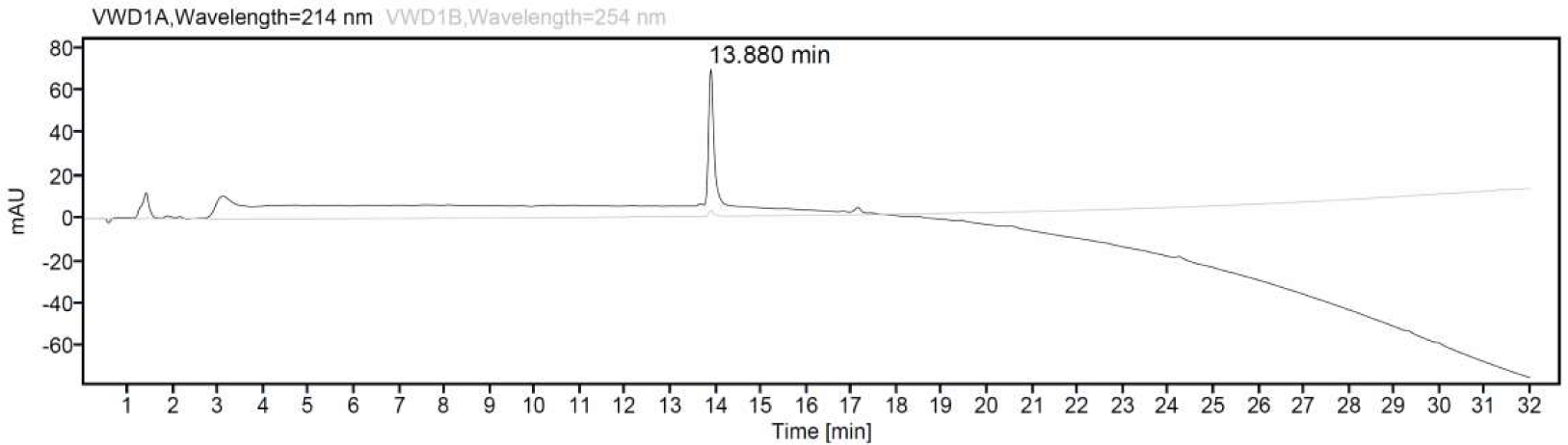
UHPLC profile of [E3A]GLP1 pure product. *R*t 13.880 min (Agilent Zorbax 300SB-C18 column, 5 µm, 2.1 × 150 mm, 5-95% MeCN over 30 min, *ca.* 3% MeCN/min), >95% purity based on Area Under Curve (AUC) at λ = 214 nm, accounting for all peaks between 4.0–29 min.

**SI Figure 8.**
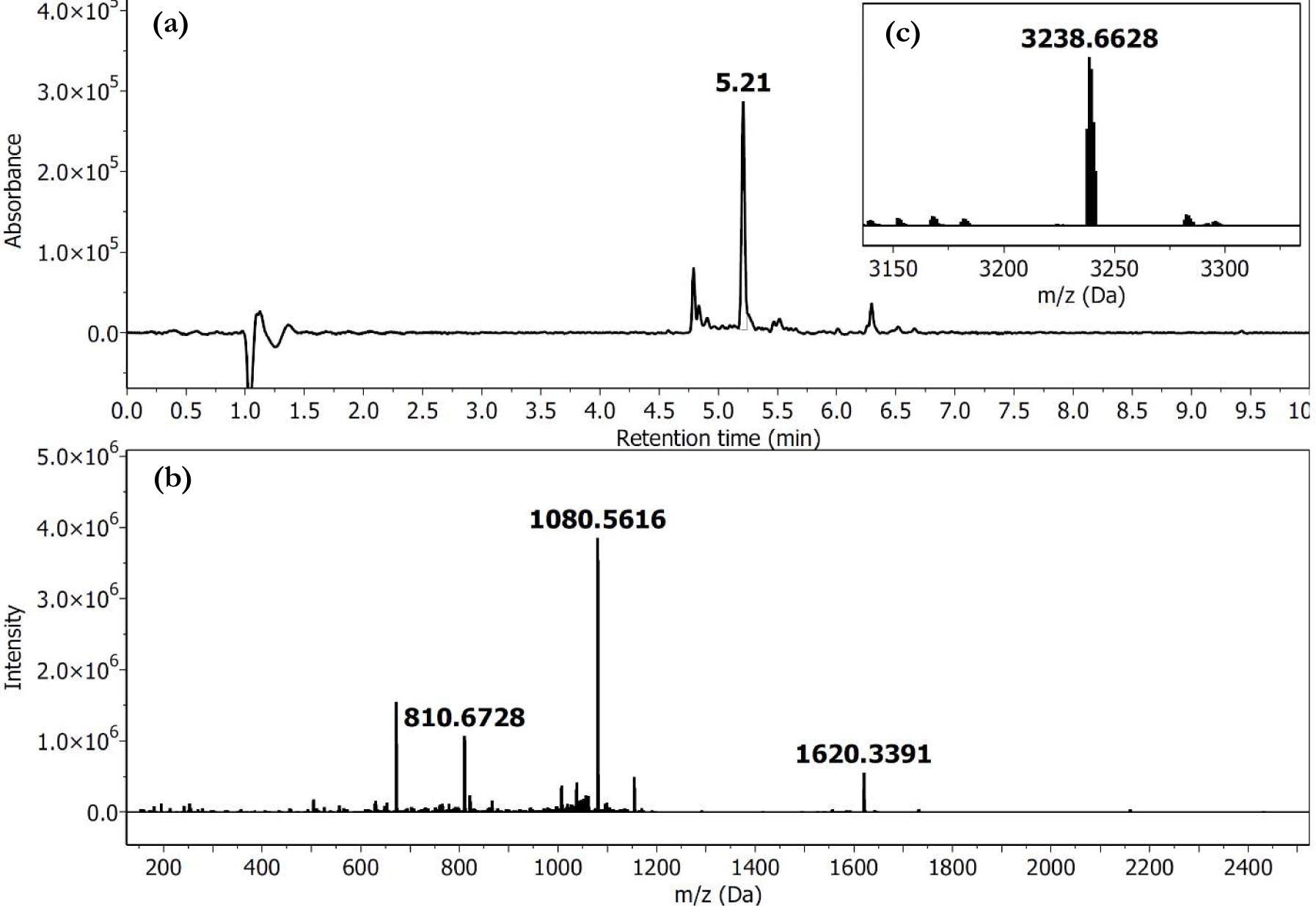
LCMS profile of crude peptide [E3A]GLP1. (**a**) Chromatogram (λ = 214 nm) of crude [E3A]GLP1; *R*t 5.210 min. (**b**) HRMS (ESI- TOF) spectrum. (**c**) Deconvoluted HRMS; monoisotopic mass calcd. for C_147_H_224_N_40_O_43_ 3237.6571, found 3237.6608.

**SI Figure 9.**
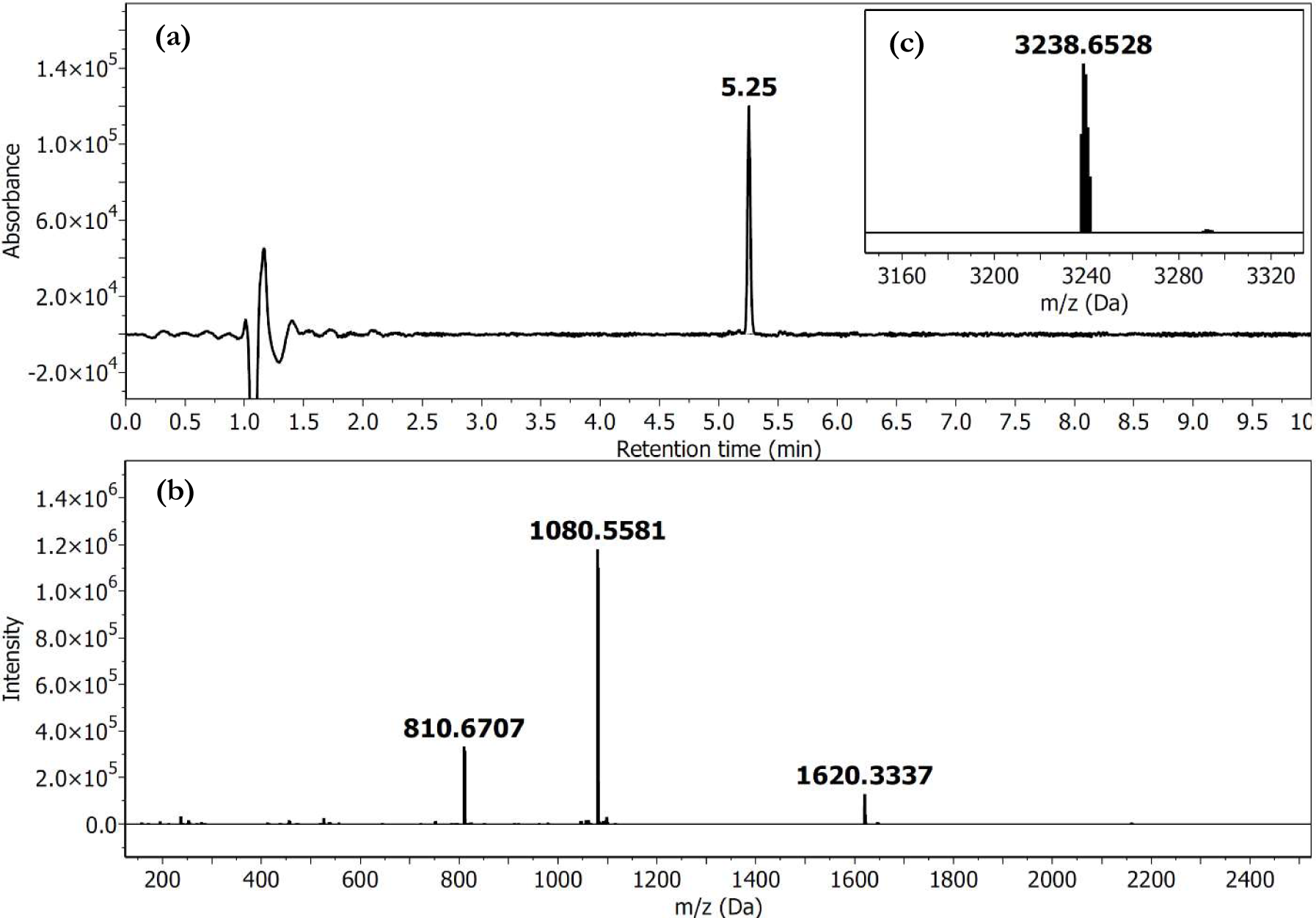
LCMS profile of pure peptide [E3A]GLP1. (**a**) Chromatogram (λ = 214 nm) of pure [E3A]GLP1; *R*t 5.253 min. (**b**) HRMS (ESI-TOF) spectrum. (**c**) Deconvoluted HRMS; monoisotopic mass calcd. for C_147_H_224_N_40_O_43_ 3237.6571, found 3237.6512.

#### Synthesis of [G4A]GLP1

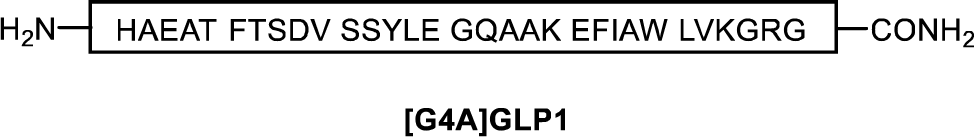

The peptide [G4A]GLP1 was prepared from resin-bound GLP1[6-30] (approx. 16 µmol) via AFPS using methods outlined in **General Procedure 2.1**. Total synthesis time to afford resin-bound [G4A]GLP1 (89 mg final resin weight) from GLP1[6-30] was approximately 20 min. Cleavage of the peptidyl-resin (47 mg) was carried out using **General Procedure 2.2** to afford the crude peptide (25 mg, 53% purity by UHPLC, monoisotopic mass calc. 3309.6782, found 3309.6778), **SI Figures 10 and 12.** The crude peptide (6.2 mg was purified by semi-prep RP-HPLC using **General Procedure 2.3** with a gradient of 10–30% B over 20 min (*ca.* 1% B/min), followed by 30–50% B over 40 min (*ca.* 0.5% B/min) on an Agilent Zorbax 300SB-C18 column at room temperature. Fractions were analyzed by LCMS and UHPLC, and fractions containing the correct m/z and high purity (>95% purity by UHPLC) were combined and lyophilized to afford the *title compound* (1.5 mg, 94% purity by UHPLC, 17% overall yield, monoisotopic mass calc. 3309.6782, found 3309.6696), **SI Figures 11 and 13**.

**SI Figure 10.**
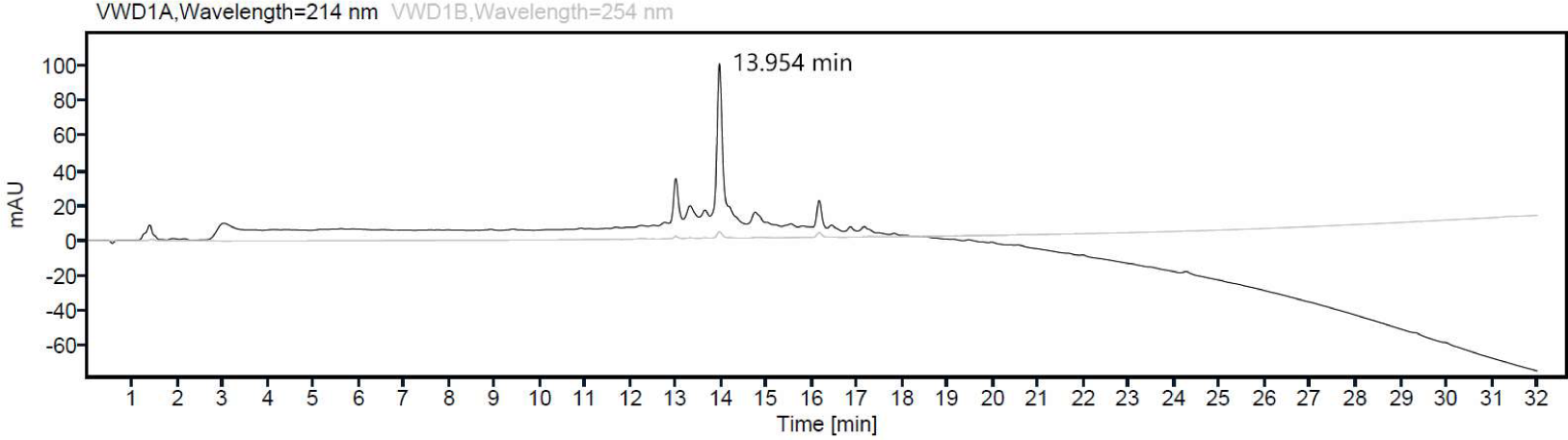
UHPLC profile of [G4A]GLP1 crude product. *R*t 13.954 min (Agilent Zorbax 300SB-C18 column, 5 µm, 2.1 × 150 mm, 5-95% MeCN over 30 min, *ca.* 3% MeCN/min), 53% purity based on Area Under Curve (AUC) at λ = 214 nm, accounting for all peaks between 4.0–29 min.

**SI Figure 11.**
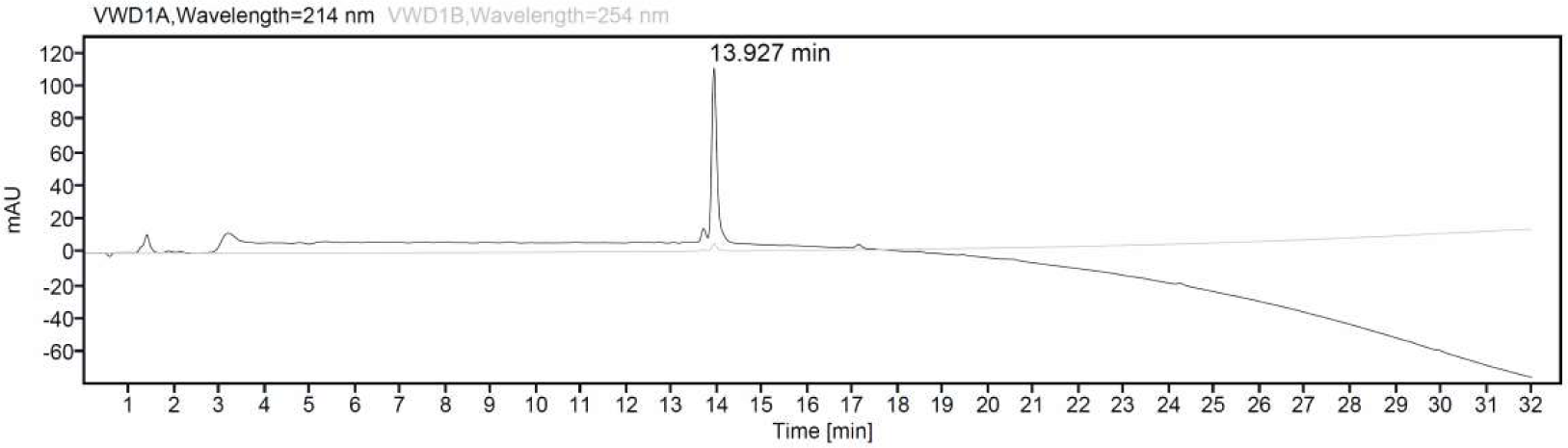
UHPLC profile of [G4A]GLP1 pure product. *R*t 13.927 min (Agilent Zorbax 300SB-C18 column, 5 µm, 2.1 × 150 mm, 5-95% MeCN over 30 min, *ca.* 3% MeCN/min), 94% purity based on Area Under Curve (AUC) at λ = 214 nm, accounting for all peaks between 4.0–29 min.

**SI Figure 12.**
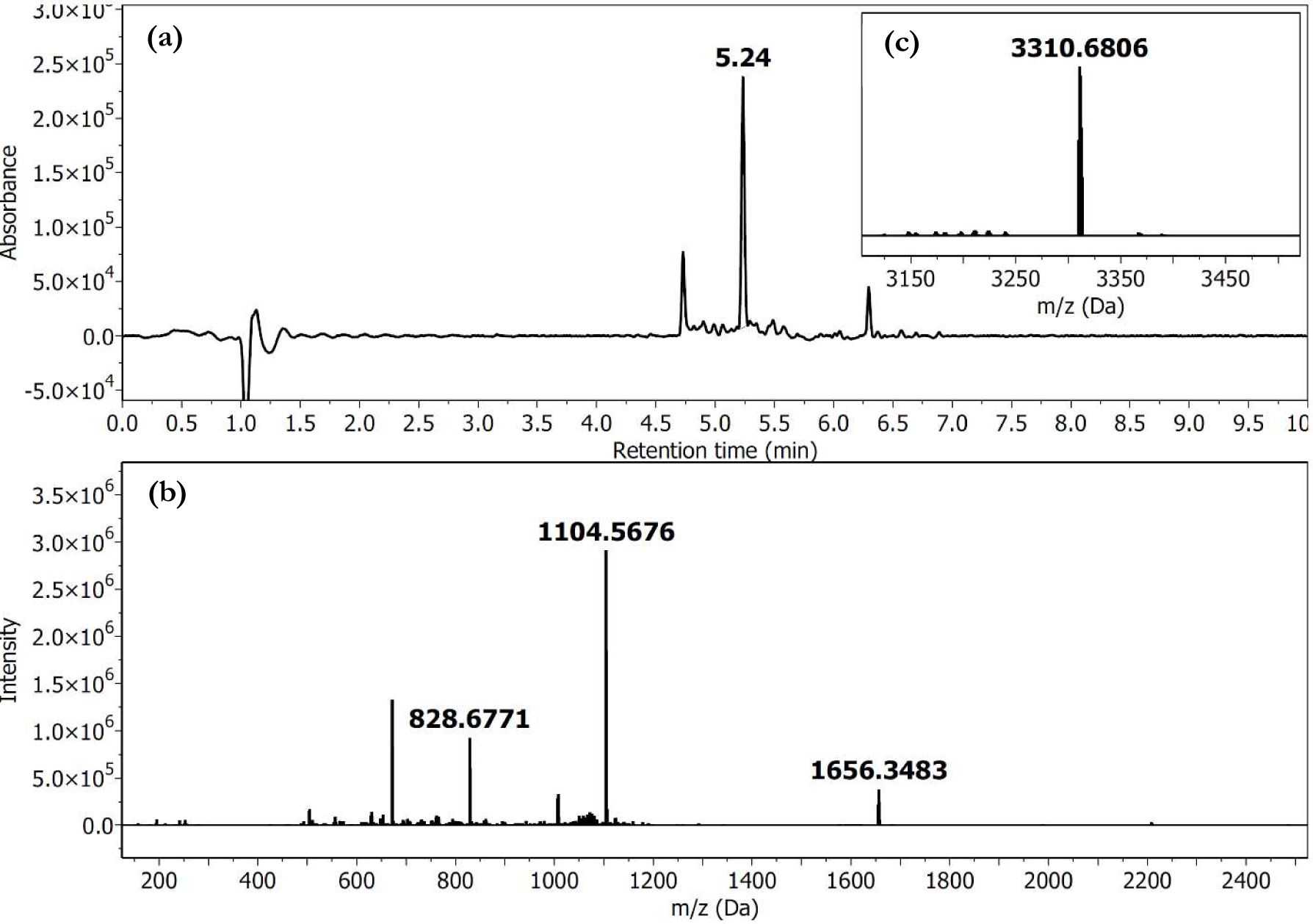
LCMS profile of crude peptide [G4A]GLP1. (**a**) Chromatogram (λ = 214 nm) of crude [G4A]GLP1; *R*t 5.235 min. (**b**) HRMS (ESI- TOF) spectrum. (**c**) Deconvoluted HRMS; monoisotopic mass (ESI+) calcd. for C_150_H_228_N_40_O_45_ 3309.6782, found 3309.6778.

**SI Figure 13.**
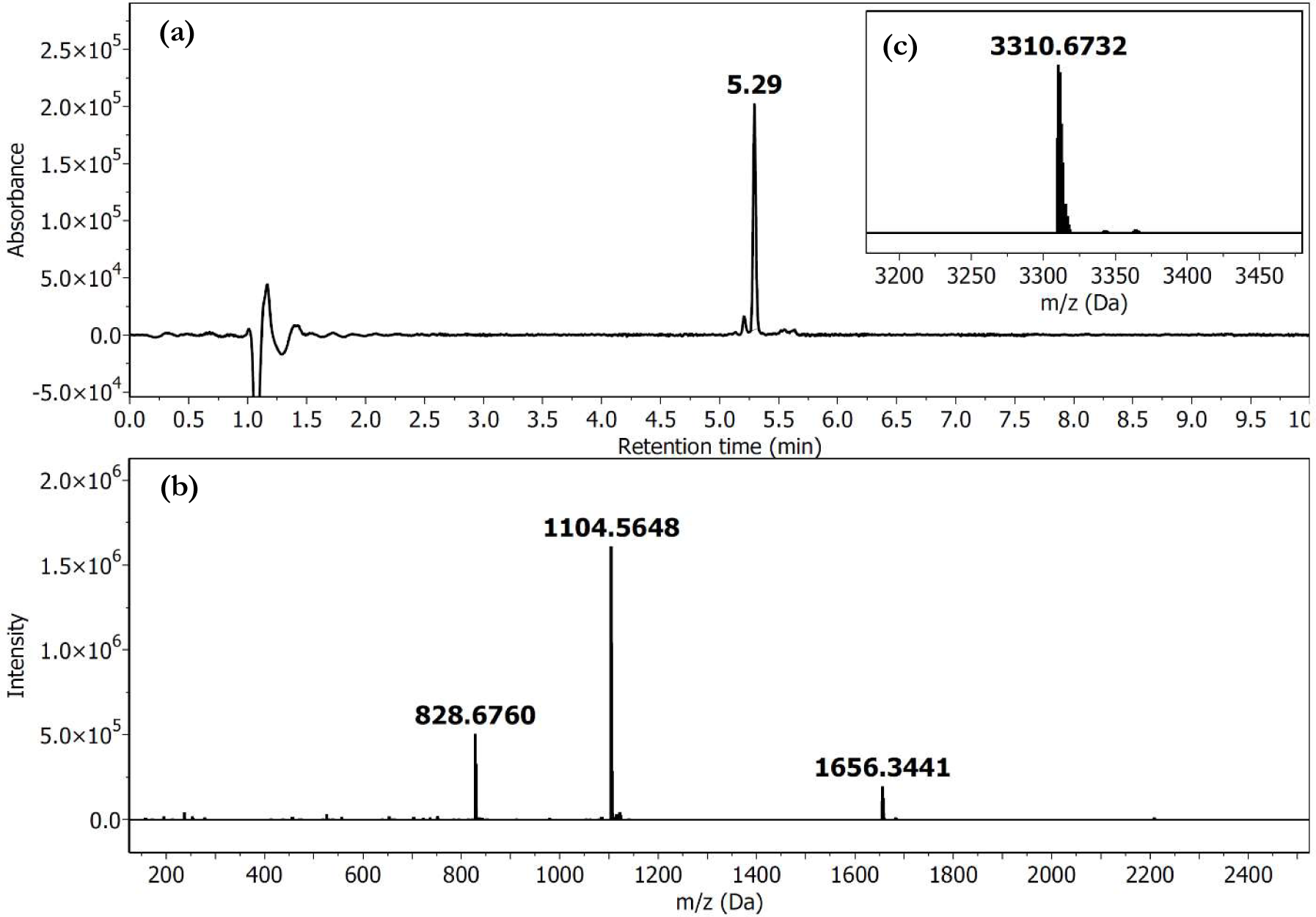
LCMS profile of pure peptide [G4A]GLP1. (**a**) Chromatogram (λ = 214 nm) of pure [G4A]GLP1; *R*t 5.294 min. (**b**) HRMS (ESI- TOF) spectrum. (**c**) Deconvoluted HRMS; monoisotopic mass (ESI+) calcd. for C_150_H_228_N_40_O_45_ 3309.6782, found 3309.6696.

#### Synthesis of [T5A]GLP1

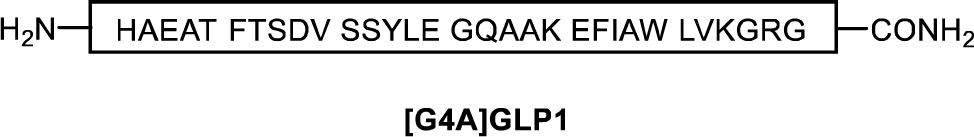

The peptide [T5A]GLP1 was prepared from resin-bound GLP1[6-30] (approx. 16 µmol) via AFPS using methods outlined in **General Procedure 2.1**. Total synthesis time to afford resin-bound [T5A]GLP1 (89 mg final resin weight) from GLP1[6-30] was approximately 20 min. Cleavage of the peptidyl-resin (49 mg) was carried out using **General Procedure 2.2** to afford the crude peptide (33 mg, 58% purity by UHPLC, monoisotopic mass calc. 3265.6520, found 3265.6516), **SI Figures 14 and 16**. The crude peptide (8.4 mg) was purified by semi-prep RP-HPLC using **General Procedure 2.3** with a gradient of 10–30% B over 20 min (*ca.* 1% B/min), followed by 30–50% B over 40 min (*ca.* 0.5% B/min) on an Agilent Zorbax 300SB-C18 column at room temperature. Fractions were analyzed by LCMS and UHPLC, and fractions containing the correct m/z and high purity (>95% purity by UHPLC) were combined and lyophilized to afford the *title compound* (0.84 mg, >95% purity by UHPLC, 9.3% overall yield, monoisotopic mass calc. 3265.6520, found 3265.6399), **SI Figures 15 and 17**.

**SI Figure 14.**
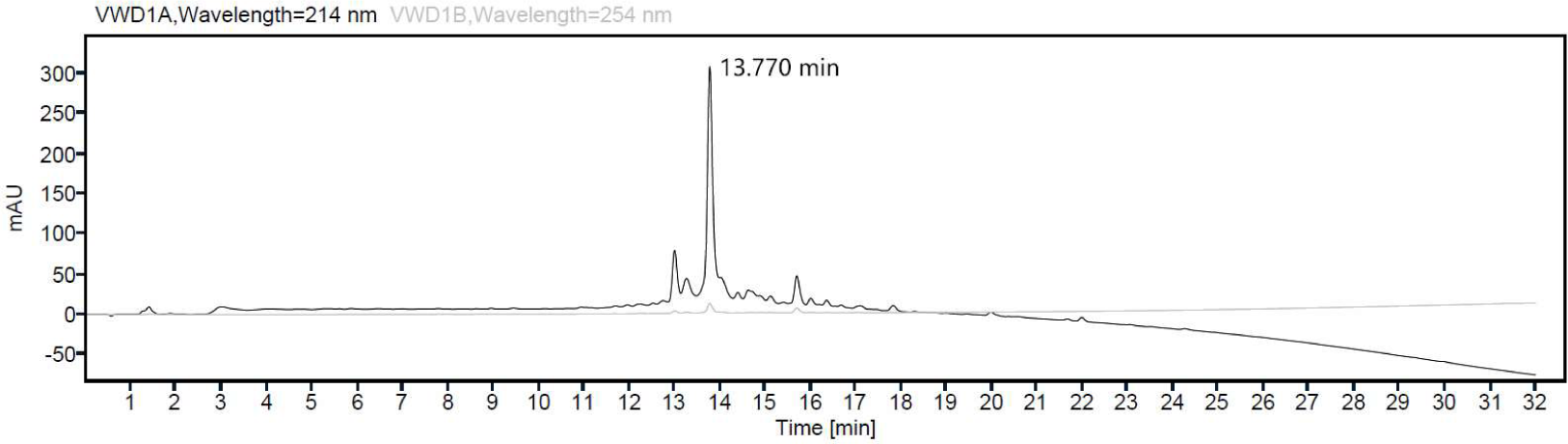
UHPLC profile of [T5A]GLP1 crude product. *R*t 13.770 min (Agilent Zorbax 300SB-C18 column, 5 µm, 2.1 × 150 mm, 5-95% MeCN over 30 min, *ca.* 3% MeCN/min), 58% purity based on Area Under Curve (AUC) at λ = 214 nm, accounting for all peaks between 4.0–29 min.

**SI Figure 15.**
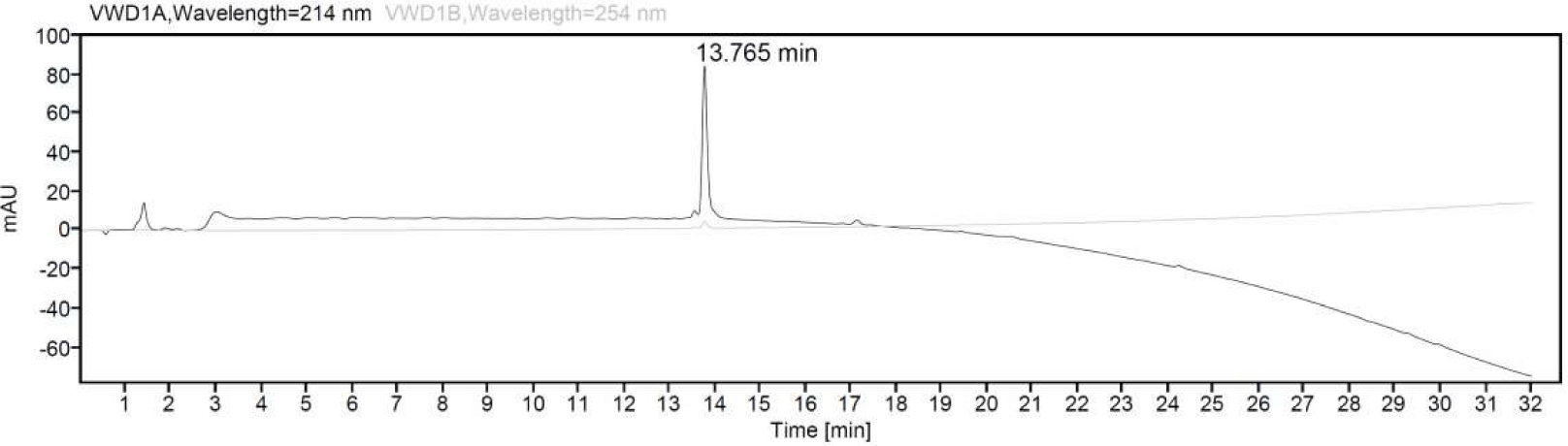
UHPLC profile of [T5A]GLP1 pure product. *R*t 13.765 min (Agilent Zorbax 300SB-C18 column, 5 µm, 2.1 × 150 mm, 5-95% MeCN over 30 min, *ca.* 3% MeCN/min), 95% purity based on Area Under Curve (AUC) at λ = 214 nm, accounting for all peaks between 4.0–29 min.

**SI Figure 16.**
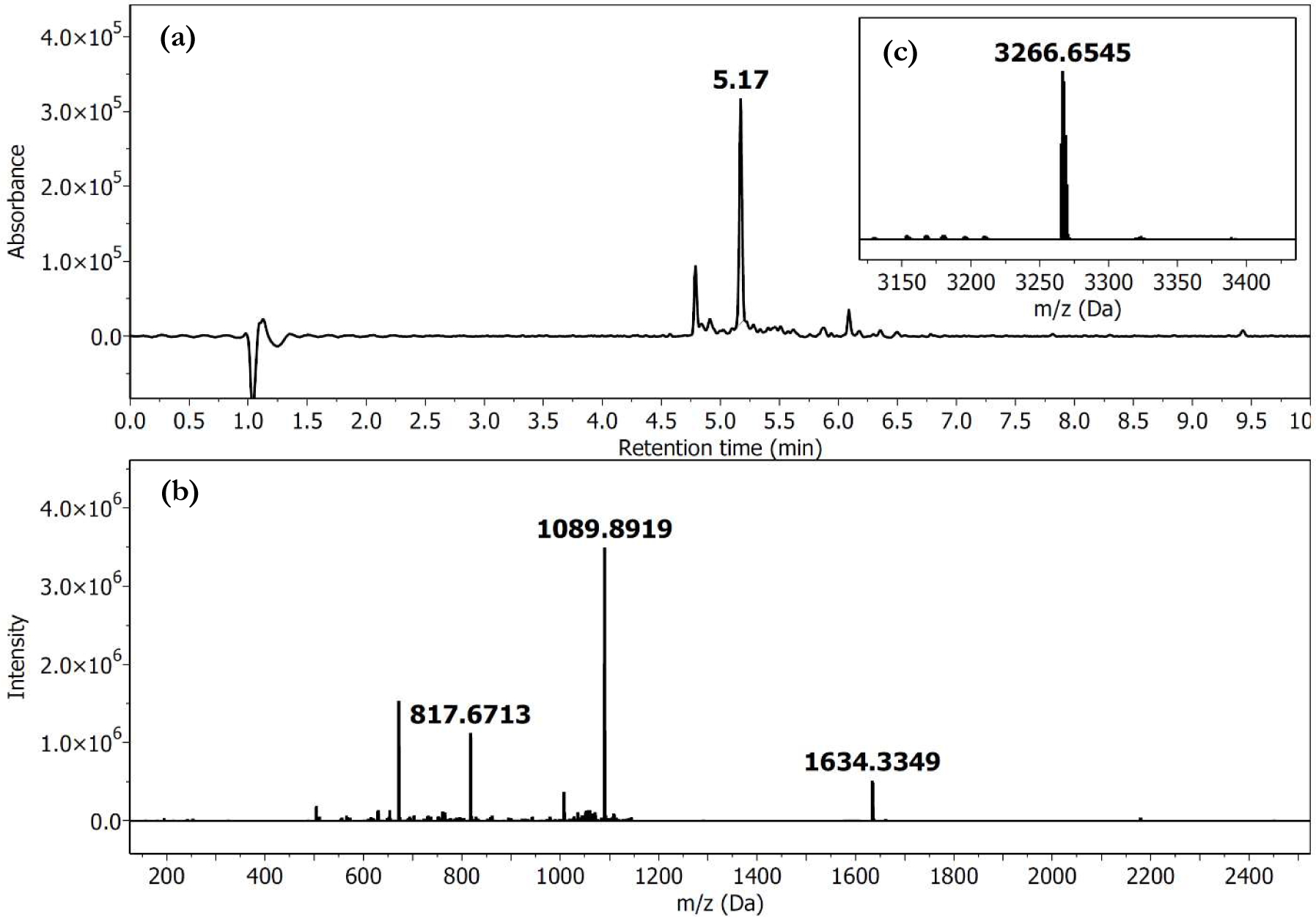
LCMS profile of crude peptide [T5A]GLP1. (**a**) Chromatogram (λ = 214 nm) of crude [T5A]GLP1; *R*t 5.172 min. (**b**) HRMS (ESI- TOF) spectrum. (**c**) Deconvoluted HRMS; monoisotopic mass (ESI+) calcd. for C_148_H_224_N_40_O_44_ 3265.6520, found 3265.6516.

**SI Figure 17.**
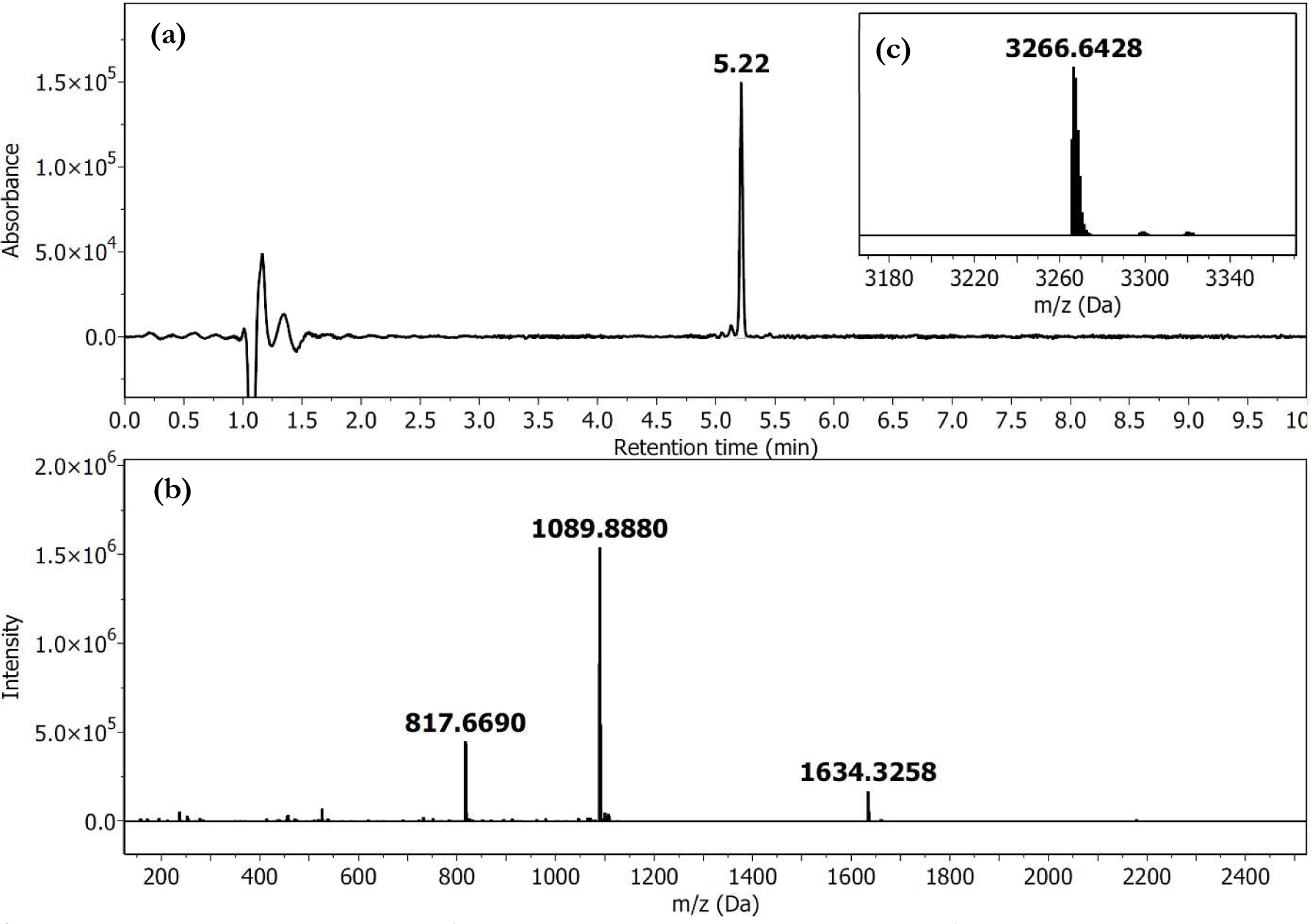
LCMS profile of pure peptide [T5A]GLP1. (**a**) Chromatogram (λ = 214 nm) of pure [T5A]GLP1; *R*t 5.215 min. (**b**) HRMS (ESI- TOF) spectrum. (**c**) Deconvoluted HRMS; monoisotopic mass (ESI+) calcd. for C_148_H_224_N_40_O_44_ 3265.6520, found 3265.6399.

#### Synthesis of Photo-GLP1

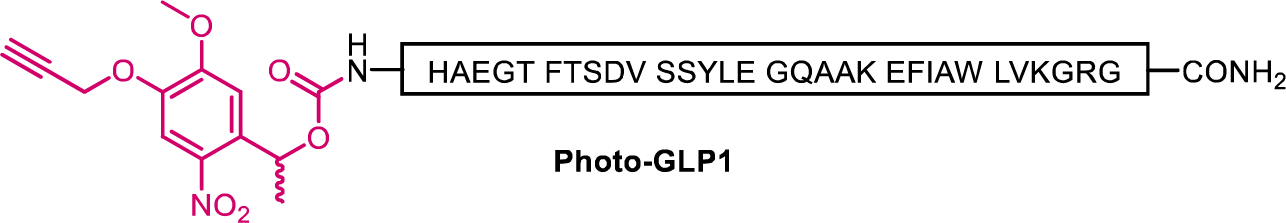

The peptide GLP-1 was first prepared via AFPS using Novabiochem® NovaPEG Rink Amide resin (0.20 mmol/g, 0.16 g, 32 µmol) using methods outlined in **General Procedure 2.1**. Total synthesis time to afford resin-bound GLP-1 (0.24 g final resin weight) was approximately 1.5 h. The peptidyl-resin (79 mg, approx. 10 µmol, 1.0 eq.) was swelled in DMF (2 mL) for 10 min, then drained. A solution of 1-(5-methoxy- 2-nitro-4-(prop-2-yn-1-yloxy)phenyl)ethyl *N*-succinimidyl carbonate (**SI1**) (39 mg, 0.10 mmol, 10 eq.) and *i*Pr_2_NEt (35 µL, 0.20 mmol, 20 eq.) in DMF (2 mL) was added to the pre-swelled peptidyl-resin, and the reaction was stirred at room temperature for 1 h. The peptidyl-resin was then drained, washed with DMF (3 × 2 mL) and CH_2_Cl_2_ (3 × 2 mL), then dried under reduced pressure to afford resin-bound Photo-GLP1 (68 mg final resin weight). Cleavage of the peptidyl-resin (32 mg) was carried out using **General Procedure 2.2** to afford the crude peptide (5.9 mg, 49% purity by UHPLC, monoisotopic mass calc. 3629.7427, found 3629.2054) **SI Figures 18 and 20.** The crude peptide (5.9 mg) was purified by semi-prep RP-HPLC using **General Procedure 2.3** with a gradient of 10–95% B over 170 min (*ca.* 0.5% B/min) on an Agilent Zorbax 300SB-C8 column at room temperature. Fractions were analyzed by LCMS and UHPLC, and fractions containing the correct m/z and high purity (>95% purity by UHPLC) were combined and lyophilized to afford the *title compound* (0.89 mg, >95% purity by UHPLC, 5% overall yield, monoisotopic mass calc. 3629.7427, found 3629.2373), **SI Figures 19 and 21**.

**SI Figure 18.**
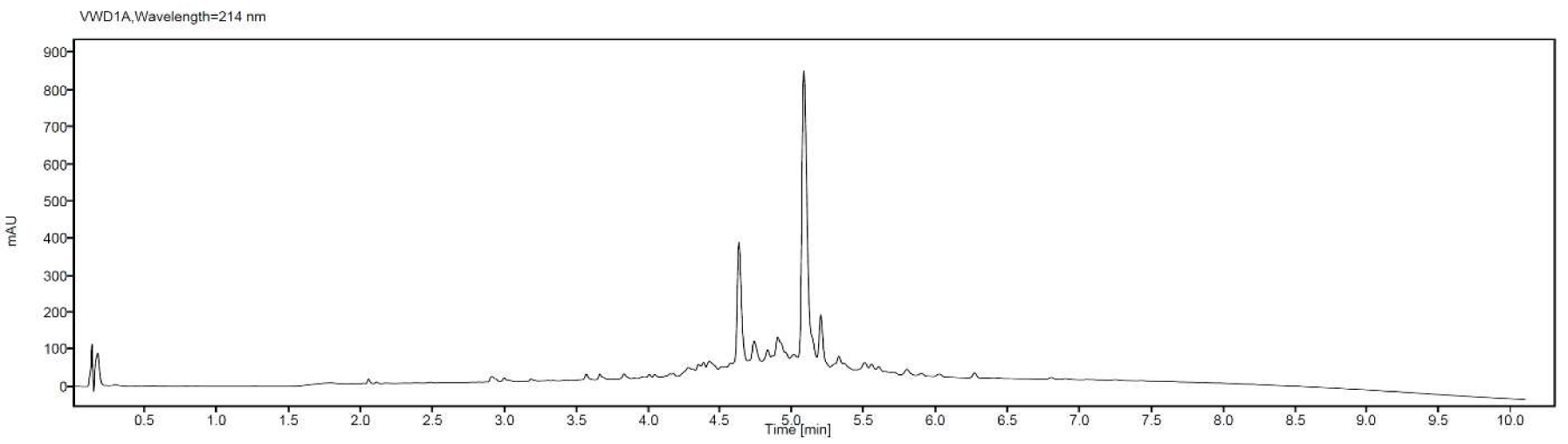
UHPLC profile of crude photo-GLP1; *R*t 5.081 min (Agilent Zorbax Eclipse Plus C18 Rapid Resolution HD column, 1.8 µm, 2.1 × 50 mm, 5-95% MeCN over 9 min, *ca.* 10% MeCN/min), 49% purity based on Area Under Curve (AUC) at λ = 214 nm, accounting for all peaks between 2.0–9.5 min.

**SI Figure 19.**
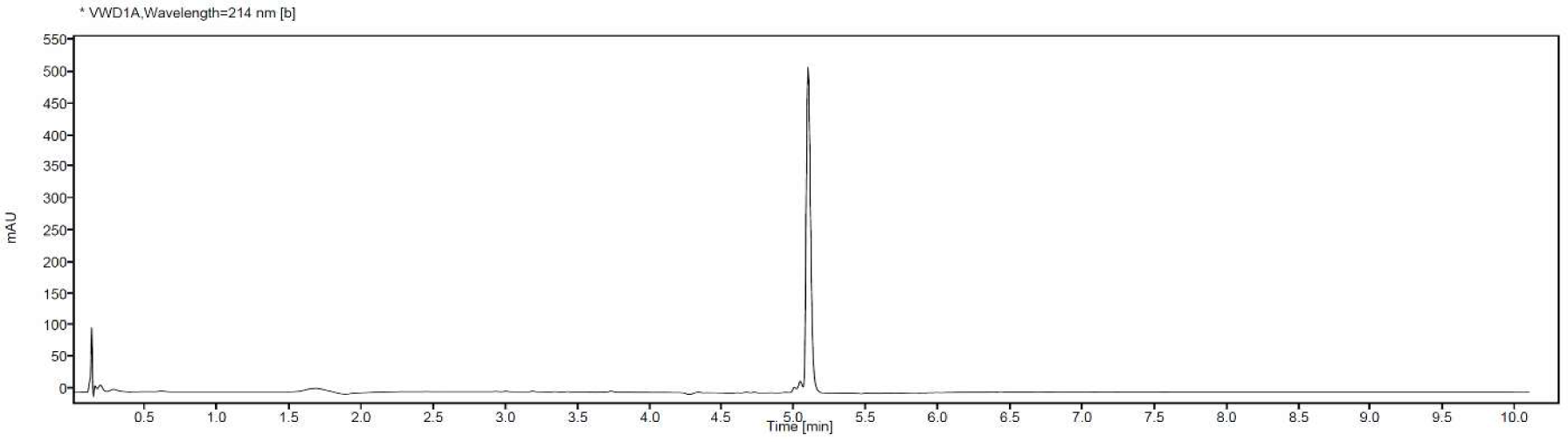
UHPLC profile of pure photo-GLP1; *R*t 5.097 min (Agilent Zorbax Eclipse Plus C18 Rapid Resolution HD column, 1.8 µm, 2.1 × 50 mm, 5-95% MeCN over 9 min, *ca.* 10% MeCN/min), >95% purity based on Area Under Curve (AUC) at λ = 214 nm, accounting for all peaks between 2.0–9.5 min

**SI Figure 20.**
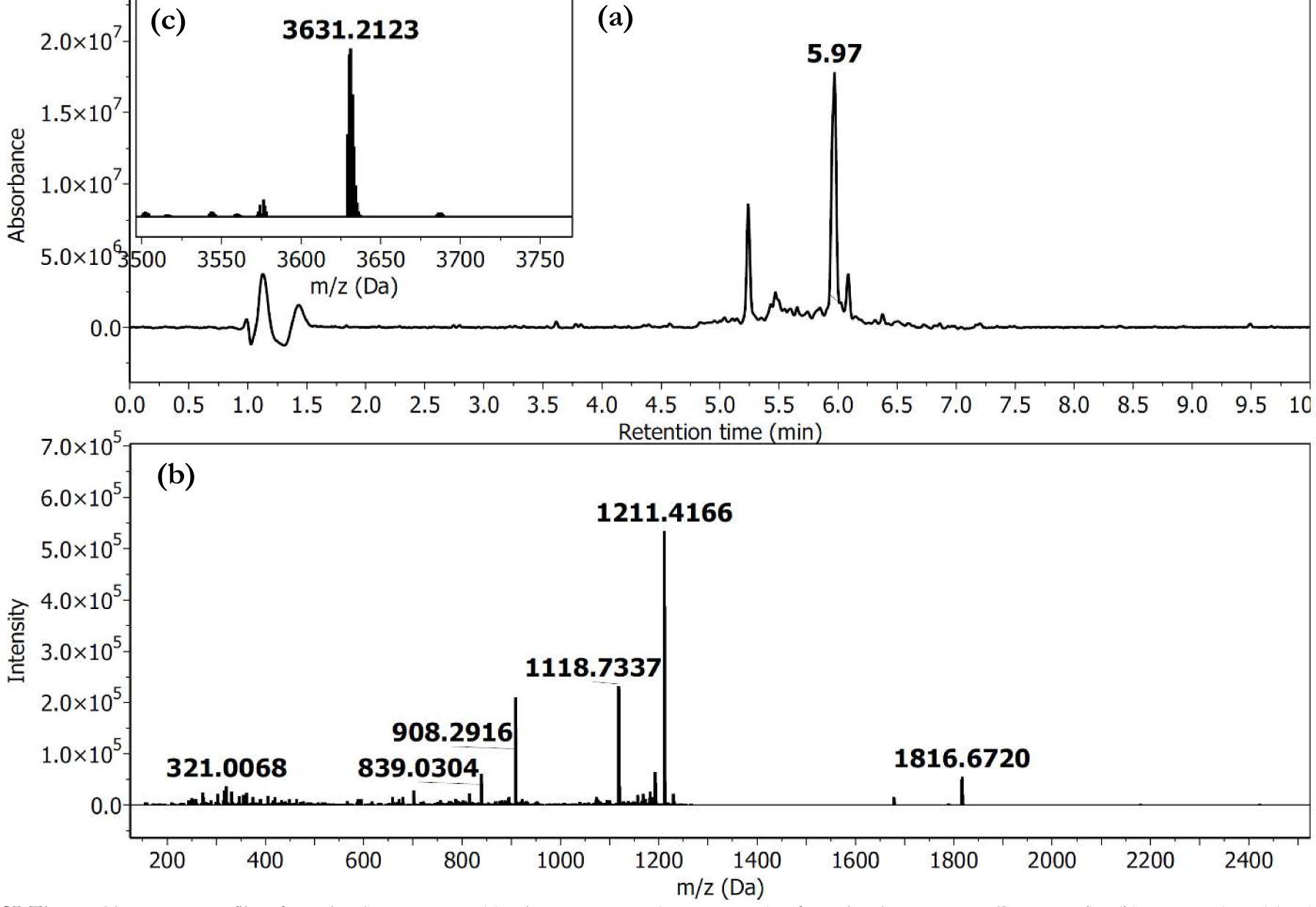
LCMS profile of crude photo-GLP1. (**a**) Chromatogram (λ = 214 nm) of crude photo-GLP1; *R*t 5.97 min. (**b**) HRMS (ESI-TOF) spectrum. (**c**) Deconvoluted HRMS; monoisotopic mass (ESI+) calcd. for C_164_H_240_N_42_O_52_ 3629.7427, found 3629. 2054.

**SI Figure 21.**
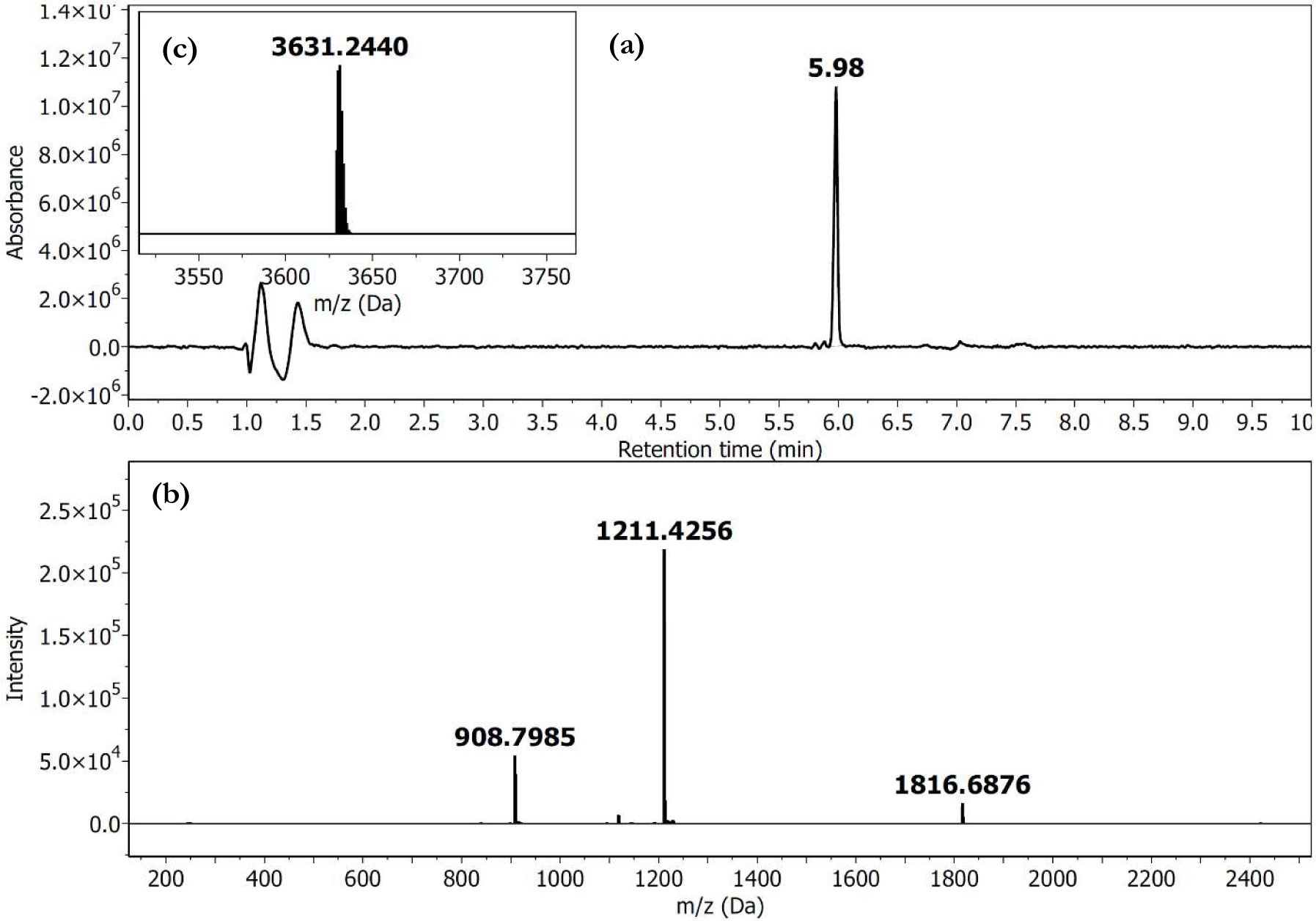
LCMS profile of pure photo-GLP1. (**a**) Chromatogram (λ = 214 nm) of pure photo-GLP1; *R*t 5.98 min. (**b**) HRMS (ESI-TOF) spectrum. (**c**) Deconvoluted HRMS; monoisotopic mass (ESI+) calcd. for C_164_H_240_N_42_O_52_ 3629.7427, found 3629.2373.

## References

Adelhorst K, Hedegaard BB, Knudsen LB, Kirk O. 1994. Structure-activity studies of glucagon- like peptide-1. J Biol Chem 269:6275–6278.

Alvarez E, Martínez MD, Roncero I, Chowen JA, García-Cuartero B, Gispert JD, Sanz C, Vázquez P, Maldonado A, de Cáceres J, Desco M, Pozo MA, Blázquez E. 2005. The expression of GLP-1 receptor mRNA and protein allows the effect of GLP-1 on glucose metabolism in the human hypothalamus and brainstem. J Neurochem 92:798–806. doi:10.1111/j.1471-4159.2004.02914.x

Amao M, Kitahara Y, Tokunaga A, Shimbo K, Eto Y, Yamada N. 2015. Simultaneous quantification of intracellular and secreted active and inactive glucagon-like peptide-1 from cultured cells. Anal Biochem 472:45–51. doi:10.1016/j.ab.2014.11.009

Andersen A, Lund A, Knop FK, Vilsbøll T. 2018. Glucagon-like peptide 1 in health and disease. Nat Rev Endocrinol 14:390–403. doi:10.1038/s41574-018-0016-2

Ast J, Arvaniti A, Fine NHF, Nasteska D, Ashford FB, Stamataki Z, Koszegi Z, Bacon A, Jones BJ, Lucey MA, Sasaki S, Brierley DI, Hastoy B, Tomas A, D’Agostino G, Reimann F, Lynn FC, Reissaus CA, Linnemann AK, D’Este E, Calebiro D, Trapp S, Johnsson K, Podewin T, Broichhagen J, Hodson DJ. 2020. Super-resolution microscopy compatible fluorescent probes reveal endogenous glucagon-like peptide-1 receptor distribution and dynamics. Nat Commun 11:467. doi:10.1038/s41467-020-14309-w

Biggs EK, Liang L, Naylor J, Madalli S, Collier R, Coghlan MP, Baker DJ, Hornigold DC, Ravn P, Reimann F, Gribble FM. 2018. Development and characterisation of a novel glucagon like peptide-1 receptor antibody. Diabetologia 61:711–721. doi:10.1007/s00125-017-4491-0

Broichhagen J, Podewin T, Meyer-Berg H, von Ohlen Y, Johnston NR, Jones BJ, Bloom SR, Rutter GA, Hoffmann-Röder A, Hodson DJ, Trauner D. 2015. Optical Control of Insulin Secretion Using an Incretin Switch. Angew Chem Int Ed Engl 54:15565–15569. doi:10.1002/anie.201506384

Brubaker PL, Schloos J, Drucker DJ. 1998. Regulation of glucagon-like peptide-1 synthesis and secretion in the GLUTag enteroendocrine cell line. Endocrinology 139:4108–4114. doi:10.1210/endo.139.10.6228

Campos RV, Lee YC, Drucker DJ. 1994. Divergent tissue-specific and developmental expression of receptors for glucagon and glucagon-like peptide-1 in the mouse. Endocrinology 134:2156–2164. doi:10.1210/endo.134.5.8156917

Cannaert A, Storme J, Franz F, Auwärter V, Stove CP. 2016. Detection and Activity Profiling of Synthetic Cannabinoids and Their Metabolites with a Newly Developed Bioassay. Anal Chem 88:11476–11485. doi:10.1021/acs.analchem.6b02600

Cardin JA, Crair MC, Higley MJ. 2020. Mesoscopic Imaging: Shining a Wide Light on Large- Scale Neural Dynamics. Neuron 108:33–43. doi:10.1016/j.neuron.2020.09.031

Chen Z, Truskinovsky L, Tzanakakis ES. 2022. Emerging molecular technologies for light- mediated modulation of pancreatic beta-cell function. Mol Metab 64:101552. doi:10.1016/j.molmet.2022.101552

Cork SC, Richards JE, Holt MK, Gribble FM, Reimann F, Trapp S. 2015. Distribution and characterisation of Glucagon-like peptide-1 receptor expressing cells in the mouse brain. Mol Metab 4:718–731. doi:10.1016/j.molmet.2015.07.008

Dixon AS, Schwinn MK, Hall MP, Zimmerman K, Otto P, Lubben TH, Butler BL, Binkowski BF, Machleidt T, Kirkland TA, Wood MG, Eggers CT, Encell LP, Wood KV. 2016. NanoLuc Complementation Reporter Optimized for Accurate Measurement of Protein Interactions in Cells. ACS Chem Biol 11:400–408. doi:10.1021/acschembio.5b00753

Drucker DJ. 2001. Minireview: The Glucagon-Like Peptides. Endocrinology 142:521–527. doi:10.1210/endo.142.2.7983

Duffet L, Kosar S, Panniello M, Viberti B, Bracey E, Zych AD, Radoux-Mergault A, Zhou X, Dernic J, Ravotto L, Tsai Y-C, Figueiredo M, Tyagarajan SK, Weber B, Stoeber M, Gogolla N, Schmidt MH, Adamantidis AR, Fellin T, Burdakov D, Patriarchi T. 2022a. A genetically encoded sensor for in vivo imaging of orexin neuropeptides. Nat Methods 19:231–241. doi:10.1038/s41592-021-01390-2

Duffet L, Tatarskiy PV, Harada M, Williams ET, Hartrampf N, Patriarchi T. 2022b. A photocaged orexin-B for spatiotemporally precise control of orexin signaling. Cell Chemical Biology 0. doi:10.1016/j.chembiol.2022.11.007

Feng J, Zhang C, Lischinsky JE, Jing M, Zhou J, Wang H, Zhang Y, Dong A, Wu Z, Wu H, Chen W, Zhang P, Zou J, Hires SA, Zhu JJ, Cui G, Lin D, Du J, Li Y. 2019. A Genetically Encoded Fluorescent Sensor for Rapid and Specific In Vivo Detection of Norepinephrine. Neuron 102:745–761.e8. doi:10.1016/j.neuron.2019.02.037

Frank JA, Broichhagen J, Yushchenko DA, Trauner D, Schultz C, Hodson DJ. 2018. Optical tools for understanding the complexity of β-cell signalling and insulin release. Nat Rev Endocrinol 14:721–737. doi:10.1038/s41574-018-0105-2

Gibson DG, Young L, Chuang R-Y, Venter JC, Hutchison CA, Smith HO. 2009. Enzymatic assembly of DNA molecules up to several hundred kilobases. Nat Methods 6:343–345. doi:10.1038/nmeth.1318

Gunaydin LA, Grosenick L, Finkelstein JC, Kauvar IV, Fenno LE, Adhikari A, Lammel S, Mirzabekov JJ, Airan RD, Zalocusky KA, Tye KM, Anikeeva P, Malenka RC, Deisseroth K. 2014. Natural neural projection dynamics underlying social behavior. Cell 157:1535 doi:10.1016/j.cell.2014.05.017

Hartrampf N, Saebi A, Poskus M, Gates ZP, Callahan AJ, Cowfer AE, Hanna S, Antilla S, Schissel CK, Quartararo AJ, Ye X, Mijalis AJ, Simon MD, Loas A, Liu S, Jessen C, Nielsen TE, Pentelute BL. 2020. Synthesis of proteins by automated flow chemistry. Science 368:980–987. doi:10.1126/science.abb2491

Helmchen F. 2009. Two-Photon Functional Imaging of Neuronal Activity In: Frostig RD, editor. In Vivo Optical Imaging of Brain Function, Frontiers in Neuroscience. Boca Raton (FL): CRC Press/Taylor & Francis.

Hölscher C. 2022. Protective properties of GLP-1 and associated peptide hormones in neurodegenerative disorders. British Journal of Pharmacology 179:695–714. doi:10.1111/bph.15508

Holst JJ, Vilsbøll T, Deacon CF. 2009. The incretin system and its role in type 2 diabetes mellitus. Mol Cell Endocrinol 297:127–136. doi:10.1016/j.mce.2008.08.012

Holz GG, Chepurny OG, Leech CA, Song W-J, Hussain MA. 2015. Molecular Basis of cAMP Signaling in Pancreatic β Cells In: Islam MdS, editor. Islets of Langerhans. Dordrecht: Springer Netherlands. pp. 565–603. doi:10.1007/978-94-007-6686-0_25

Ino D, Tanaka Y, Hibino H, Nishiyama M. 2022. A fluorescent sensor for real-time measurement of extracellular oxytocin dynamics in the brain. Nat Methods 1–9. doi:10.1038/s41592-022-01597-x

Jazayeri A, Rappas M, Brown AJH, Kean J, Errey JC, Robertson NJ, Fiez-Vandal C, Andrews SP, Congreve M, Bortolato A, Mason JS, Baig AH, Teobald I, Doré AS, Weir M, Cooke RM, Marshall FH. 2017. Crystal structure of the GLP-1 receptor bound to a peptide agonist. Nature 546:254–258. doi:10.1038/nature22800

Knudsen LB, Kiel D, Teng M, Behrens C, Bhumralkar D, Kodra JT, Holst JJ, Jeppesen CB, Johnson MD, de Jong JC, Jorgensen AS, Kercher T, Kostrowicki J, Madsen P, Olesen PH, Petersen JS, Poulsen F, Sidelmann UG, Sturis J, Truesdale L, May J, Lau J. 2007. Small-molecule agonists for the glucagon-like peptide 1 receptor. Proceedings of the National Academy of Sciences 104:937–942. doi:10.1073/pnas.0605701104

Kuhre RE, Albrechtsen NJW, Deacon CF, Balk-Møller E, Rehfeld JF, Reimann F, Gribble FM, Holst JJ. 2016. Peptide production and secretion in GLUTag, NCI-H716, and STC-1 cells: a comparison to native L-cells. Journal of Molecular Endocrinology 56:201–211. doi:10.1530/JME-15-0293

Labouesse MA, Patriarchi T. 2021. A versatile GPCR toolkit to track in vivo neuromodulation: not a one-size-fits-all sensor. Neuropsychopharmacol 46:2043–2047. doi:10.1038/s41386-021-00982-y

Longwell CK, Hanna S, Hartrampf N, Sperberg RAP, Huang P-S, Pentelute BL, Cochran JR. 2021. Identification of N-Terminally Diversified GLP-1R Agonists Using Saturation Mutagenesis and Chemical Design. ACS Chem Biol 16:58–66. doi:10.1021/acschembio.0c00722

Manandhar B, Ahn J-M. 2015. Glucagon-like Peptide-1 (GLP-1) Analogs: Recent Advances, New Possibilities, and Therapeutic Implications. J Med Chem 58:1020–1037. doi:10.1021/jm500810s

Mijalis AJ, Thomas DA, Simon MD, Adamo A, Beaumont R, Jensen KF, Pentelute BL. 2017. A fully automated flow-based approach for accelerated peptide synthesis. Nat Chem Biol 13:464–466. doi:10.1038/nchembio.2318

Mirdita M, Schütze K, Moriwaki Y, Heo L, Ovchinnikov S, Steinegger M. 2022. ColabFold: making protein folding accessible to all. Nat Methods 1–4. doi:10.1038/s41592-022-01488-1

Nauck MA, Meier JJ. 2018. Incretin hormones: Their role in health and disease. Diabetes Obes Metab 20 Suppl 1:5–21. doi:10.1111/dom.13129

O’Neil PM, Birkenfeld AL, McGowan B, Mosenzon O, Pedersen SD, Wharton S, Carson CG, Jepsen CH, Kabisch M, Wilding JPH. 2018. Efficacy and safety of semaglutide compared with liraglutide and placebo for weight loss in patients with obesity: a randomised, double-blind, placebo and active controlled, dose-ranging, phase 2 trial. The Lancet 392:637–649. doi:10.1016/S0140-6736(18)31773-2

Patriarchi T, Cho JR, Merten K, Howe MW, Marley A, Xiong W-H, Folk RW, Broussard GJ, Liang R, Jang MJ, Zhong H, Dombeck D, von Zastrow M, Nimmerjahn A, Gradinaru V, Williams JT, Tian L. 2018. Ultrafast neuronal imaging of dopamine dynamics with designed genetically encoded sensors. Science 360:eaat4422. doi:10.1126/science.aat4422

Patriarchi T, Cho JR, Merten K, Marley A, Broussard GJ, Liang R, Williams J, Nimmerjahn A, von Zastrow M, Gradinaru V, Tian L. 2019. Imaging neuromodulators with high spatiotemporal resolution using genetically encoded indicators. Nature Protocols 14:3471–3505. doi:10.1038/s41596-019-0239-2

Rowlands J, Heng J, Newsholme P, Carlessi R. 2018. Pleiotropic Effects of GLP-1 and Analogs on Cell Signaling, Metabolism, and Function. Frontiers in Endocrinology 9.

Runge S, Wulff BS, Madsen K, Bräuner-Osborne H, Knudsen LB. 2003. Different domains of the glucagon and glucagon-like peptide-1 receptors provide the critical determinants of ligand selectivity. British Journal of Pharmacology 138:787–794. doi:10.1038/sj.bjp.0705120

Shah M, Vella A. 2014. Effects of GLP-1 on appetite and weight. Rev Endocr Metab Disord 15:181–187. doi:10.1007/s11154-014-9289-5

Stockklausner C, Klocker N. 2003. Surface expression of inward rectifier potassium channels is controlled by selective Golgi export. J Biol Chem 278:17000–17005. doi:10.1074/jbc.M212243200

Sun F, Zeng J, Jing M, Zhou J, Feng J, Owen SF, Luo Y, Li F, Wang H, Yamaguchi T, Yong Z, Gao Y, Peng W, Wang L, Zhang S, Du J, Lin D, Xu M, Kreitzer AC, Cui G, Li Y. 2018. A Genetically Encoded Fluorescent Sensor Enables Rapid and Specific Detection of Dopamine in Flies, Fish, and Mice. Cell 174:481–496.e19. doi:10.1016/j.cell.2018.06.042

Thompson A, Kanamarlapudi V. 2015. Distinct regions in the C-Terminus required for GLP- 1R cell surface expression, activity and internalisation. Mol Cell Endocrinol 413:66–77. doi:10.1016/j.mce.2015.06.012

Trapp S, Brierley DI. 2022. Brain GLP-1 and the regulation of food intake: GLP-1 action in the brain and its implications for GLP-1 receptor agonists in obesity treatment. British Journal of Pharmacology 179:557–570. doi:10.1111/bph.15638

Turton MD, O’Shea D, Gunn I, Beak SA, Edwards CM, Meeran K, Choi SJ, Taylor GM, Heath MM, Lambert PD, Wilding JP, Smith DM, Ghatei MA, Herbert J, Bloom SR. 1996. A role for glucagon-like peptide-1 in the central regulation of feeding. Nature 379:69–72. doi:10.1038/379069a0

Wan Q, Okashah N, Inoue A, Nehmé R, Carpenter B, Tate CG, Lambert NA. 2018. Mini G protein probes for active G protein–coupled receptors (GPCRs) in live cells. J Biol Chem 293:7466–7473. doi:10.1074/jbc.RA118.001975

Wang L, Wu C, Peng W, Zhou Z, Zeng J, Li X, Yang Y, Yu S, Zou Y, Huang M, Liu C, Chen Y, Li Yi, Ti P, Liu W, Gao Y, Zheng W, Zhong H, Gao S, Lu Z, Ren P-G, Ng HL, He J, Chen S, Xu M, Li Yulong, Chu J. 2022. A high-performance genetically encoded fluorescent indicator for in vivo cAMP imaging. Nat Commun 13:5363. doi:10.1038/s41467-022-32994-7

Wu F, Yang L, Hang K, Laursen M, Wu L, Han GW, Ren Q, Roed NK, Lin G, Hanson MA, Jiang H, Wang M-W, Reedtz-Runge S, Song G, Stevens RC. 2020. Full-length human GLP-1 receptor structure without orthosteric ligands. Nat Commun 11:1272. doi:10.1038/s41467-020-14934-5

Zhang X, Belousoff MJ, Zhao P, Kooistra AJ, Truong TT, Ang SY, Underwood CR, Egebjerg T, Šenel P, Stewart GD, Liang Y-L, Glukhova A, Venugopal H, Christopoulos A, Furness SGB, Miller LJ, Reedtz-Runge S, Langmead CJ, Gloriam DE, Danev R, Sexton PM, Wootten D. 2020. Differential GLP-1R Binding and Activation by Peptide and Non-peptide Agonists. Molecular Cell 80:485–500.e7. doi:10.1016/j.molcel.2020.09.020

## SI References

[1] N. Hartrampf, A. Saebi, M. Poskus, Z. P. Gates, A. J. Callahan, A. E. Cowfer, S. Hanna, S. Antilla, C. K. Schissel, A. J. Quartararo, X. Ye, A. J. Mijalis, M. D. Simon, A. Loas, S. Liu, C. Jessen, T. E. Nielsen, B.L. Pentelute, Science 2020, 368, 980–987.

[2] K. K.-S. Tso, K.-K. Leung, H.-W. Liu, K. K.-W. Lo, Chem. Commun. 2016, 52, 4557–4560.

[3] S. Kaneko, H. Nakayama, Y. Yoshino, D. Fushimi, K. Yamaguchi, Y. Horiike, J. Nakanishi, Phys. Chem. Chem. Phys. 2011, 13, 4051–4059.

